# Diploid genomes of the booklouse reveal evolutionary consequences of asexuality

**DOI:** 10.1101/2025.09.02.673870

**Authors:** Shiqian Feng, Andrea Pozzi, George Opit, Vaclav Stejskal, Zhihong Li

## Abstract

The “Meselson Effect“—a cornerstone of evolutionary theory predicting that asexual organisms accumulate mutations independently across haplotypes due to the absence of recombination—has long eluded robust empirical validation. Here, we address this gap using three high-quality diploid genome assemblies of the obligately parthenogenetic booklouse *Liposcelis bostrychophila*, a species characterized by apomictic meiosis and striking intraspecific variation. Combining Illumina, PacBio, and Hi-C sequencing, we generated three diploid genomes of three populations (Lb_1, Lb_2, Lb_3), revealing exceptionally high heterozygosity rates (1.67–3.92%) and minimal recombination—hallmarks of prolonged asexuality. Critically, in population Lb_2, intra-individual SNP divergence surpassed inter-population levels, directly demonstrating the Meselson Effect. Phylogenetic analyses of 7,029 genes revealed that while haplotypes within populations clustered as expected under sexual reproduction, nearly 15% of gene trees exhibited topologies consistent with asexual divergence—a signature of incipient Meselson-driven evolution. Comparative genomic analyses further uncovered a twofold expansion of repetitive elements in asexual lineages relative to their sexual relatives (*L. brunnea*) and widespread relaxation of purifying selection, particularly affecting transcription factor activities. We propose that relaxed selection on transcriptional regulators may act as a compensatory mechanism, buffering the functional consequences of escalating heterozygosity in non-recombining genomes. By integrating population genomics, phylogenomics, and selection analyses, this work not only provides the clearest evidence to date of the Meselson Effect but also establishes *L. bostrychophila* as a pivotal model for unraveling the genomic paradox of parthenogenesis.

**Significance Statement:** The Meselson Effect, a theoretical prediction that asexual organisms should accumulate mutations independently in their two haplotypes, has been difficult to confirm empirically due to limited genomic data and contradictory findings in prior studies. Here, we provide strong evidence for the Meselson Effect in the parthenogenetic booklouse *Liposcelis bostrychophila*, using high-quality diploid genomes from three populations. We show that one population exhibits higher genetic divergence within individuals than between populations, a key signature of the Meselson Effect. Additionally, we found a possible novel compensatory mechanism for maintaining genomic stability in the face of increased heterozygosity. These findings advance our understanding of the genomic impacts under asexuality and establish *L. bostrychophila* as a model for studying the evolutionary consequences of parthenogenesis.

## Introduction

A key prediction for the evolution of asexual organisms, where recombination and segregation are absent, is that the two haplotypes in a diploid clonal lineage will independently accumulate mutations (1). This prediction is known as the “Meselson Effect” (Fig. 1). The existence of this effect is yet to be proven, and testing of the Meselson Effect has been focused on only two clades: oribatid mites and the stick insect genus *Timema* (2). Researchers observed greater mutation accumulation within individuals than between populations in oribatid mites, signals consistent with the Meselson effect. However, in *Timema*, the opposite was found: asexuality led to reduced heterozygosity. It is unclear why these studies found contradictory results, but technical limitations might have played a role. Indeed, the absence of well-assembled diploid genomes remains a major limitation for accurate haplotypic-level variation calling, as prior studies were limited to contig-level genomes or coding gene analyses.

**Fig. 1.**
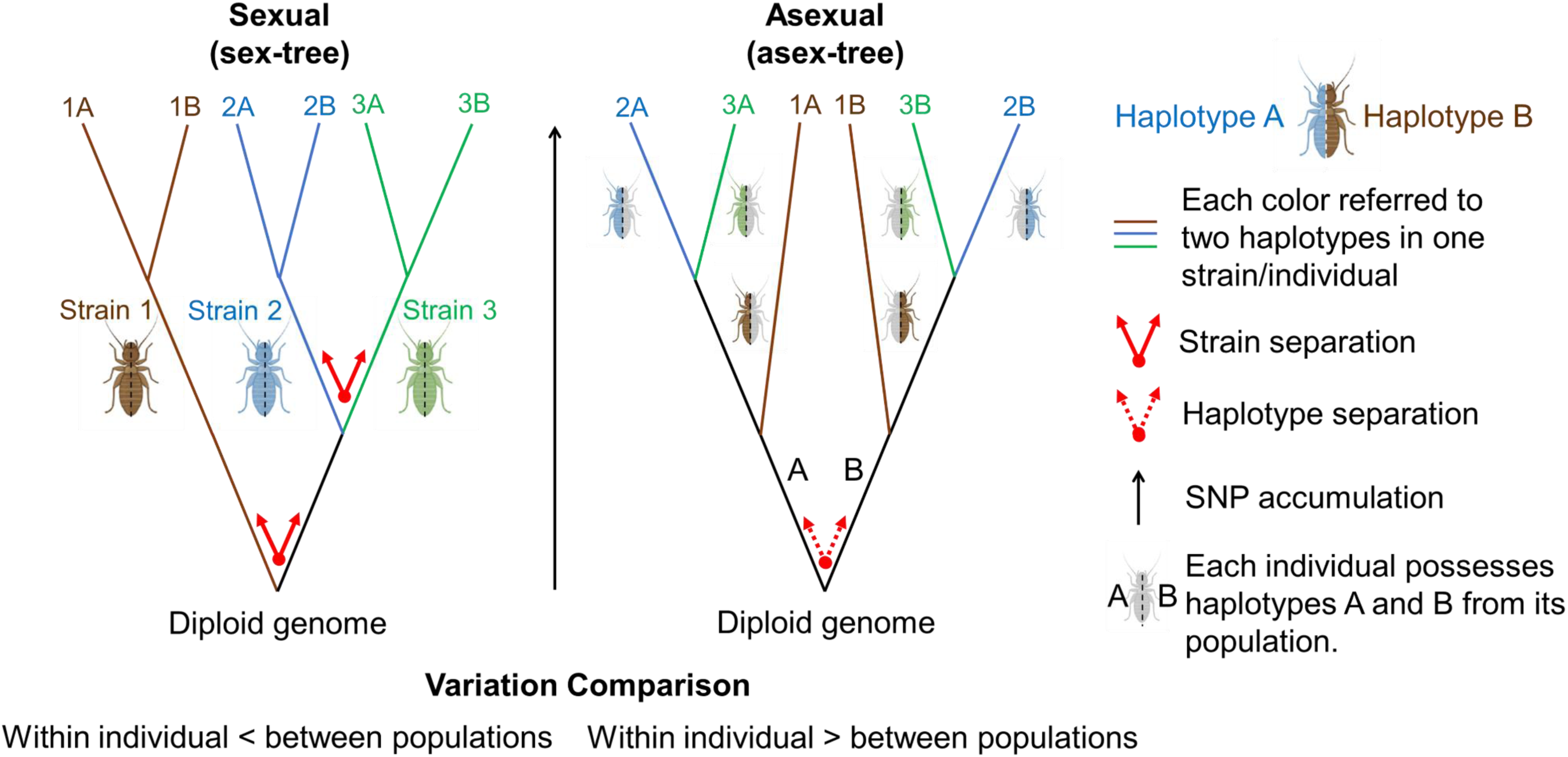
Nuclear haplotype trees expected under (long-term) obligate asexual and sexual reproduction. In diploid, functionally clonal organisms, homologous chromosomes (A and B) accumulate mutations independently of each other both of which will ultimately possess stable and distinguishable mutations across the chromosome or specific regions. These intra-individual/intra-genomic divergences can even exceed divergence from different populations (inter-population/inter-genomic level), termed “Meselson Effect”. In asexuals, the haplotype tree (asex-tree) fully separates homologous haplotypes at its deepest split (dashed red arrow), reflected by a heightened heterozygosity. In sexuals, the haplotype tree (sex-tree) separates the populations with haplotypes from the same population form their own clade (full red arrow).

It is, therefore, necessary to identify more asexual species with extensive intraspecific variation, that could be tested for the presence of the Meselson Effect. Previous research suggests that species belonging to the clade *Liposcelis* (a type of booklice) could be ideal candidates. Booklice, small insects commonly found in many locations, from bird nests to grain facilities, have been at the center of agricultural and evolutionary research. Indeed, several species in the clade *Liposcelis* have sex distortion in offsprings (3, 4) and show significant intraspecific sequence variation (5). In particular, a parthenogenetic species, *Liposcelis bostrychophila*, conducting apomictic meiosis (6), has shown structural rearrangements and substantial sequence variation (48.7∼87.4% identity of mitochondrial genes across different populations) in its mitochondrial genome (7). Given that both mitochondrial and nuclear genomes are maternally inherited in asexual species, theory predicts that significant sequence differences might exist between the nuclear genomes of three *L. bostrychophila* populations (Lb_1, Lb_2, and Lb_3), and we tested this prediction through several genomic analyses on their newly sequenced diploid genomes.

## Results

### De Novo Genomes

To retrieve the complete diploid genomes of Lb_1, Lb_2, and Lb_3, we combined short (Illumina), long reads (PacBio), and Hi-C technology (Table S1). This approach yielded sixteen chromosomes successfully assembled for each population (2n=16), with distinct Hi-C signals between homologous chromosomes (Fig. 2a), >90% BUSCO completeness (Fig. S1, Table S2), ∼500 MB in sizes (Table S3), and resolved assembly SNPs (Fig. S2), suggesting a substantial improvement on the previously published booklouse genomes (8, 9) (see methods for details). We tested the validity of the assembly by aligning the DNA data from a single individual to the diploid genome assembly. This test found a uniform high coverage across the genomes, indicating the two haplotypes assembled coexisted in one individual across all three populations. The three populations were indicated with numbers (1, 2, 3), and the two haplotypes were labeled with letters (A and B). These assemblies suggest low levels of recombination between homologous chromosomes, as expected of species with prolonged asexuality (10). Similarly, the synteny analysis of the three diploid genomes of *L. bostrychophila* and a closely related species, *L. brunnea,* confirmed a low recombination rate (Fig. 2b). The analysis revealed strong synteny between the two species, and no inter-chromosomal rearrangements were detected among the six haplotypes of *L. bostrychophila*, indicating consistent genome architecture both within an individual and between populations, supporting the robustness of our assemblies. Nonetheless, we detected high heterozygosity rates (Fig. S3, 3.92%, 3.19%, and 1.67%), much higher even when compared to species studied for the Meselson effect (eg. media 1% for mites) (11). High heterozygosity, representing substantial genetic variation within an individual, is common in parthenogenetic plants due to polyploidy but is unusual in animals and insects (12). The Meselson effect can generate high heterozygosity through the independent accumulation of mutations not purged due to a low recombination rate, thus these genomes are likely affected by the Meselson effect.

**Fig. 2.**
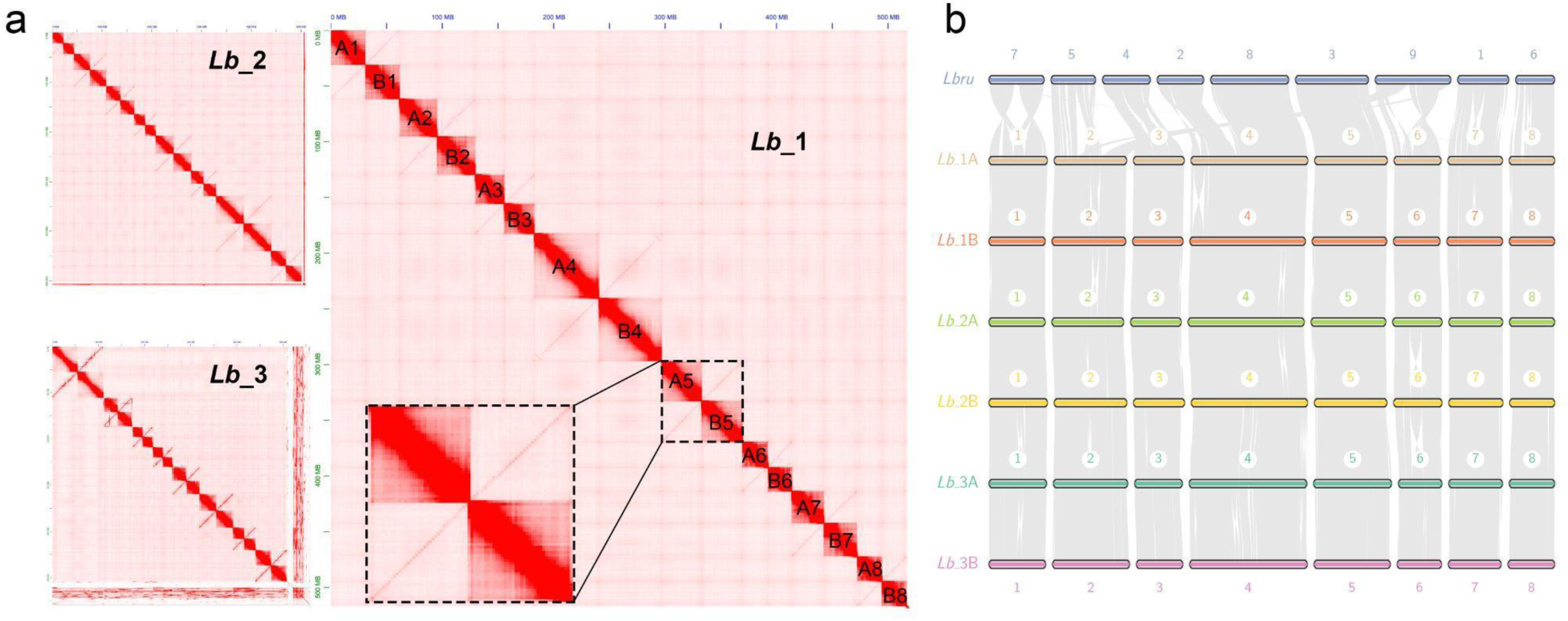
HiC signals and syntenic analysis of booklouse diploid genomes. **a,** HiC signals of the three diploid genomes. **b,** Synteny detection among three diploid genomes of *Liposcelis bostrychophila* and haplotypic genome of *L. brunnea*.

### Heterozygosity

We performed a heterozygosity analysis, testing if the variation within individuals is higher than between populations, a typical phenomenon of the Meselson effect (Fig. 3). To perform this test, we used the diploid genomes generated before, where each pair of chromosomes belonged to a different haplotype (either A or B). The comparisons within individuals are named “Lb_1A vs Lb_1B”, “Lb_2A vs Lb_2B” and “Lb_3A vs Lb_3B”. Similarly, we tested the differences between populations using the following nomenclature: “Lb_1A vs Lb_2X or Lb_3X”, “Lb_2A vs Lb_1X or Lb_3X” and “Lb_3A vs Lb_1X or Lb_2X” where “X” represents either haplotype A or B.

**Fig. 3.**
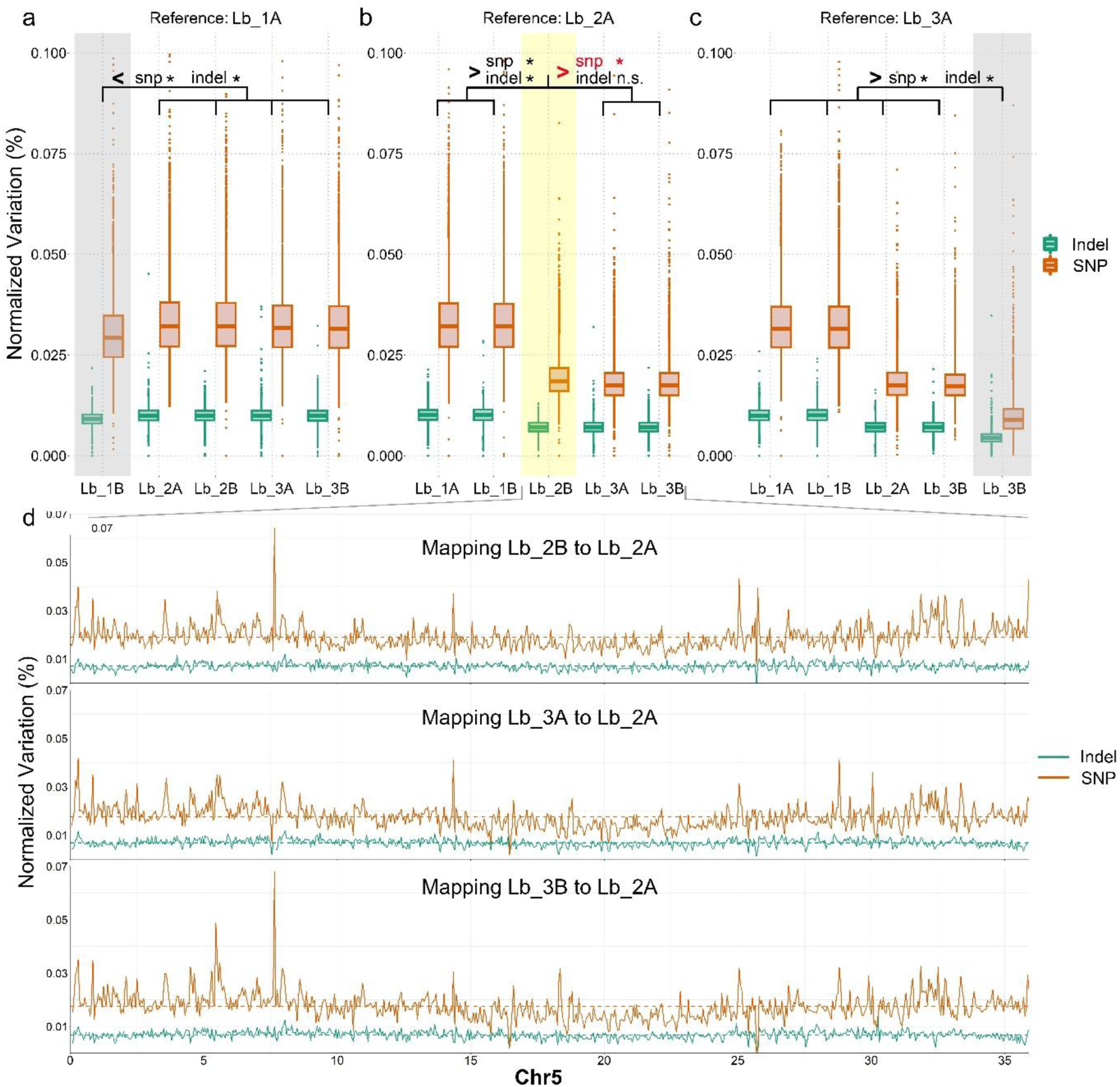
SNP and indel detection between *L. bostrychophila* haplotypes. **a,** Lb_1A-based variant calling. * means significantly lower variants between Lb_1A and Lb_1B than in other analyses. **b,** Lb_2A-based variant calling. * means intra-individual variation (between Lb_2A and Lb_2B) is significantly lower than Lb_1X but higher than Lb_3X. **c,** Lb_3A-based variant calling. * means significantly lower variants between Lb_3A and Lb_3B than other comparisons. **d,** a case showing higher intra-individual (between Lb_2A and Lb_2B) variation level than inter-population level (between Lb_2A and Lb_3X) in chromosome 5.

When comparing Lb_1A to the other haplotypes, we found higher SNP and indel ratios between populations (vs Lb_2X or Lb_3X) than within individuals (eg. Lb_1A vs Lb_1B, Fig. 3a, S4-S11), indicating a similar evolutionary history of the two haplotypes in Lb_1. We then compared Lb_2A to Lb_1X, finding ∼2x more SNPs (Fig. 3b, S12-S19) than to Lb_3X, suggesting long independent evolution for Lb_1X. However, the most important finding for Lb_2A is that it is more divergent from its “homologous chromosome” Lb_2B than to the other population (eg. a higher number of SNPs than to Lb_3X, no differences in indels, Fig. 3B). This pattern exemplifies the Meselson effect, where haplotypes within the same diploid genome (Lb_2A vs Lb_2B) have a higher degree of divergence than between populations (Lb_2A vs Lb_3X, Fig.3d). The last comparison, focused on Lb_3A, with low differences within individuals and high between populations (Fig. 3c, S20-S27). These results indicated that while the pattern observed on Lb_2 strongly supports the Meselson effect, this effect is unstable, as the other populations, Lb_1 and Lb_3, do not show signs of high variation within individuals. Overall, these results align well with previous results, finding that the Meselson effect is not always present, but can only be detected in certain populations.

### Phylogeny

A typical indication of haplotype divergence in asexual organisms is the complete segregation of haplotypes A and B at the most basal split of a haplotype tree. In contrast, in sexual organisms, haplotype divergence usually aligns with the patterns of population divergence (Fig. 1). To test this prediction, we reconstructed the phylogeny of the six haplotypes using whole chromosomes and individual genes. Across all six haplotypes, we identified between 534 and 1,736 ortholog genes per chromosome, for a total of 7,029 genes (Table S4). As it is challenging to determine which chromosomes constitute a complete haplotype, we instead used the concatenation of genes from each pair of chromosomes, generating eight phylogenetic trees (Fig. 4a). All eight trees possessed a tree topology consistent with what is expected for a sexual species (Fig. 1). All nodes have bootstrap values of 100 and this robust phylogenetic signal indicates that Lb_1 diverged earlier than Lb_2 and Lb_3, while the two haplotypes from each strain formed distinct clades, supporting a typical sexual reproduction (see Fig.1). Along with the chromosome trees, we generated the independent maximum-likelihood trees for 7,029 individual genes and coalesced them into an ASTRAL tree (Fig. 4a). The ASTRAL tree shows the same topology to the chromosomal trees, confirming that the overall phylogenetic signals resemble what observed in sexually reproducing species. Theoretically, the chromosomal/ASTRAL trees indicated an overall common phylogenetic signal from their last divergence whereby the Meselson Effect related evolutionary pathway cannot produce an identical signal, making the sex-inherited signals dominant in sequence analyses. To test this, we re-checked the topologies of the 7,029 individual gene trees and surprisingly found them classified into 824 tree topologies (Figure S28, Table S5) with the six main topologies possessing frequencies only from 1.75% to 3.83% (Fig. 4b). The many topologies indicate that the group of genes could show a different phylogenetic signal, thus we performed a more detailed analysis of these genes to identify where any of them is under Meselson Effect.

**Fig. 4.**
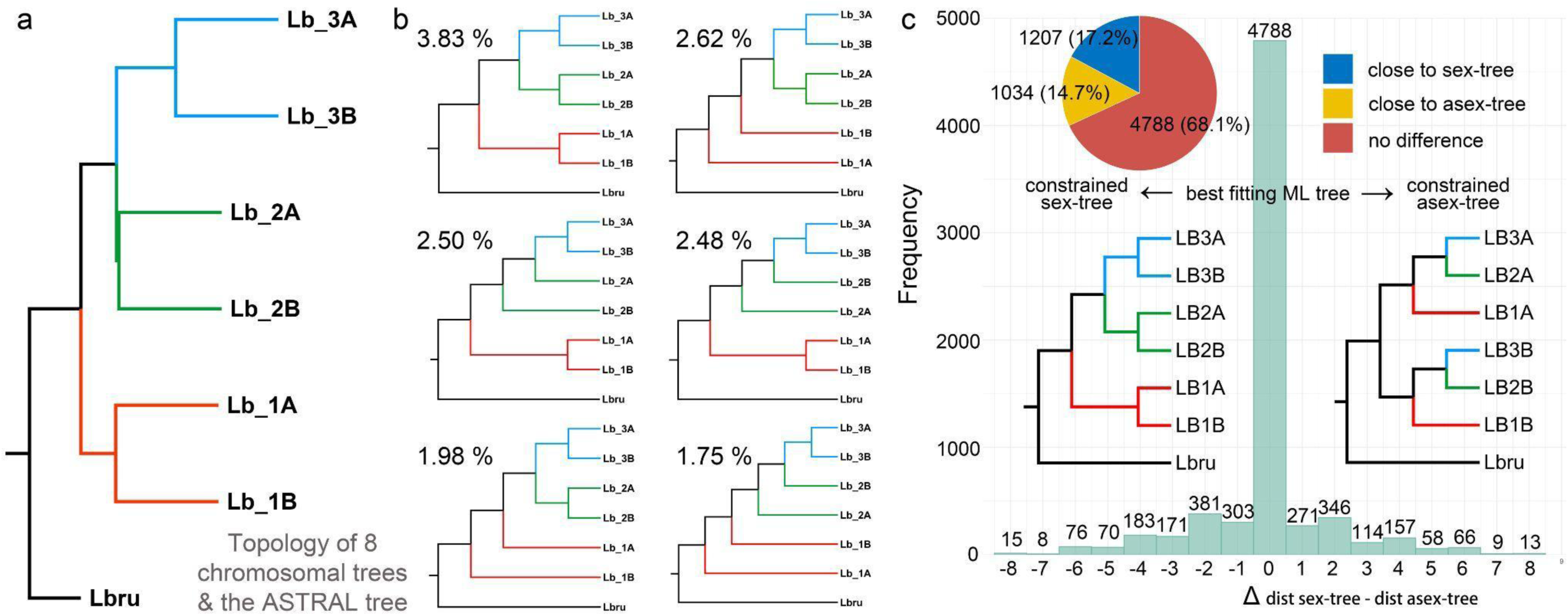
Phylogenetic analyses. **a,** phylogenetic inference using IQTREE based on concatenated CDS sequences for each of 8 chromosomes and the ASTRAL tree based on 7,029 ortholog genes, resulting in the same topology. **b,** top 6 phylogenetic trees of 7,029 individual gene trees and the frequencies. **c,** Frequency distribution of 7,029 trees’ distance score comparisons (Δdist. sex-tree − dist. asex-tree), showing the Delta of the best-fitting ML tree to the sex-tree and asex-tree. A positive value indicates a topology close to asex-tree while individual genes with negative values are close to the sex-tree. Zero indicated equal distances of individual genes to either tree.

We next detected if the 7,029 individual gene trees are similar to the “sex-tree”, splitting the clades (Lb_1, Lb_2, Lb_3) at the basal or the “asex-tree”, separating the haplotypes (A, B) at the basal (Fig. 1, Fig. 4c). We found 4788 genes (68.1%) showed no preference to both topologies, which suggested most genes are under an intermediate state. For the rest of genes, there are more showing a close relationship to sex-tree (1207 genes, 17.2%) than to asex-tree (1034 genes, 14.7%). Our finding suggested an evolutionary process of reproduction mode and a slight preference for sex-tree against asex-tree (173 genes more), which could technically explain why the chromosomal and ASTRAL trees showed the same topology to sex-tree.

Similar to what was observed in other analyses, the analysis of sequence similarity (p-distance) across species showed that haplotypes within each population were more similar, followed by population and species levels (Fig. S29). Based on the previous analyses, we have found traces of the Meselson effect, as while the overall phylogenetic signal shows a pattern consistent with sexual reproduction, there are almost as many genes showing a topology consistent with parthenogenetic reproduction.

### Repeat expansion

Along with the Meselson effect, another effect of parthenogenesis could be an increase in the number of repeat regions (RRs), such as transposable elements (13, 14). Thus, we analyzed the RRs across the diploid genomes available for booklice, using the three asexual populations of *L. bostrychophila,* and the sexual *L. brunnea* (Fig. 5ab, Table S6), comparing the abundance of RRs across chromosomes. We found about twofold more RRs in the asexual species than in the sexual species. However, we found only small differences (< 2%) in the RR quantity across three asexual populations (Fig. 5c). Our findings also supported the theory that parthenogenesis species had a reduced RR due to the dysfunctional recombination process (2, 15).

**Fig. 5.**
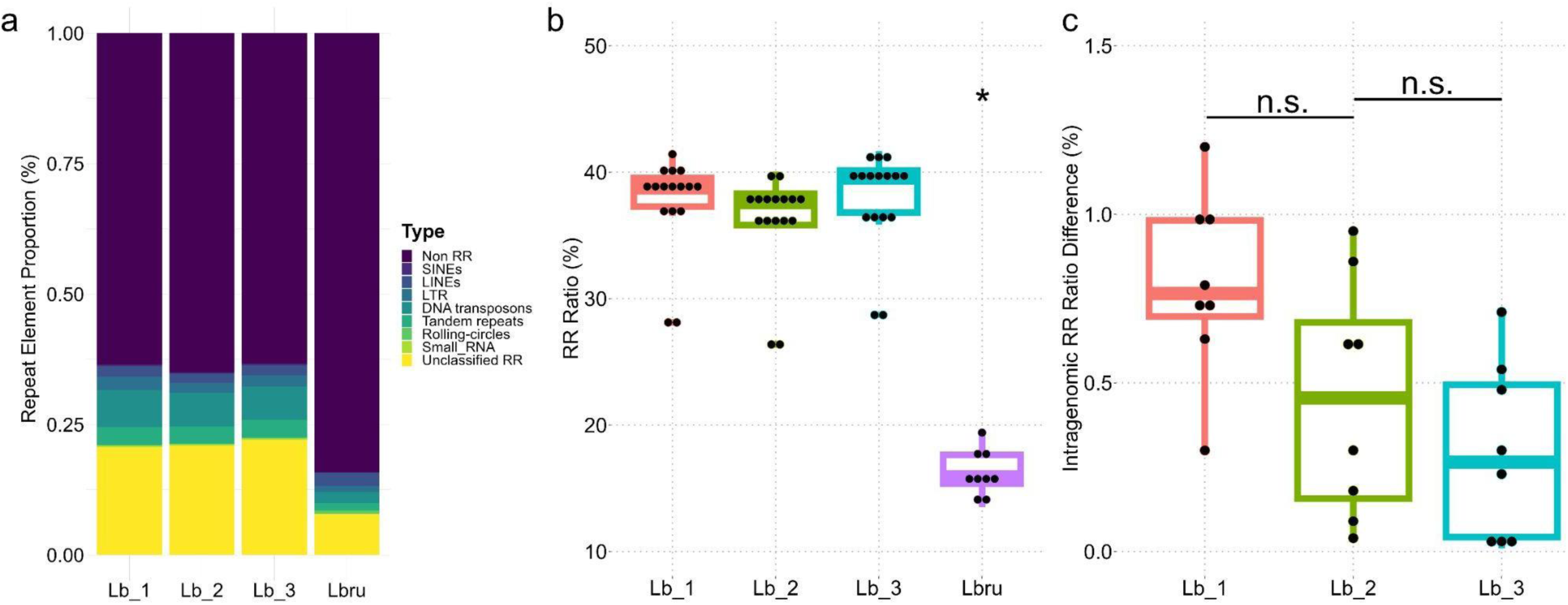
Repeat Region (RR) analyses. **a,** repeat element ratios in booklice diploid genomes (Lb_1, Lb_2, Lb_3) and haplotypic genome (Lbru). **b,** RR ratios of all chromosomes in booklice genomes. * means significantly lower RR ratio in Lbru. **c,** RR ratio differences between 8 pairs of homologous chromosomes in each strain of *Liposcelis bostrychophila*. * means significant lower RR difference in Lb_3 than Lb_1; n.s. means non-significant differences between Lb_1/3 and Lb_2.

### Selection Pressure

Obligate parthenogenesis represents an extreme case of sex loss, often resulting in the defunctionalization of sex-related genes, causing relaxed selective and eventual pseudogenization(16). Through the high-quality assembly generated, we can avoid paralog effects that might otherwise confound our analyses, and detect eventual functional divergence between the alleles of the two haplotypes 17(17, 18). We calculated the dN/dS ratios between the two haplotypes in each of the three populations to evaluate how selection pressure changed across alleles (Fig. 6a). We observed a positive correlation between chromosome length and the number of genes with dN/dS >0.6, indicating a uniform distribution of relaxed selection pressures, with no specific region enriched for this effect (Fig. S30). We classified genes as under relaxed pressure when the dN/dS is higher than 0.6, identifying 1,384 genes in Lb_1, 1,951 in Lb_2, and 2,865 in Lb_3 (Fig. 6b). We then performed a GO term enrichment analysis, finding 69, 17, and 3 enriched GO terms in the three populations, respectively (Fig. 6c, Table S7-S10). Among these, transcription factor-related functions were enriched across all three populations: GO:0003705, transcription factor activity with RNA polymerase II distal enhancer sequence-specific binding in Lb_1, Lb_2; and GO:0001077, transcriptional activator activity with RNA polymerase II core promoter proximal region binding in Lb_3. Reduced selection pressure on transcriptional factors has broad consequences, influencing different aspects of cell regulation, and far exceeding our expectation of an effect only on sex-related genes (Table S11). This result might be evidence of a general compensatory mechanism for the Meselson effect. Lower selection pressure on transcriptional factors increases the number of mutations generated, potentially increasing the chance of still interacting with the increasingly different alleles present within the same cells.

**Fig. 6.**
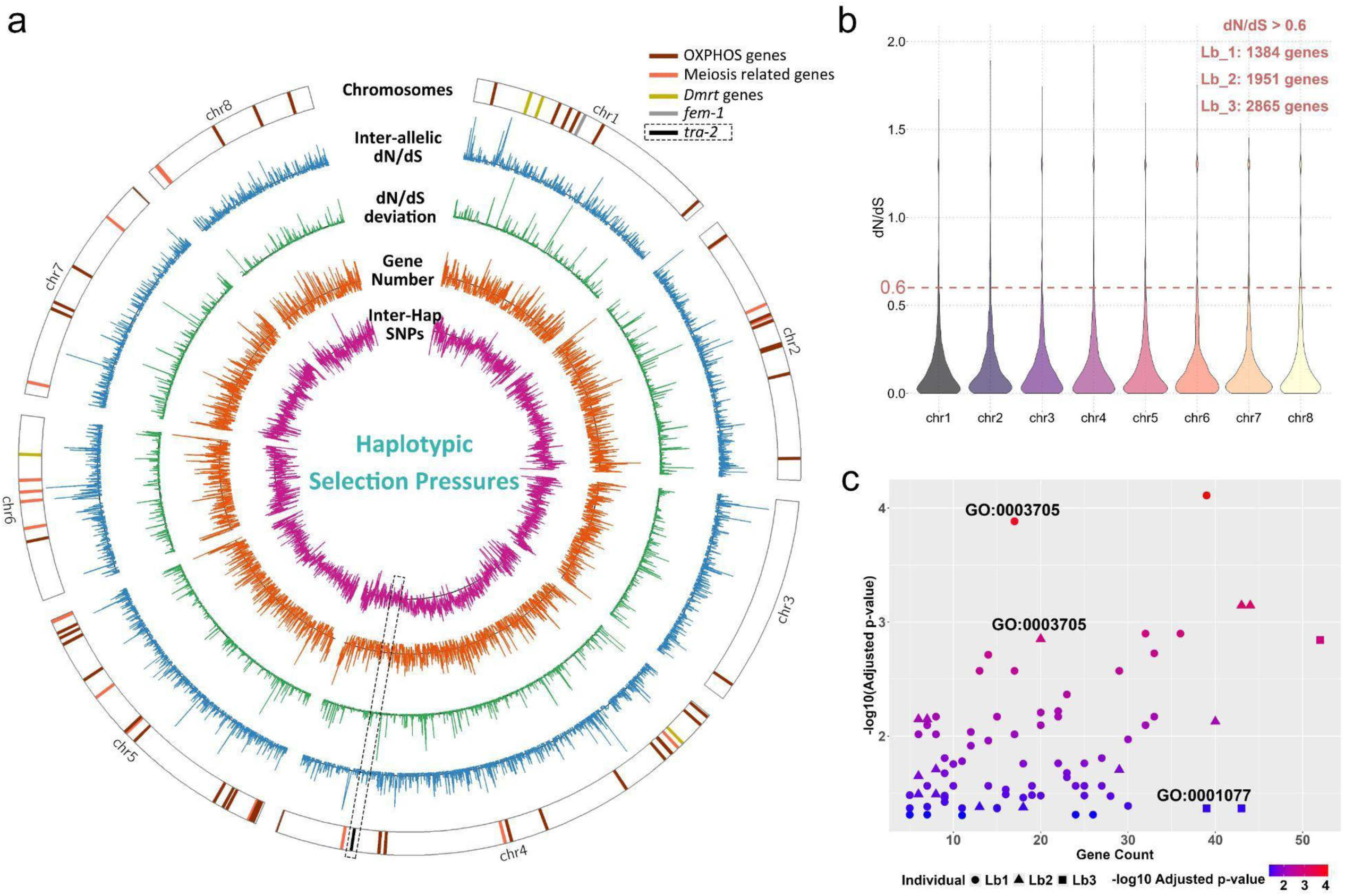
Allelic dN/dS calculation and function enrichment. **a,** Purple line representing inter-haplotypic SNPs, orange line representing gene numbers, green line representing dN/dS deviation |dN/dS(A vs *L. brunnea*) – dN/dS(B vs *L. brunnea*)|blue line representing inter-allelic dN/dS. OXPHOS genes, meiosis-related genes, *Dmrt* genes, *fem-1* gene, and *tra-2* gene were labeled in the chromosomes with different colors. The sliding window is set at 50kb. **b,** Allelic dN/dS were calculated and summarized for each chromosome of *Liposcelis bostrychophila*. **c,** GO terms enriched with dN/dS > 0.6 genes in three strains of *L. bostrychophila*.

## Discussion

In this study, we have found several supporting evidence for the Meselson effect in *L. bostrychophila*. One of the key predictions of this effect is the increased diversity within individuals than between populations, which we have found in Lb_2. The higher heterozygosity of this population perfectly matches this prediction, however, the effect was not strong enough to distort the overall phylogenetic signal, suggesting that the diverging of haplotypes might still be at its initial stages. Another evidence of the Meselson effect is the increase in RRs in *L. bostrychophila* compared to its closest sexually reproducing relative, although this relies only on one pair of comparisons. Furthermore, we have documented a potentially novel effect of parthenogenesis.

We detected relaxed selection on reproductive genes, as predicted by theory, and, unexpectedly, on transcription factors. Many transcriptional factors bind DNA sequences directly to facilitate transcription, making mutations on these DNA sequences likely to affect the binding. In a diploid genome where homologous chromosomes have high heterozygosity but low recombination rate, transcriptional factors under weaker purifying selection have more chances to develop mutations with higher affinity to both homologous chromosomes. Therefore, the relaxed selection of transcriptional factors could be a compensatory mechanism that mitigates the consequences of heightened heterozygosity. This compensatory mechanism might work similarly to what is observed in mitonuclear interactions, where nuclear proteins interacting with the mitochondria evolve faster to match the higher mutation rate in the mitochondria (19). These findings position *L. bostrychophila* as a model for studying the genomic impacts of parthenogenesis and the Meselson Effect.

## Materials and Methods

### Sample Preparation, Nucleic Acid Extraction, and Sequencing

Three strains of *L.bostrychophila* were collected in Beijing, China (Lb_1), Prague, Czech Republic (Lb_2) and Kansas State, United States (Lb_3). Morphological and molecular identification was conducted in our previous published works (5, 7). The rearing environment was maintained at 25°C, 75% relative humidity, and complete darkness, with a diet ratio of whole wheat flour: yeast: high-calcium milk powder = 10:1:1.

The genomic sequencing of all three strains was divided into five parts: 1) PacBio sequencing of genomic DNA inferred from a mixed sample from 500 individuals, to sequence on one PacBio CLR cell for genome assembly; 2) Illumina sequencing of genomic DNA, using a mixed sample from 40 individuals to sequence 20 Gb for genome survey and polishing; 3) Illumina sequencing of single individual genomic DNA, yielding 6 Gb for assembly result verification; 4) RNA-seq using Illumina platform, with a mixed sample from 40 individuals to sequence 6 Gb for genome annotation; 5) Hi-C sequencing, using 500 female individuals processed according to Hi-C protocols, retrieving 30 Gb data to help locate scaffolds into chromosomes. All DNA extractions were performed using the Promega Genomic DNA Purification Kit (A1125), and RNA extractions were conducted using the Tiangen RNA Extraction Kit (DP431). All Pacbio and Illumina sequencing were completed by Berry Genomics, utilizing the NovaSeq 6000 platform with an insert fragment size of 350 bp and paired-end 150 bp mode.

### Genome Survey, Assembly and Evaluation

We first conducted genome surveys using over 80× Illumina sequencing data (Fig. S3, Table S1). The results indicated similar genome sizes (245.4 Mb, 240.3 Mb, and 261.2 Mb). Next, we performed PacBio sequencing (PBS) and Hi-C technology, yielding over 100× PBS and 100× Hi-C data for each population. For all Illumina sequencing data, quality control was performed using fastp v0.20.1 (20), filtering low-quality sequencing data. Genome sizes and heterozygosities of the three populations were assessed using GenomeScope v2.0 (21) leveraging illumina sequencing data. Genome assembly was performed using the PacBio CLR sequencing data with Canu v2.1.1 (22), setting parameters as follows: minReadLength=2000, minOverlapLength=500, corOutCoverage=120, corMinCoverage=2, correctedErrorRate=0.035. The setting of “correctedErrorRate” was aimed at splitting the two haplotypes contained in the genome as much as possible to obtain more complete genome contigs. For the genome draft obtained from the Canu assembly, the first round of correction was performed using pbmm2 v1.40 and gcpp v1.9.0 (https://www.pacb.com/support/software-downloads/) using PacBio sequencing data. The second round of correction was conducted using Pilon v1.23 (23) based on the illumina sequencing data of genomic DNA, resulting in three corrected diploid genomes. For the corrected genome, scaffold relocation was conducted using 3D-DNA v180114 and Juicebox v1.11.08 software with Hi-C data to obtain diploid genomes at the chromosome level (24). BUSCO v5.1.3 was used for assembly evaluation at each assembly step with the insecta_odb10 database.

To validate the assembly quality of the three diploid genomes, we conducted align-back tests, synteny analysis, and completeness evaluations. For example, total DNA from a single individual of Lb_1 population was extracted for Illumina sequencing and aligned to both the haploid (Lb_1A) and diploid (Lb_1AB) genomes. When aligned to one haplotype, the alignment rate was ∼80%, with chimeric sites showing heterozygosity rates around 50%, indicating stable inter-haplotypic SNPs (Fig. S2). In contrast, alignment to the diploid genome (Lb_1AB) resulted in a significant increase in correct alignment rates to 96.77%, resolving nearly all discrepancies at variant sites. Similar results were obtained for Lb_2 and Lb_3.

### Genome Annotation

To obtain chromosome-level diploid genomes, we constructed *de novo* repeat sequence libraries and masked repeat sequences using RepeatModeler v2.0.1 (25) and RepeatMasker v4.1.0 (26). We then annotated the genome using the Maker v3.01.03 (27) pipeline, utilizing protein sequence data from the human louse (*Pediculus humanus*), western flower thrips (*Frankliniella occidentalis*), red flour beetle (*Tribolium castaneum*), fruit fly (*Drosophila melanogaster*), and honeybee (*Apis mellifera*). Using Trinity v2.11.0 (28), we conducted *de novo* assemblies of the transcriptomes for the three populations, obtaining relatively complete transcript information; homologous sequence searches and polishing were conducted using BLAST v2.11.0 (29) and Exonerate v2.58.3 (30); two rounds of gene model training were performed using SNAP v2006-07-28 (31) and Augustus v3.3.3 (32), ultimately yielding the annotation results of three diploid genomes. Functional annotations of the obtained protein sequences were performed using eggNOG-mapper v2 (33) against nr, GO term, KEGG pathway, COG, and InterPro databases.

### Validation of Genome Assembly Results

The assembled diploid genomes of *L. bostrychophila* are 517.6 Mb (Lb_1), 502.0 Mb (Lb_2), and 509.4 Mb (Lb_3), approximately twice the estimated haplotypic sizes (Fig. S3). The homologous chromosomes within each population are in similar sizes, with size variation ranging from 0 to 6.7%. After annotation, we identified 34,469 34,989 and 34,630 genes in the three diploid genomes, reflecting approximately double the gene count typical of a haplotypic insect genome (8, 34) (Table S2). The three diploid genomes also exhibited high BUSCO completeness (> 97%), and contained similar repeat element profiles (Table S6). Notably, while we designated homologous chromosomes with “A” and “B,” these do not represent distinct haplotypes within the Lb_1A chromosome set. Instead, all 16 chromosomes in each population individual should be considered individually due to the absence of homologous recombination in asexual reproduction.

Assembly validation of three diploid genomes were carried out from three aspects:

(1) To ensure that the diploid genome is present in the same booklouse individual, Illumina sequencing data of genomic DNA from one individual was aligned to the haploid or diploid genome using BWA v0.7.17 (35). By comparing and analyzing whether the diploid genome can accurately resolve heterozygous sites and improve the alignment rate of sequencing data, we will confirm the reliability of the assembled diploid genomes.
(2) Syntenic analysis of six haplotypes from three populations of *L. bostrychophila* and the merged haplotypic genome of *L. brunnea* was conducted using the JCVI toolkit (36) to analyze the haplotypic synteny within and between *L. bostrychophila* strain, as well as between the two booklice species. A good syntenic result could indicate a good genome assembly.
(3) BUSCO completeness assessment of the six haplotypes from three diploid genomes of *L. bostrychophila* was conducted, where good BUSCO assessment results for each haplotype will indicate good gene integrity.

### Sequence Feature Analysis

After repeat masking, we conducted sequence alignments between any two of the six haplotypes using “nucmer” function from the Mummer v3.1 software package(37), with minimum continuous similarity length setting to 20 bp (-l 20) and minimum cluster size of 65 bp (-c 65), followed by “delta-filter” function with the minimum alignment identity of 50% (-i 50) and retaining only one-to-one alignment results (−1). The alignments were conducted with three haplotypes (Lb_1A, Lb_2A and Lb_3A) as the references, respectively, and the other five haplotypes were aligned to each reference sequence in each round. We used the “makewindows” tool from BEDTools v2.30.0(38) to create sliding windows of 50 kb length for each reference haplotype, with a step size of 50 kb. Based on the “nucmer” results, we counted the number of SNPs and indels in each window using the “intersect” tool from BEDTools. SNPs and indels inferred were treated as differences between the two haplotypes, and used for testing if intra-genomic variation exceeded inter-genomic variation, termed “Meselson Effect”.

### Identification and Analysis of homologous genes

The identification of syntenic homologous genes relies on the one-to-one alignment results from synteny analysis, which includes 3 cases: 1) Analyzing the syntenic relationships among the six haplotypes from the three populations of *L. bostrychophila* and the genome of *L. brunnea*, to confirm the overall homologous gene relationships of existing booklice genomes; 2) Analyzing the syntenic relationships between the two haplotypes within each individual of *L. bostrychophila* to obtain pairwise homologous genes as much as possible; 3) Analyzing the syntenic homologous gene pairs between the three populations of *L. bostrychophila* and *L. brunnea* to examine the genetic distances and selection pressures within each population. The identification of syntenic homologous genes/synteny modules was performed using the jcvi.compara.catalog function from the JCVI toolkit, with parameters set to: --no_strip_names –full --dbtype=prot, inputting the protein sequences (.pep) and annotation files (.gff) for each haplotype. The resulting .anchors.new file contains the retained high-quality co-linearity analysis results for downstream analysis.

### Phylogenetic Analyses

Ortholog genes across six haplotypes of *L. bostrychophila* and the genome of *L. brunnea* were inferred from syntenic analysis. Each protein coding gene was aligned using Muscle v3.8.1551(39) with default settings and Gblocks 0.91b(40) was leveraged to retrieve those conserved regions with the parameter “-t c -b1 6 -b2 9 -b3 8 -b4 10 -b5 n”. After sequence concatenation for each chromosome, IQTREE v2.2.3(41) was used for phylogenetic reconstruction with extended model selection followed by tree inference (-m MFG) and bootstrap setting to 1,000. In total 8 phylogenetic trees were inferred for each chromosome with *L. brunnea* as the outgroup.

In total 7,029 pairwised homologous genes were identified across 6 haplotypes of *L. bostrychophila* and the genome of *L. brunnea*. These genes were used in phylogeny inference one-by-one, following the IQTREE settings above to construct 7,029 gene trees. These trees were summarized using SumTrees in DendroPy 5 toolkit(42) and got topology types and counts. All 7,029 trees were subsequently fed into ASTRAL(43) to produce a coalescent-based summary tree. These trees were used for gene diversification analyses.

### Topology Testing

To enable testing whether 7,029 gene trees are better explained by a topology separating the haplotypes (as expected under Meselson Effect) compared to a topology separating populations (expected under sexual reproduction) or vice versa, we conducted constrained tree searches following the methods described in Brandt et al. (2021). Specifically, two constrained ML trees, with a fixed haplotype-separating topology (asex-tree, Fig.1), the other with a fixed population-separating topology (sex-tree, Fig.1), were calculated for each gene using IQTREE with 1,000 bootstrap replicates and model-testing included. The asex-tree separated the haplotypes A and B at its base, while sex-tree split the population at its base and both haplotype clustered together in the tips. Next, the distance between the two resulting trees and an unrestricted best-fitting ML tree was estimated according to Penny and Hendy’s method (44) with the dist.topo function implemented in the R package ape. To enable the comparison between distances of these genes, the topological distances of the best-fitting ML tree to sex-tree and asex-tree were combined as Δ_dist. sex-tree - dist. asex-tree_ value for each gene: < 0 values indicated the unconstrained best-fitting ML tree was closer to sex-tree, while > 0 values suggested a closer relationship of the ML tree to asex-tree.

### Selection Pressure and Functional Enrichment Analysis

For each strain, the protein sequences of all pairs of syntenic homologous genes between the two haplotypes were aligned using Muscle v3.8.1551, and DNA alignment files were obtained using TranslatorX v1.1(45). Using PAML v4.9 and based on Model0, we calculated the dN/dS for each pair of alleles within each population. The dN/dS values were filtered with a criterion of dN/dS > 0.6 and these genes were usually recognized as under relaxed selection pressure. Additionally, we filtered for alleles that were completely identical within the same population, as these regions are likely to contain conserved, important functions or be subject to active recombination. These genes were subsequently used for functional enrichment leveraging clusterProfiler v4.2.1 package in R v4.1.2(46) to identify pathways undergoing rapid functional changes. For each strains of *L. bostrychophila*, we calculated dN/dS value between syntenic homologous genes of haplotypes A and B. Nonetheless, dN/dS values were calculated between pairwise genes from each haplotype and *L. brunnea*, to derive the absolute value S of the dN/dS difference within each individual (|dN/dSAvsLbru-dN/dSBvsLbru|), comparing the magnitude of functional deviations between the two gene sets. A larger S value indicates a greater potential functional difference in the respective genes on the two haplotypes. Analyzing S can validate whether one of the alleles in parthenogenetic species remains biologically stable while the other exhibits significant functional variation, providing more functional predictions of related genes.

## Data Availability

The Pacbio genome sequencing, RNA Illumina sequencing, DNA Illumina sequencing and Hi-C sequencing raw data have been updated to NCBI as a BioProject PRJNA1207765. Genome assemblies and annotation files can be found in figshare: https://figshare.com/articles/dataset/Booklice_Diploid_Genomes. Default software parameters were set during our analyses if unspecified. All pipelines and codes used for data processing can be found in https://github.com/fsq9451/Booklice-Diploid-Genomes.

## Acknowledgements

We would like to express our sincere gratitude to Prof. Tanja Schwander and Dr. Alexander Brandt from the University of Lausanne for their invaluable inspiration and insightful suggestions. Our heartfelt thanks also go to Dr. Chentao Yang from BGI for his expert guidance on inter-haplotypic sequence analyses. We are also deeply grateful for the support and contributions of Dr. Qianqian Yang, Aohan Pang, Youting Pang, Yun Su, Yueyang Zhou, Wenxin Deng, and Jixinang Cui, whose assistance with sample collection, rearing, and management has been essential to this work. This work was supported by the Key Research Program of International Collaboration between China and Czech Republic (2018YFE0108700) to Z.L., the 2115 Talent Development Program of China Agricultural University and the earmarked fund for CARS (grant number: CARS-02-32) to Z.L.

## Notes

**Competing Interest Statement:** The authors declare that they have no conflict of interest.

### Competing Interest Statement

The authors have declared no competing interest.

