## Supplementary Table for "Diploid genomes of the booklouse reveal evolutionary consequences of asexuality": MS_Supplementary.docx

**Figures**

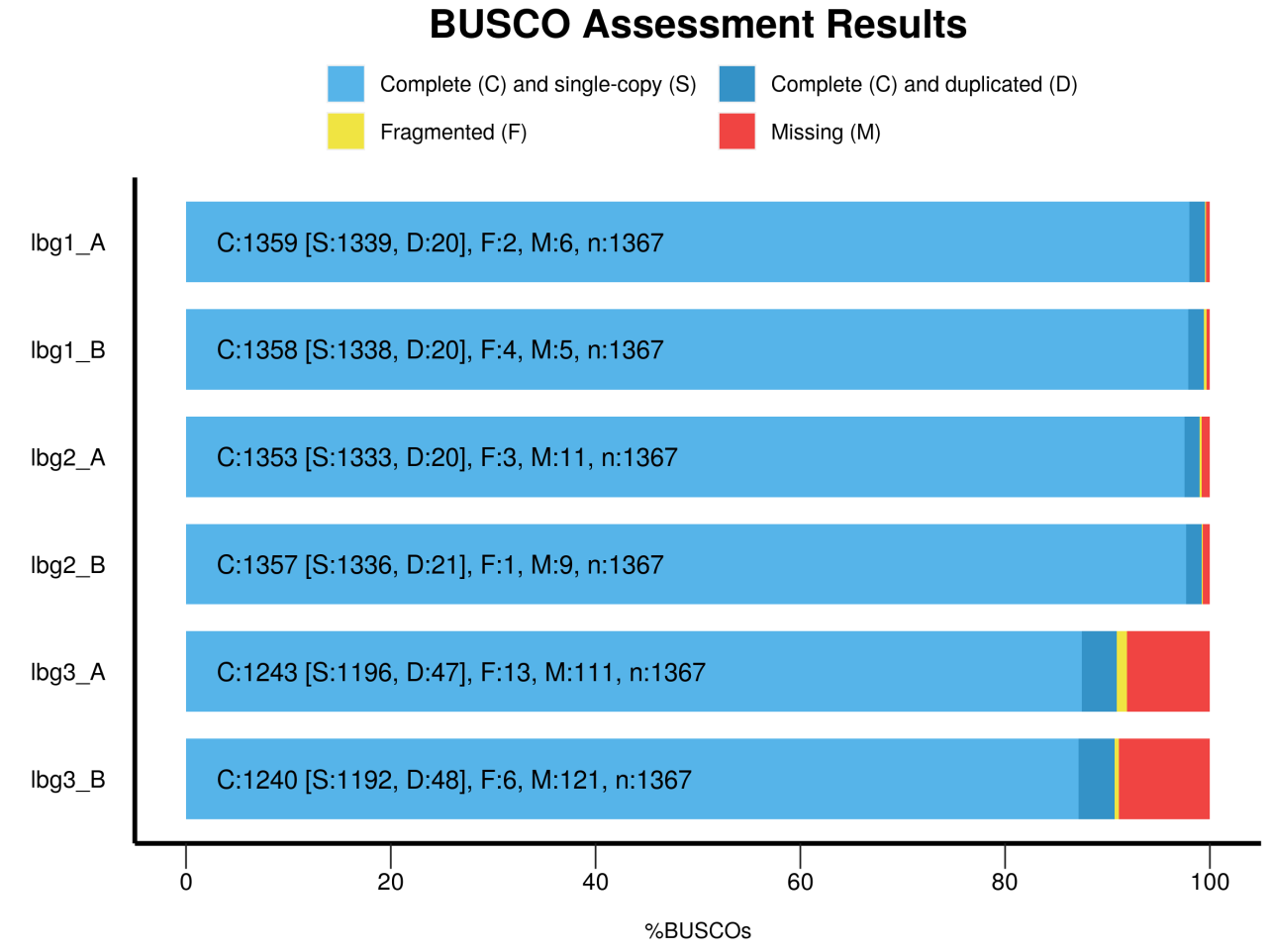

**Fig. S1 BUSCO completeness of six haplotypes of *Liposcelis bostrychophila*.**

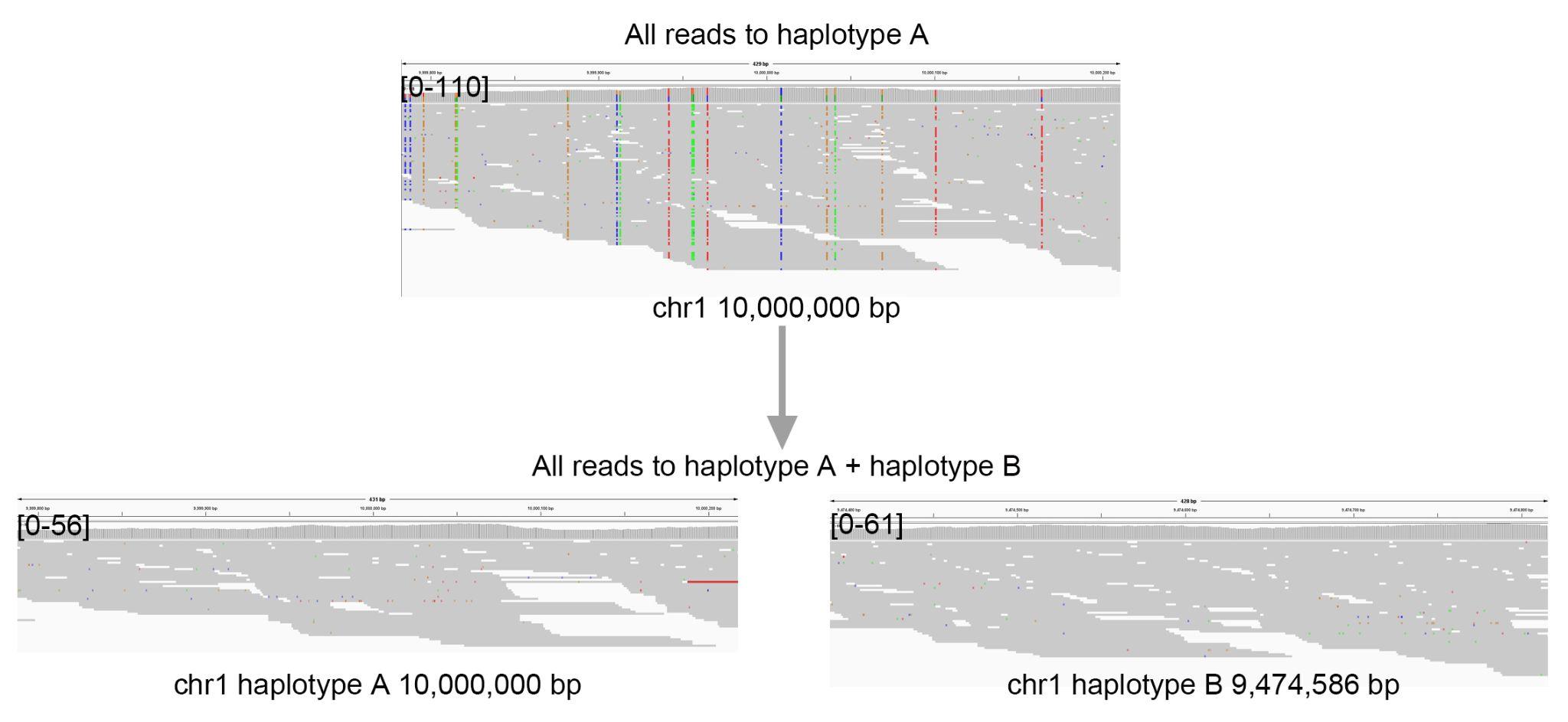

**Fig. S2 Illumina sequencing data remapping to haplotypic or diploid assembly.**

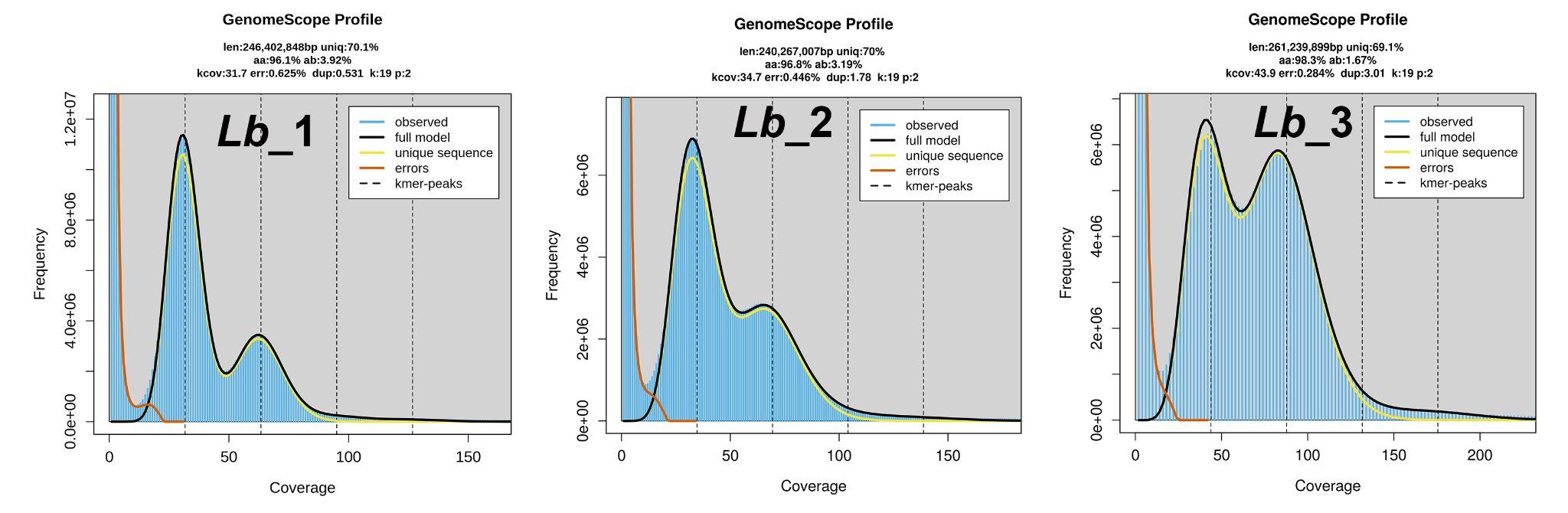

**Fig. S3 Genome size and heterozygosity estimation using GenomeScope.**

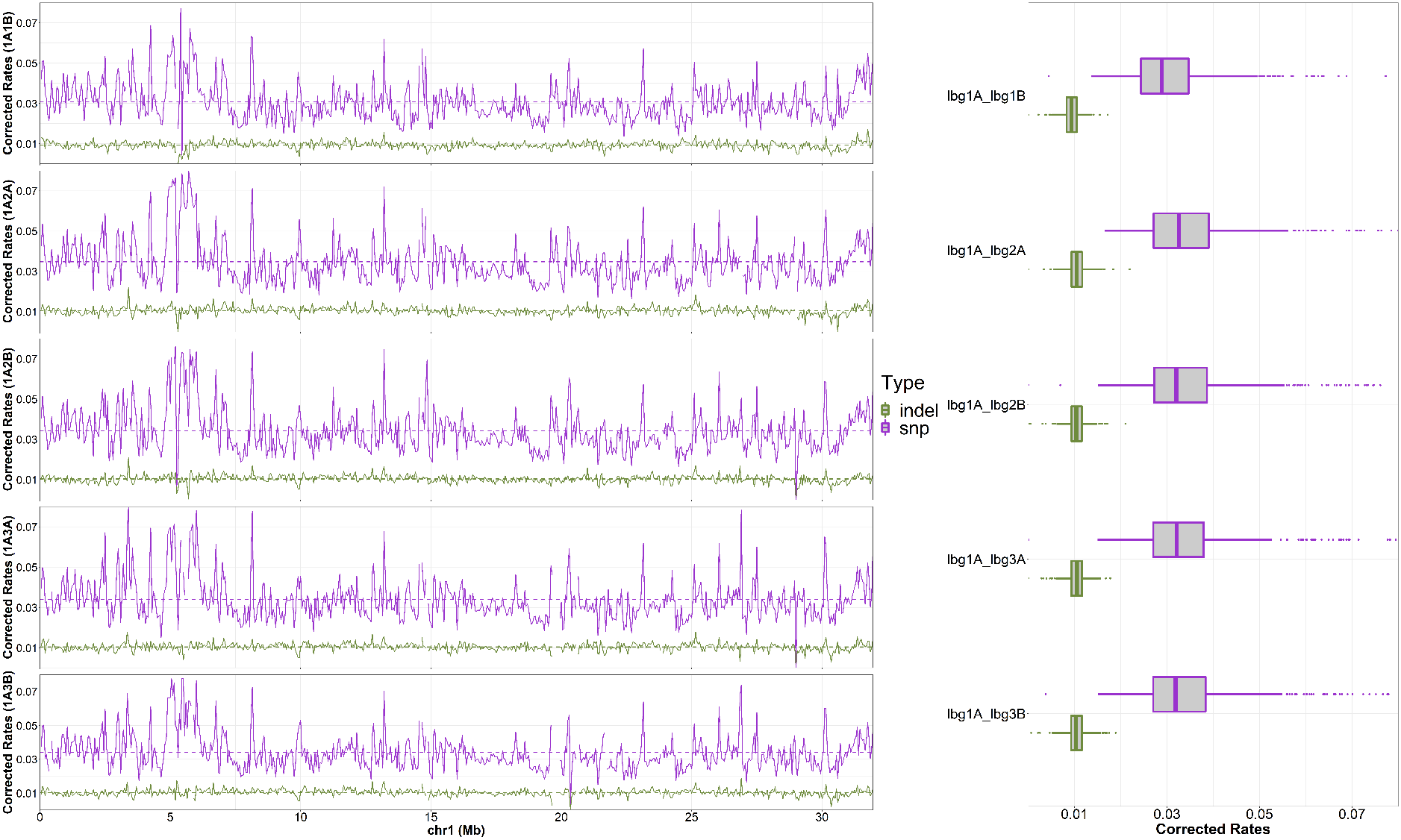

**Fig. S4 SNPs and INDELs based mapping to Lb_1A chromosome 1.**

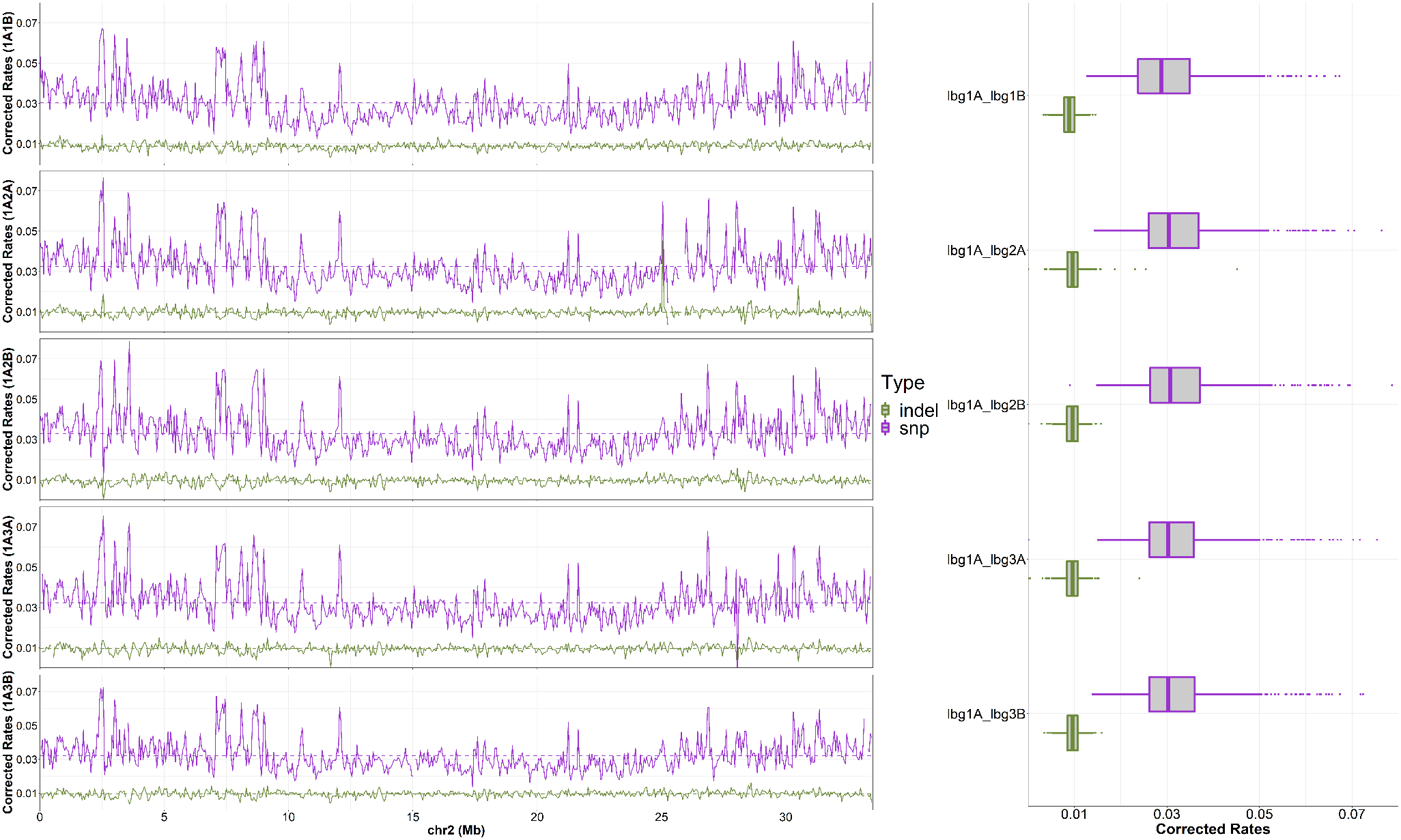

**Fig. S5 SNPs and INDELs based mapping to Lb_1A chromosome 2.**

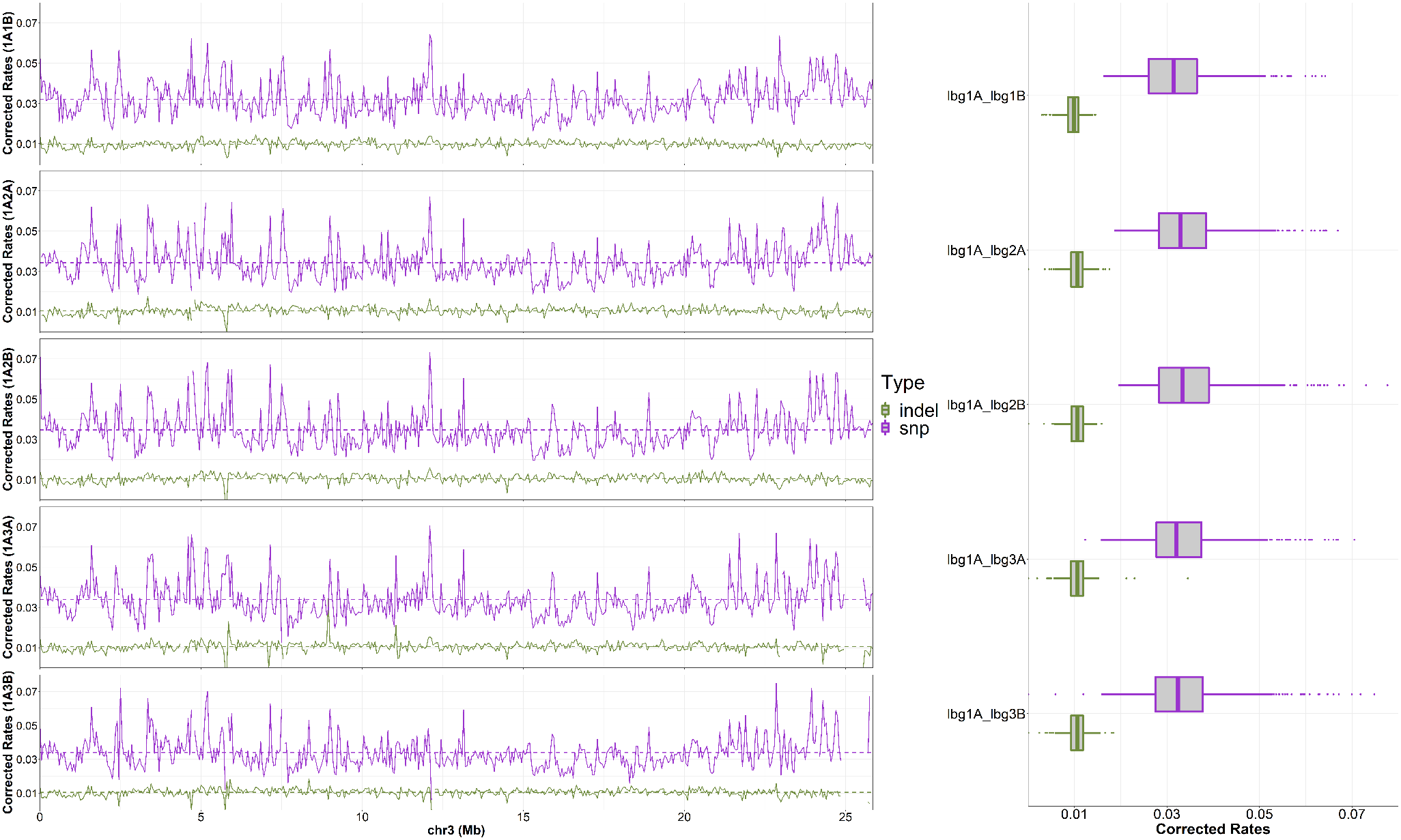

**47Fig. S6 SNPs and INDELs based mapping to Lb_1A chromosome 3.**

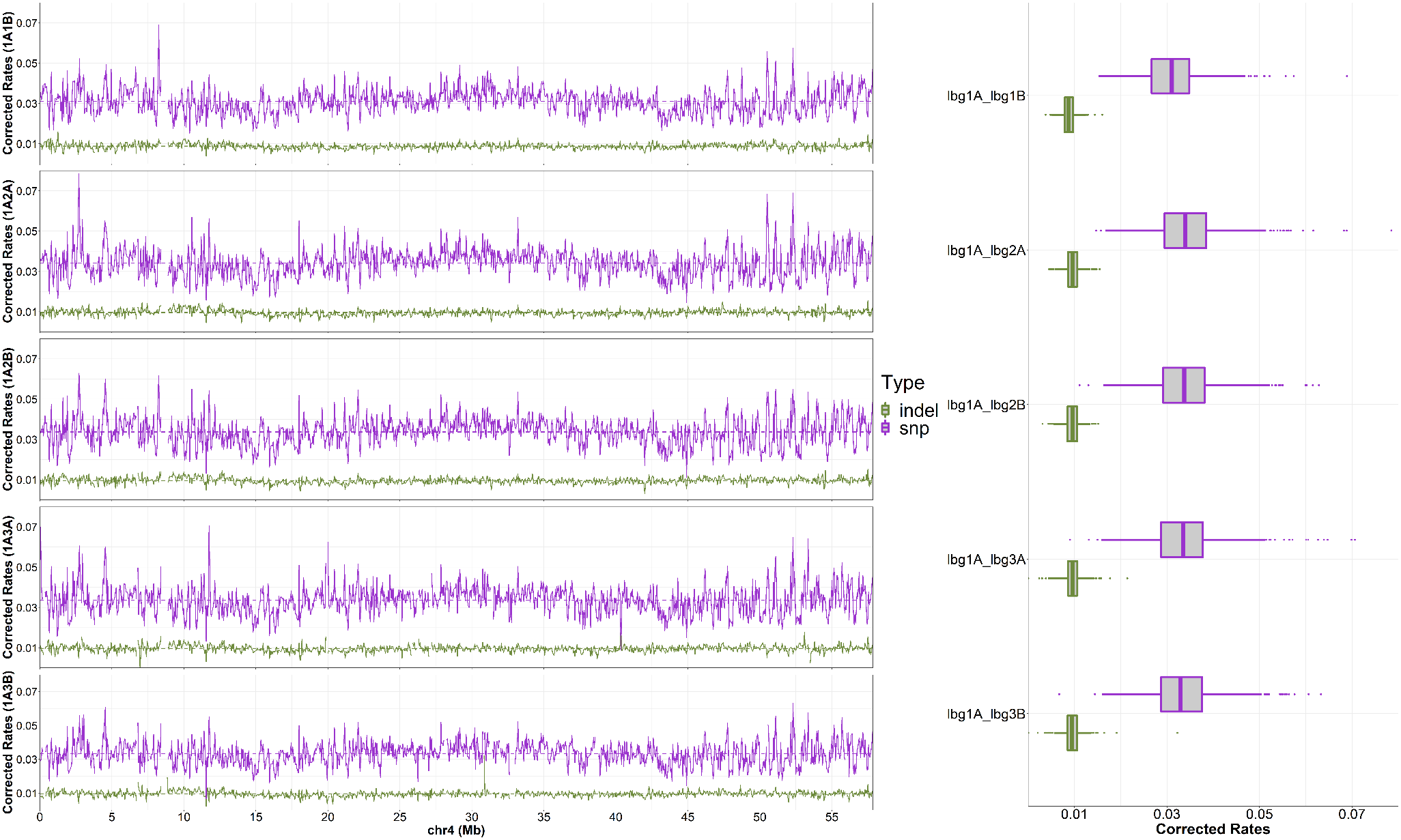

**Fig. S7 SNPs and INDELs based mapping to Lb_1A chromosome 4.**

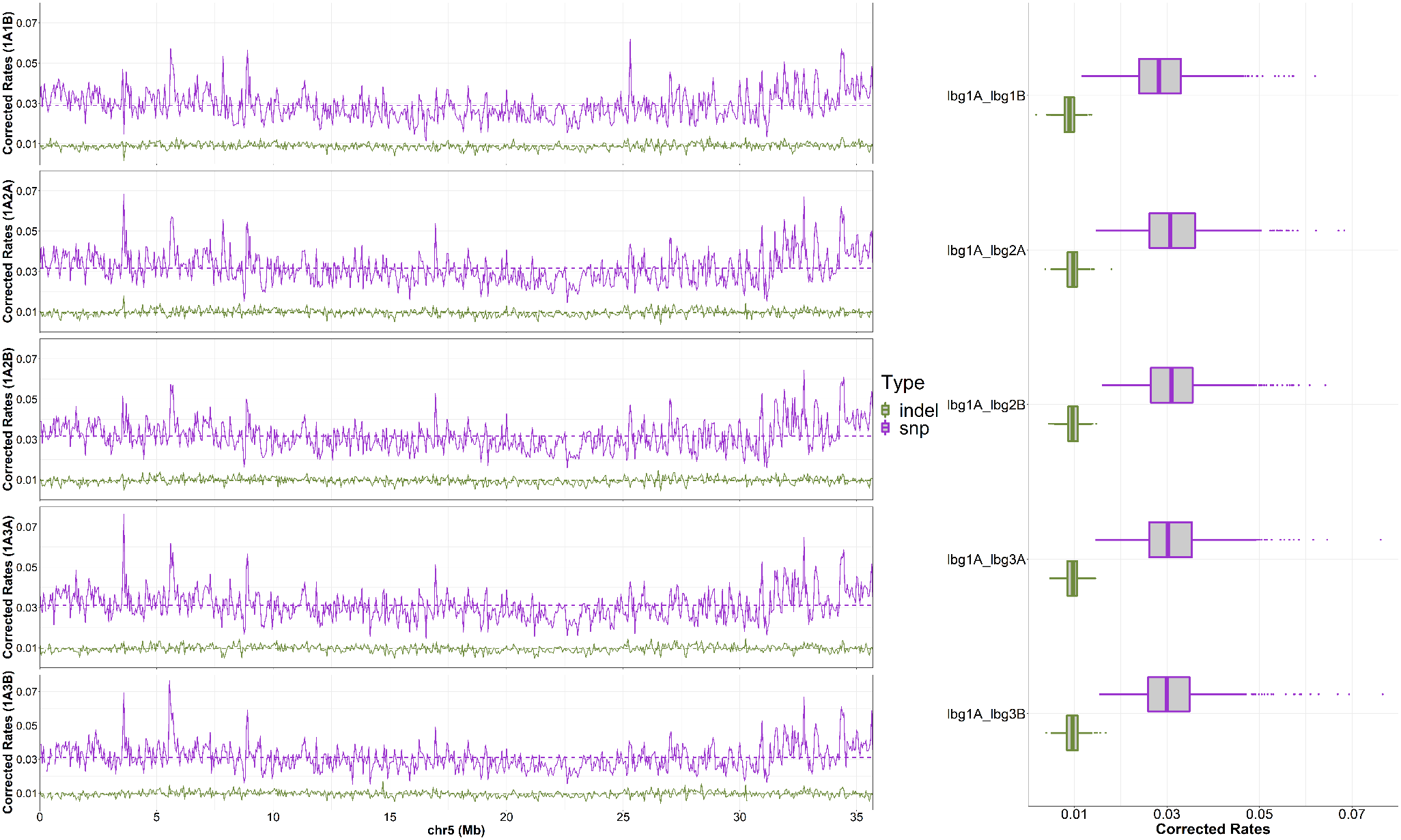

**Fig. S8 SNPs and INDELs based mapping to Lb_1A chromosome 5.**

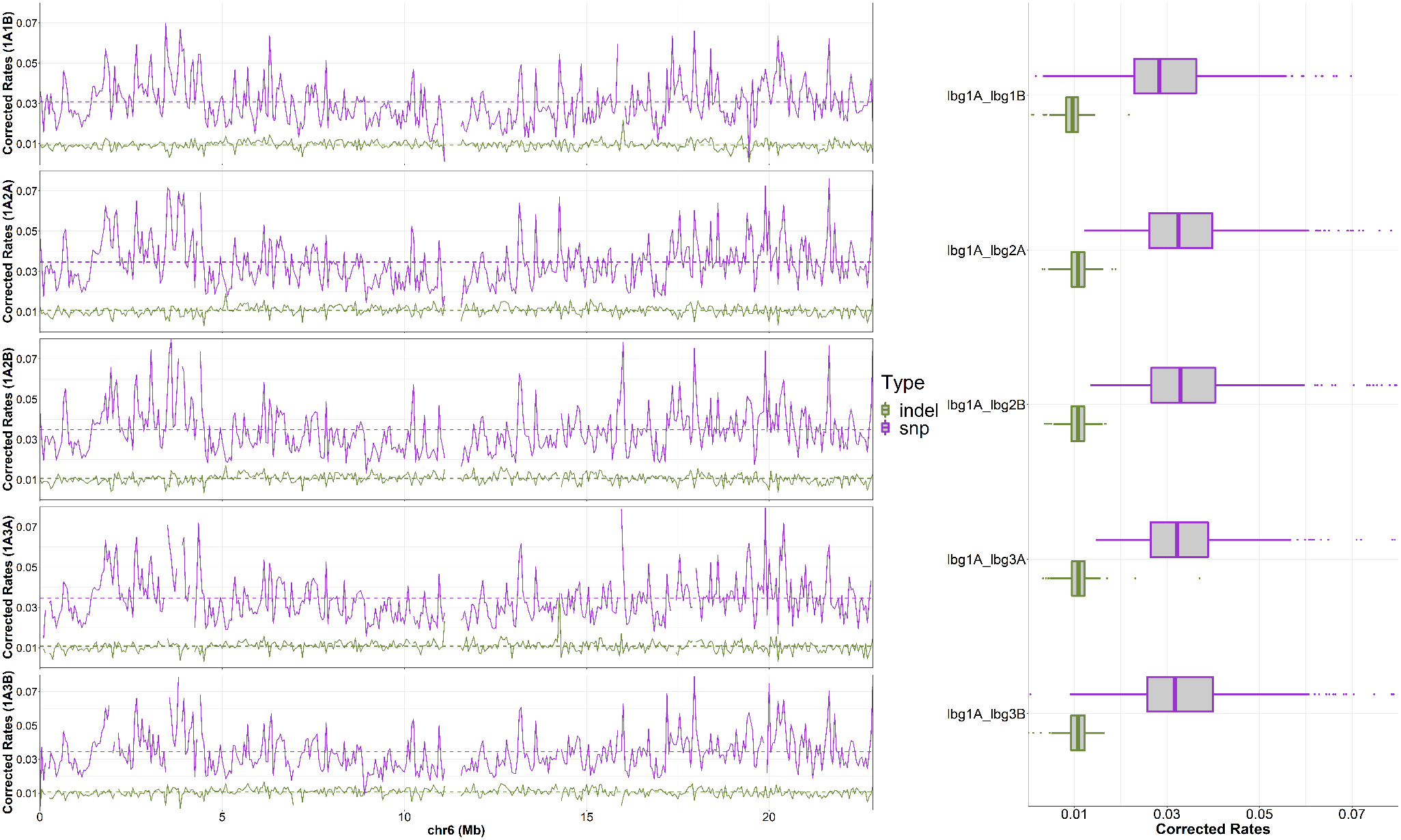

**Fig. S9 SNPs and INDELs based mapping to Lb_1A chromosome 6.**

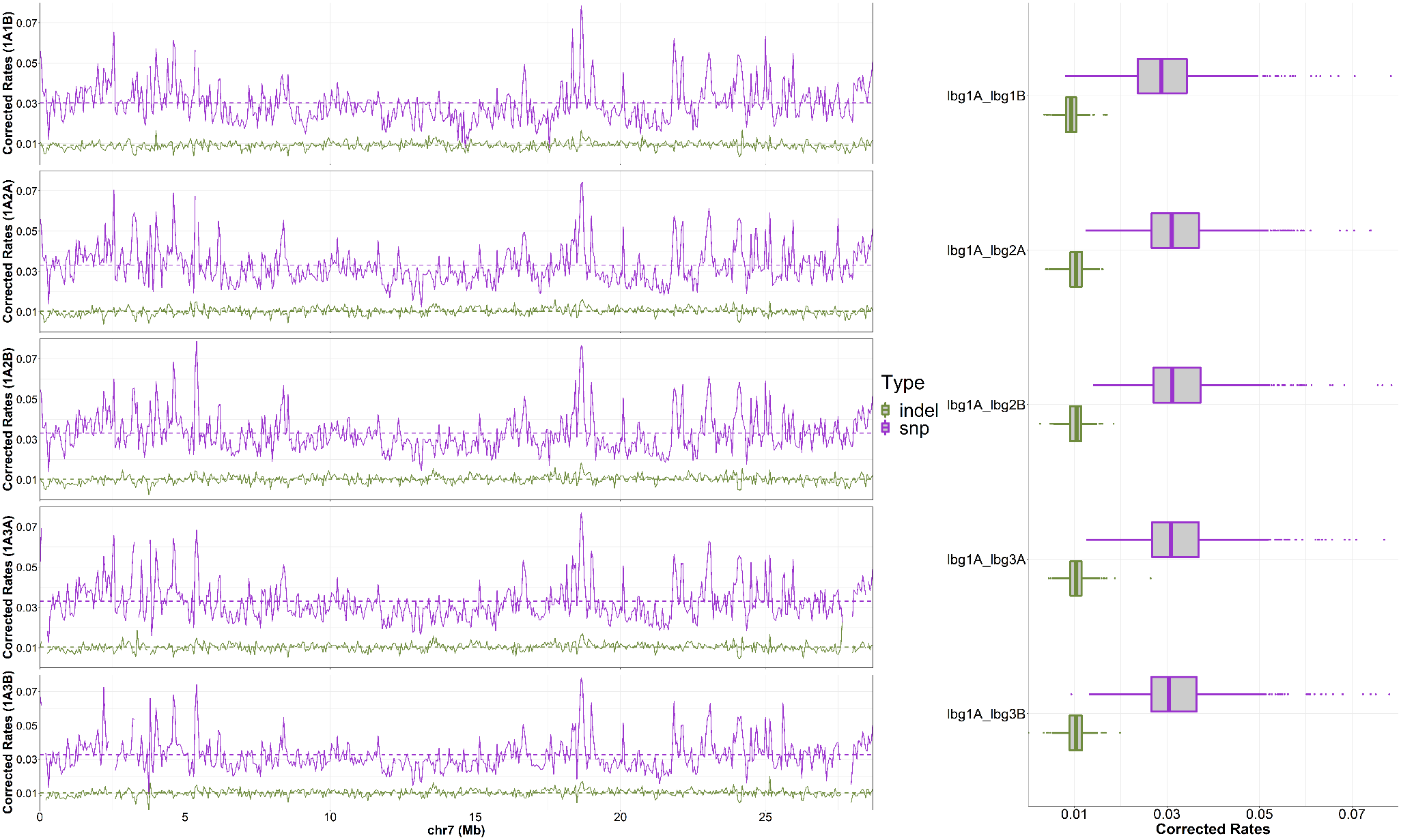

**Fig. S10 SNPs and INDELs based mapping to Lb_1A chromosome 7.**

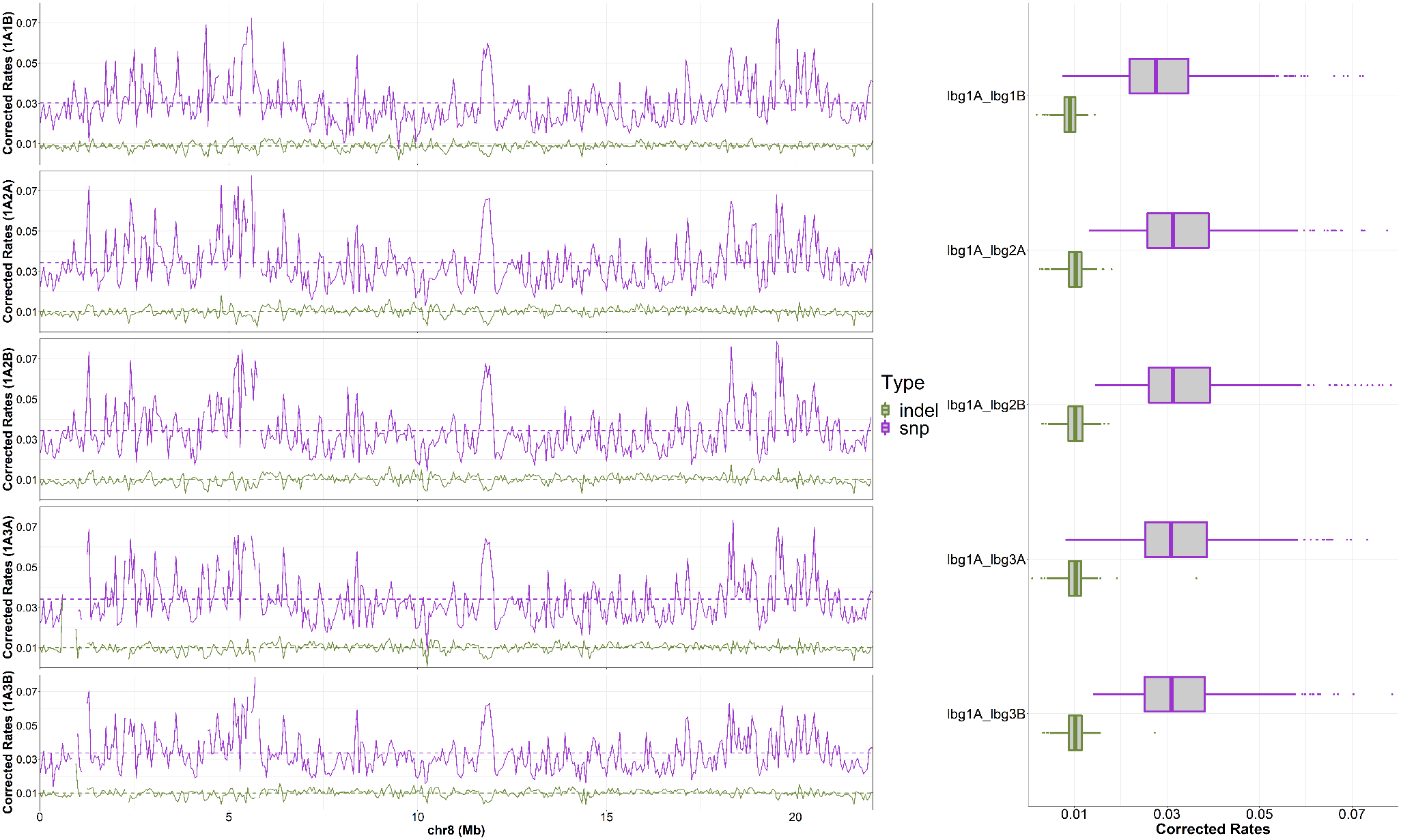

**Fig. S11 SNPs and INDELs based mapping to Lb_1A chromosome 8.**

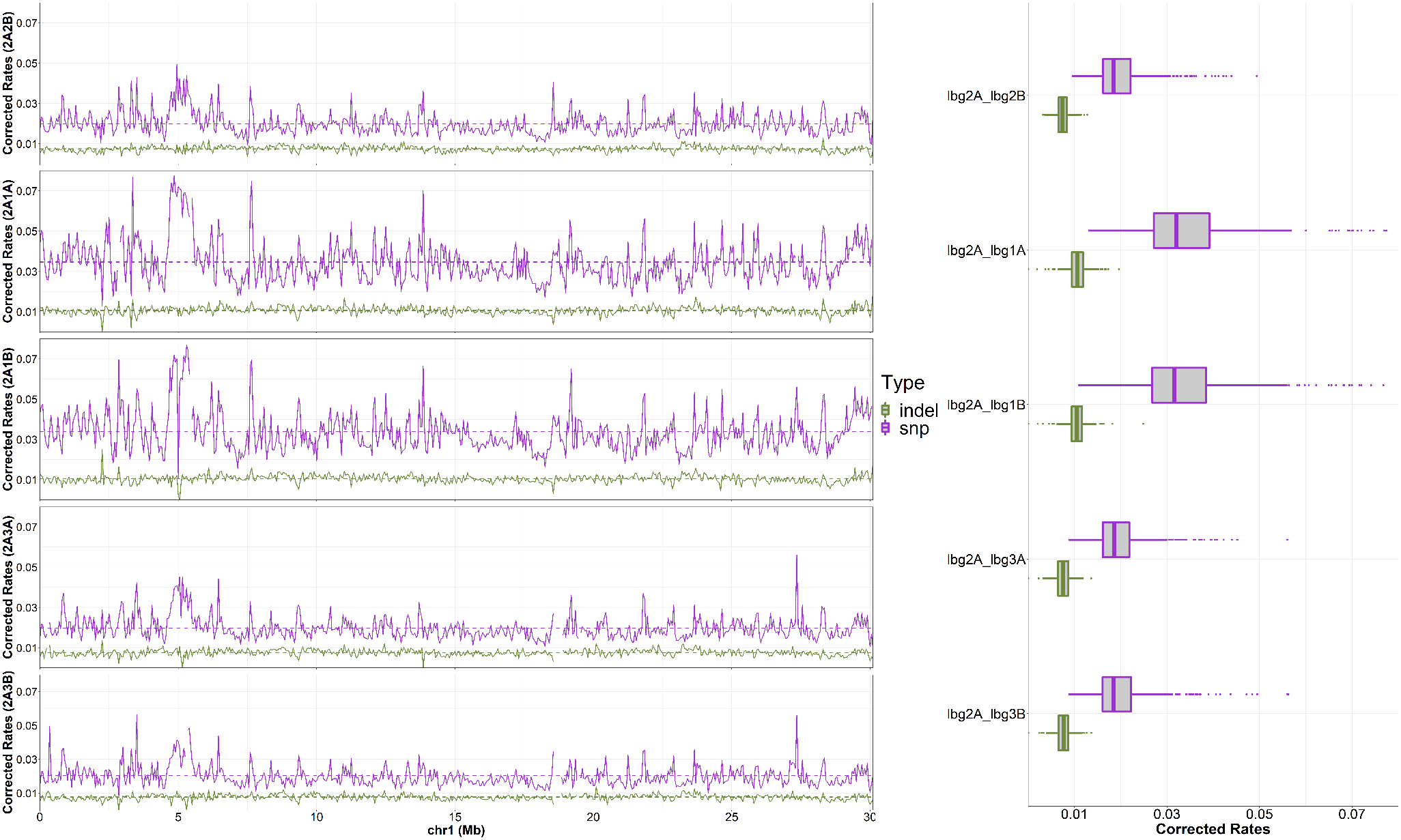

**Fig. S12 SNPs and INDELs based mapping to Lb_2A chromosome 1.**

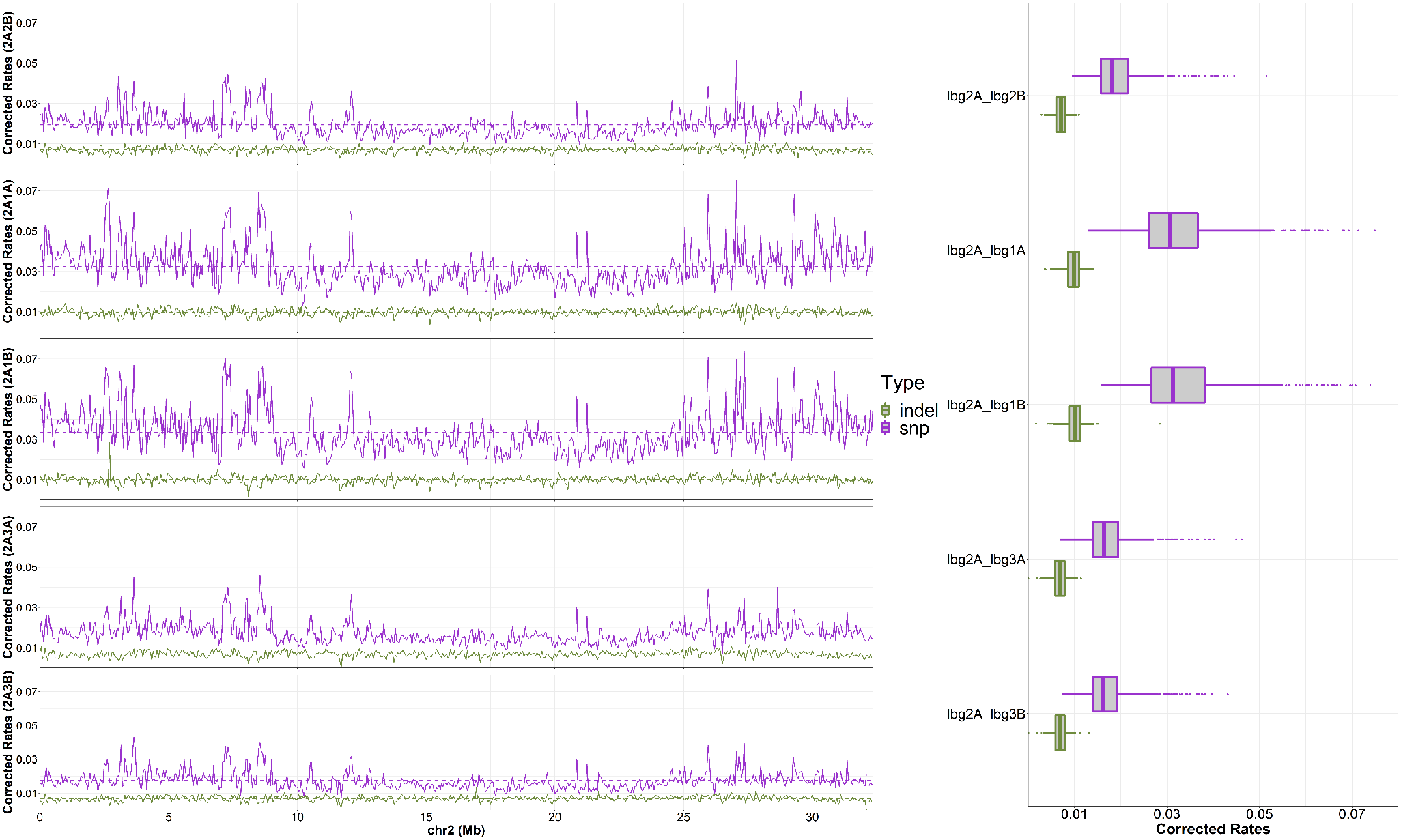

**Fig. S13 SNPs and INDELs based mapping to Lb_2A chromosome 2.**

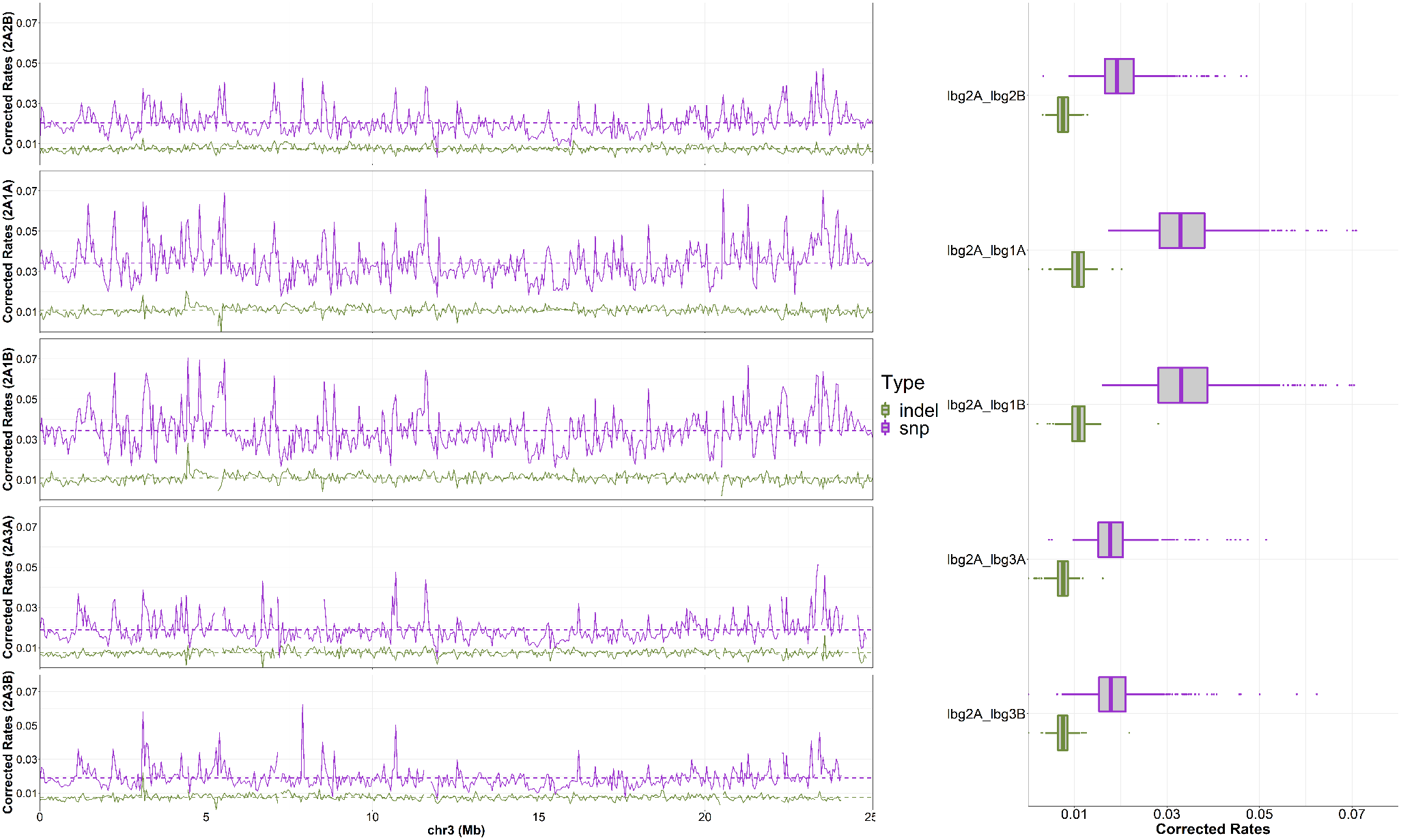

**Fig. S14 SNPs and INDELs based mapping to Lb_2A chromosome 3.**

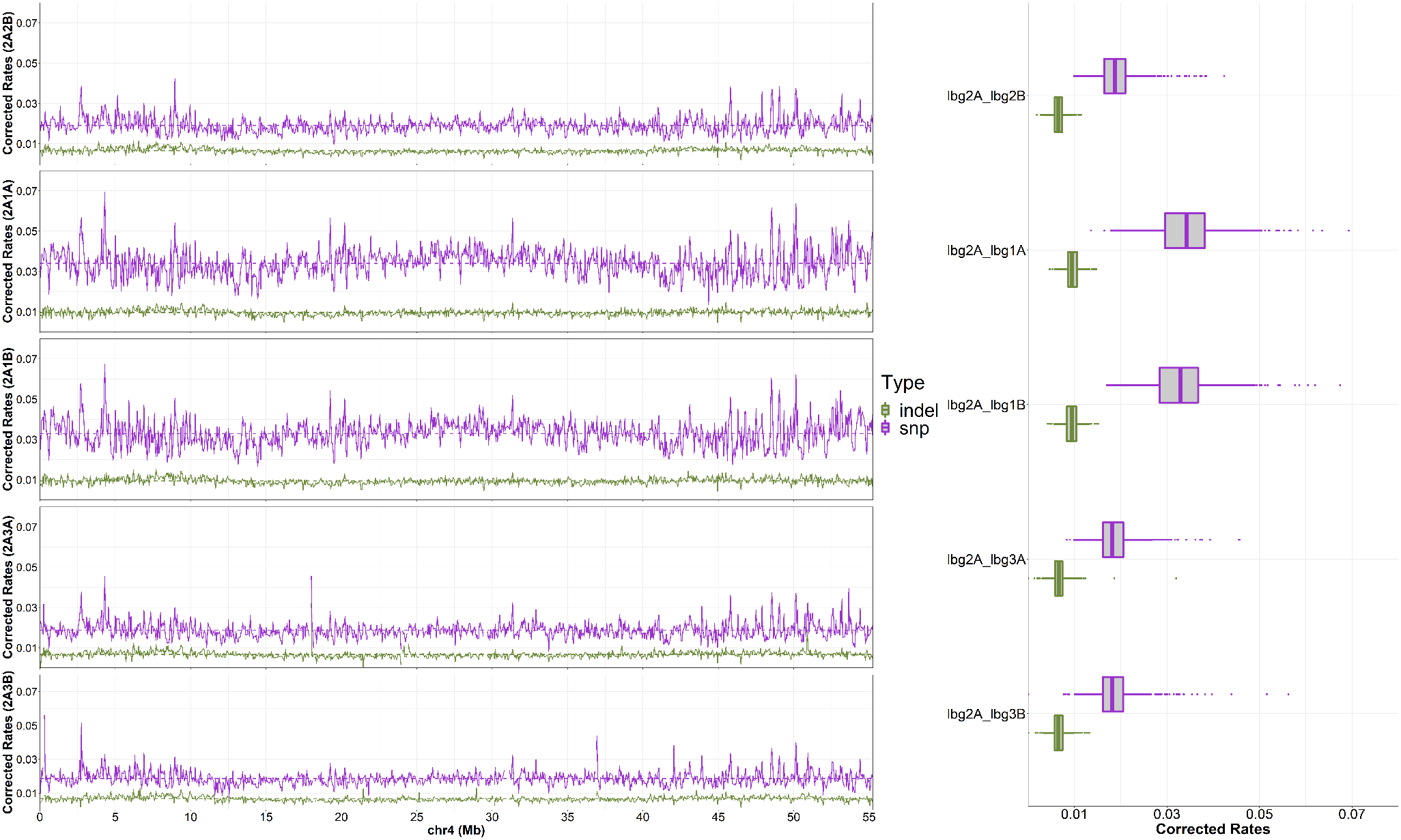

**Fig. S15 SNPs and INDELs based mapping to Lb_2A chromosome 4.**

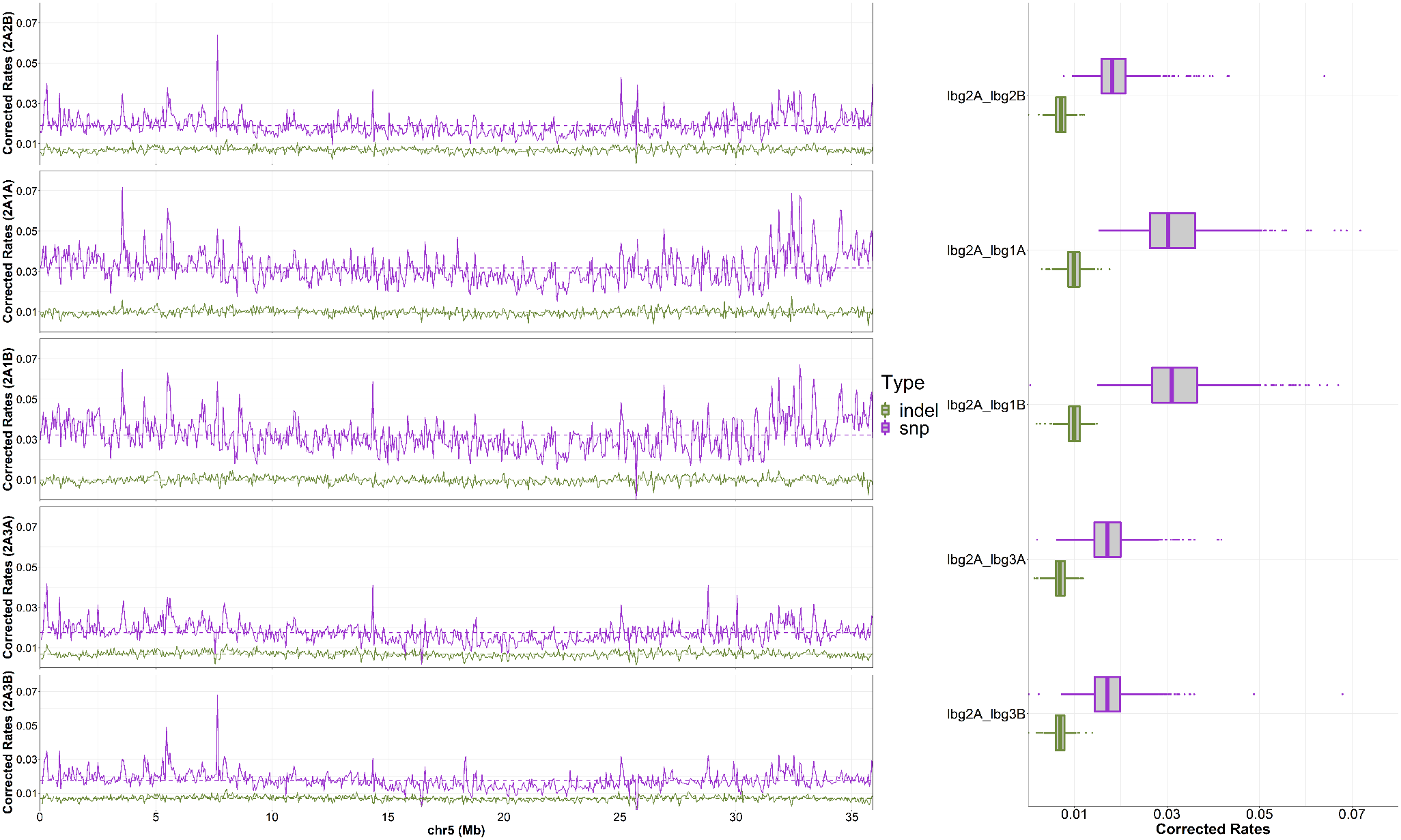

**Fig. S16 SNPs and INDELs based mapping to Lb_2A chromosome 5.**

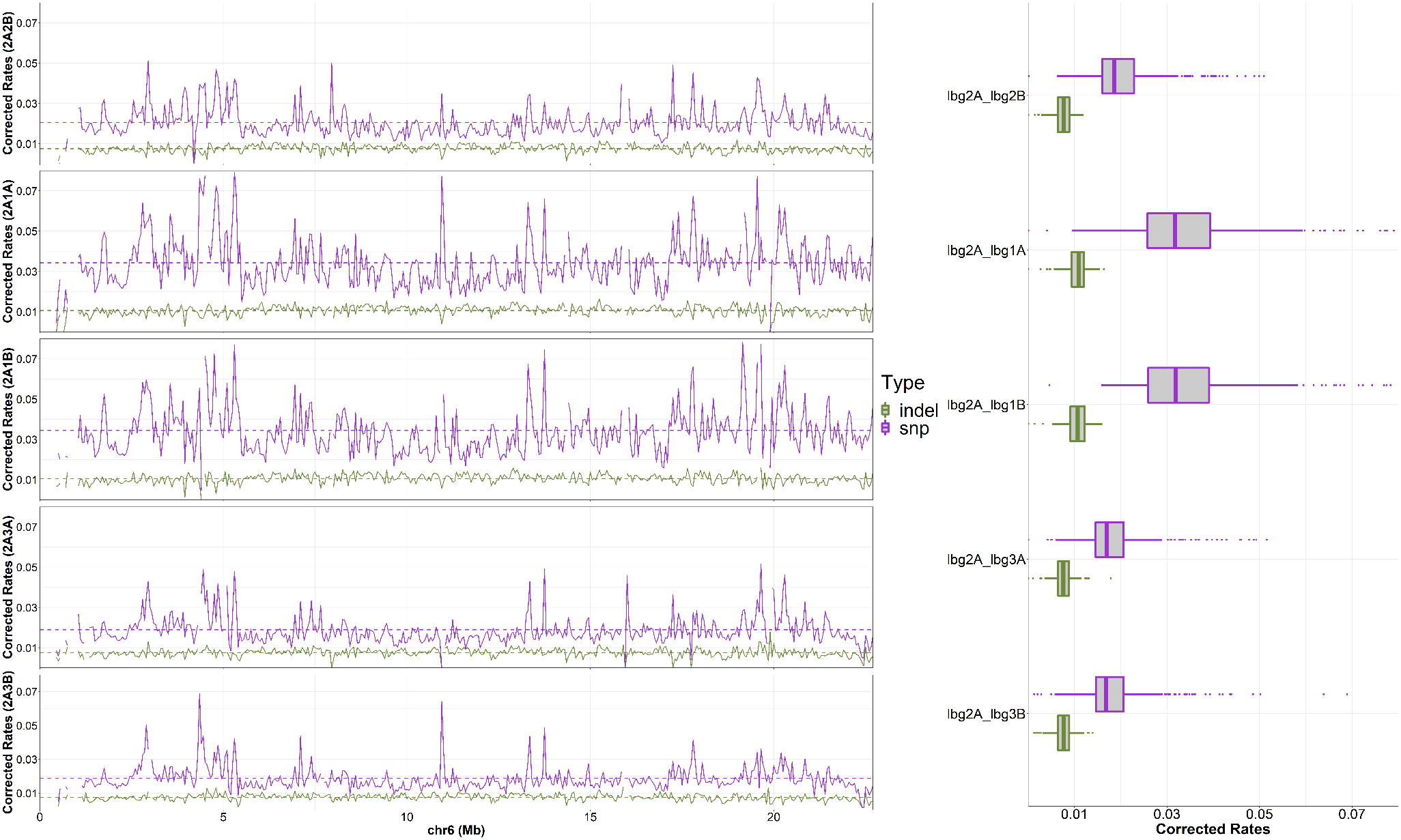

**Fig. S17 SNPs and INDELs based mapping to Lb_2A chromosome 6.**

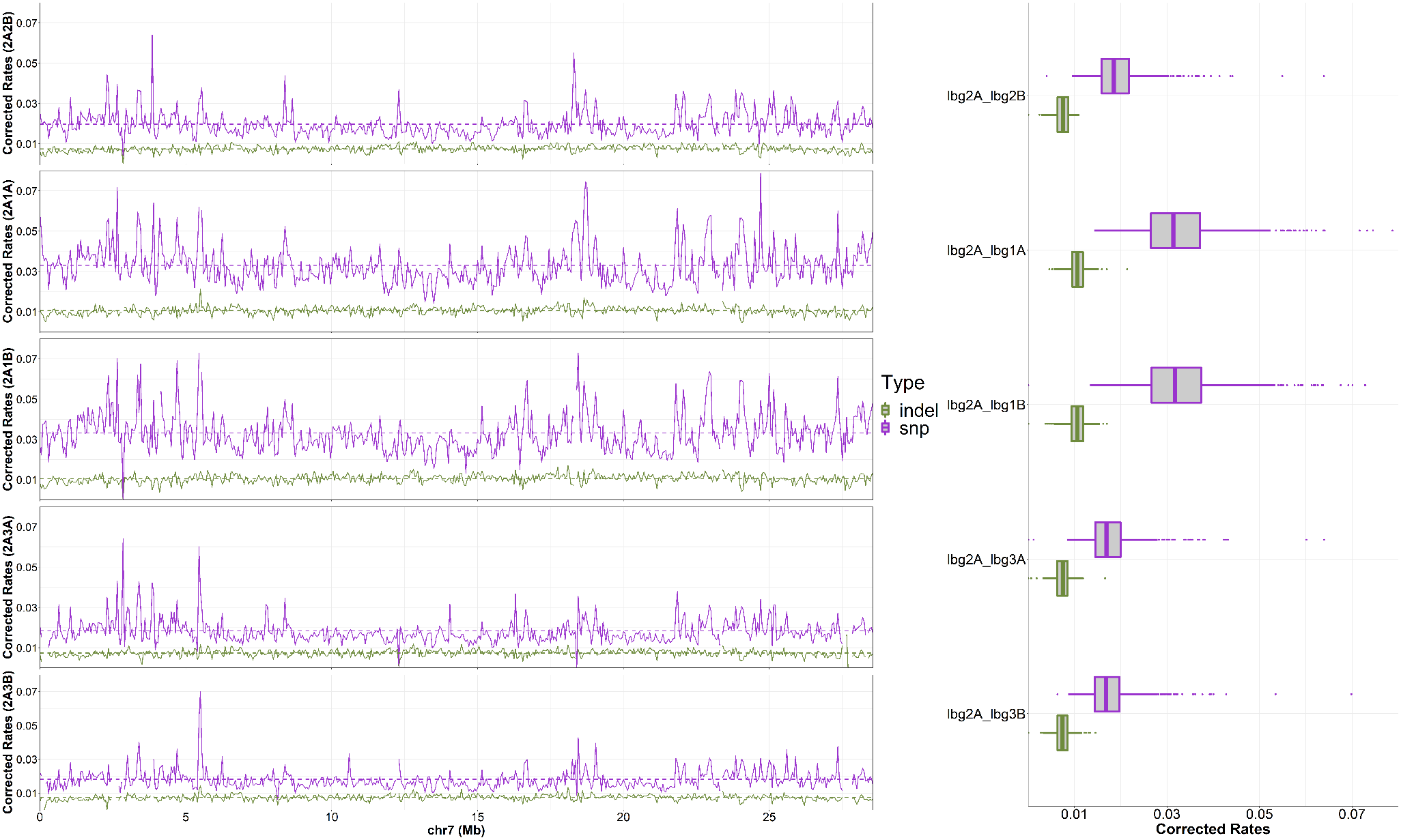

**Fig. S18 SNPs and INDELs based mapping to Lb_2A chromosome 7.**

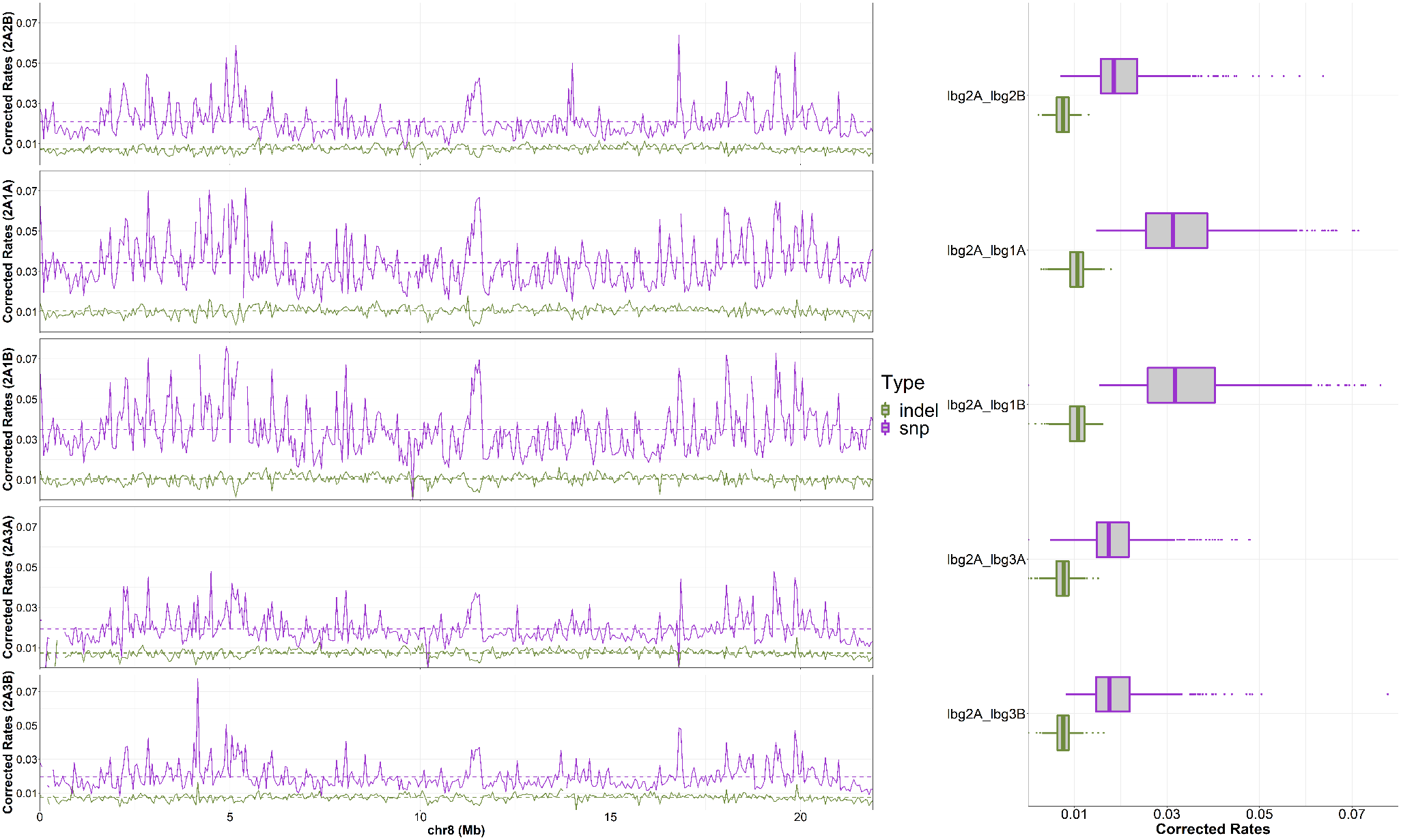

**Fig. S19 SNPs and INDELs based mapping to Lb_2A chromosome 8.**

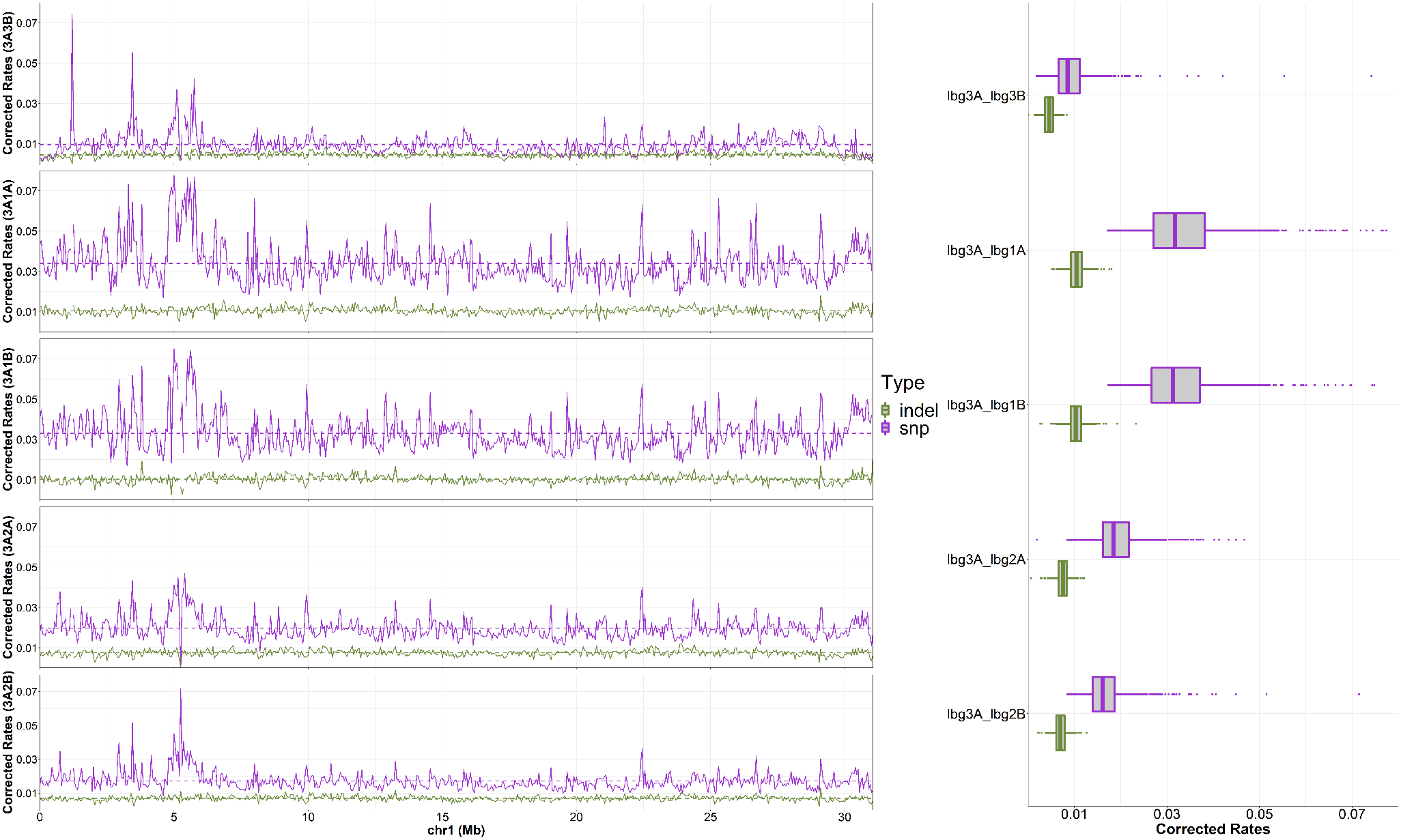

**Fig. S20 SNPs and INDELs based mapping to Lb_3A chromosome 1.**

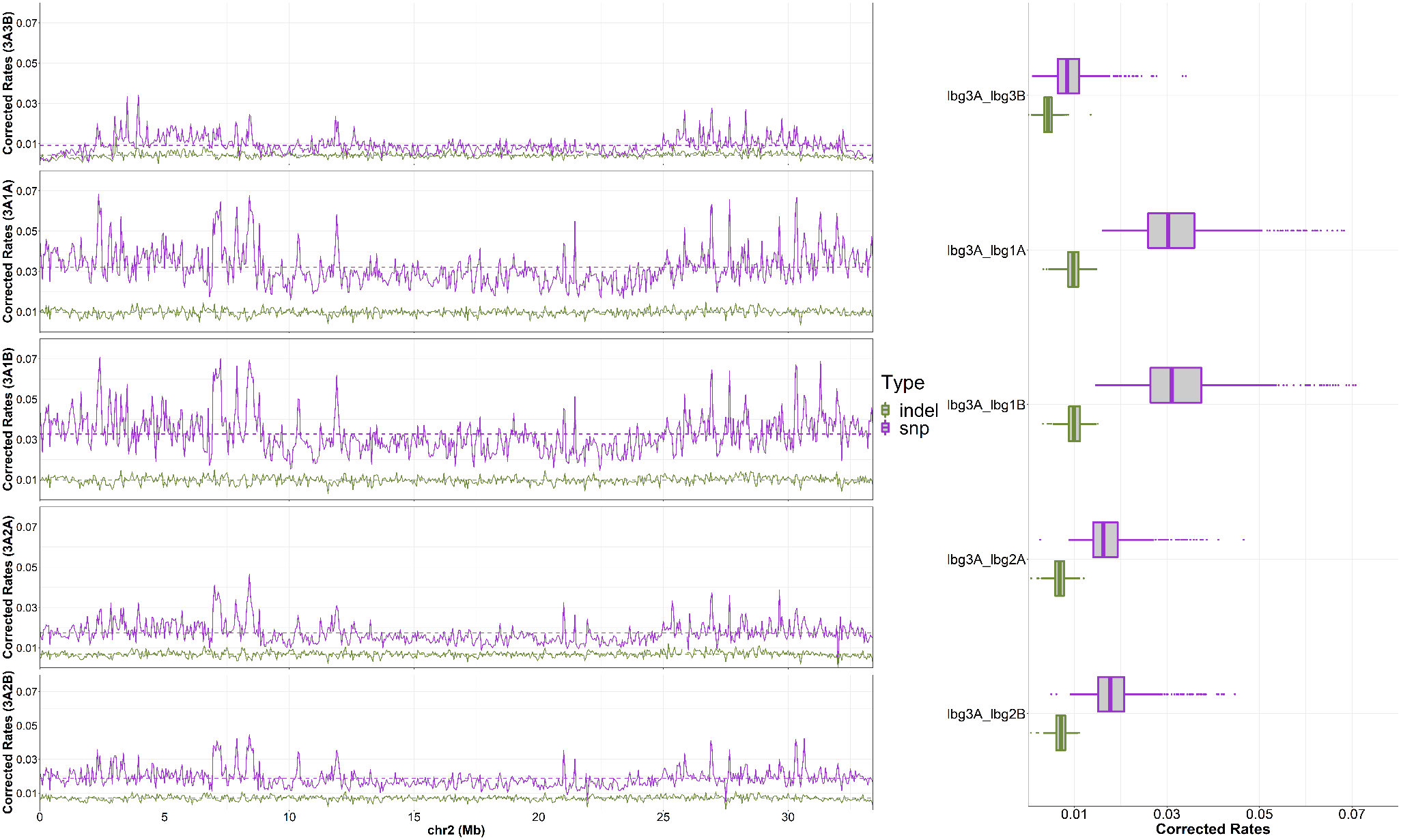

**Fig. S21 SNPs and INDELs based mapping to Lb_3A chromosome 2.**

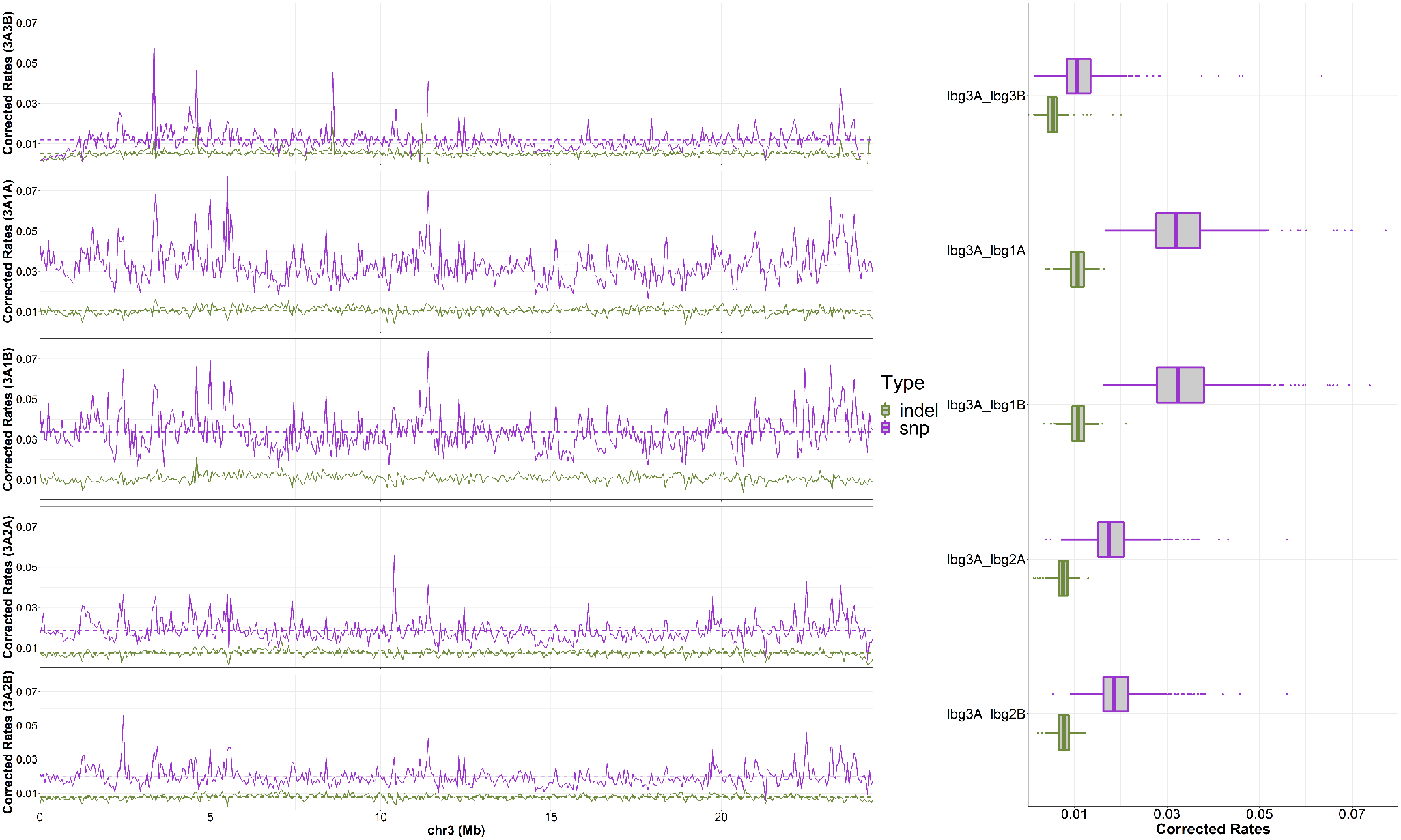

**Fig. S22 SNPs and INDELs based mapping to Lb_3A chromosome 3.**

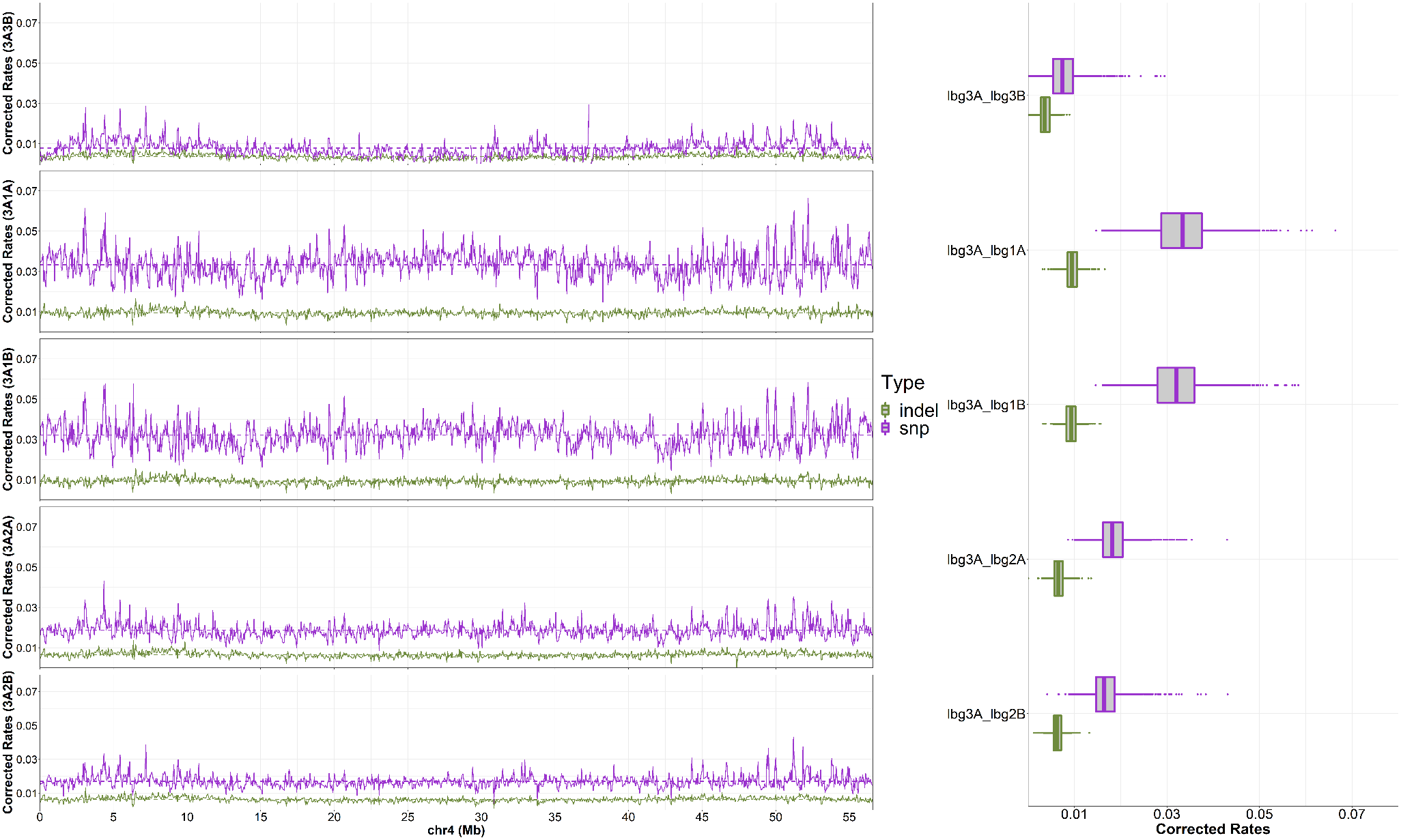

**Fig. S23 SNPs and INDELs based mapping to Lb_3A chromosome 4.**

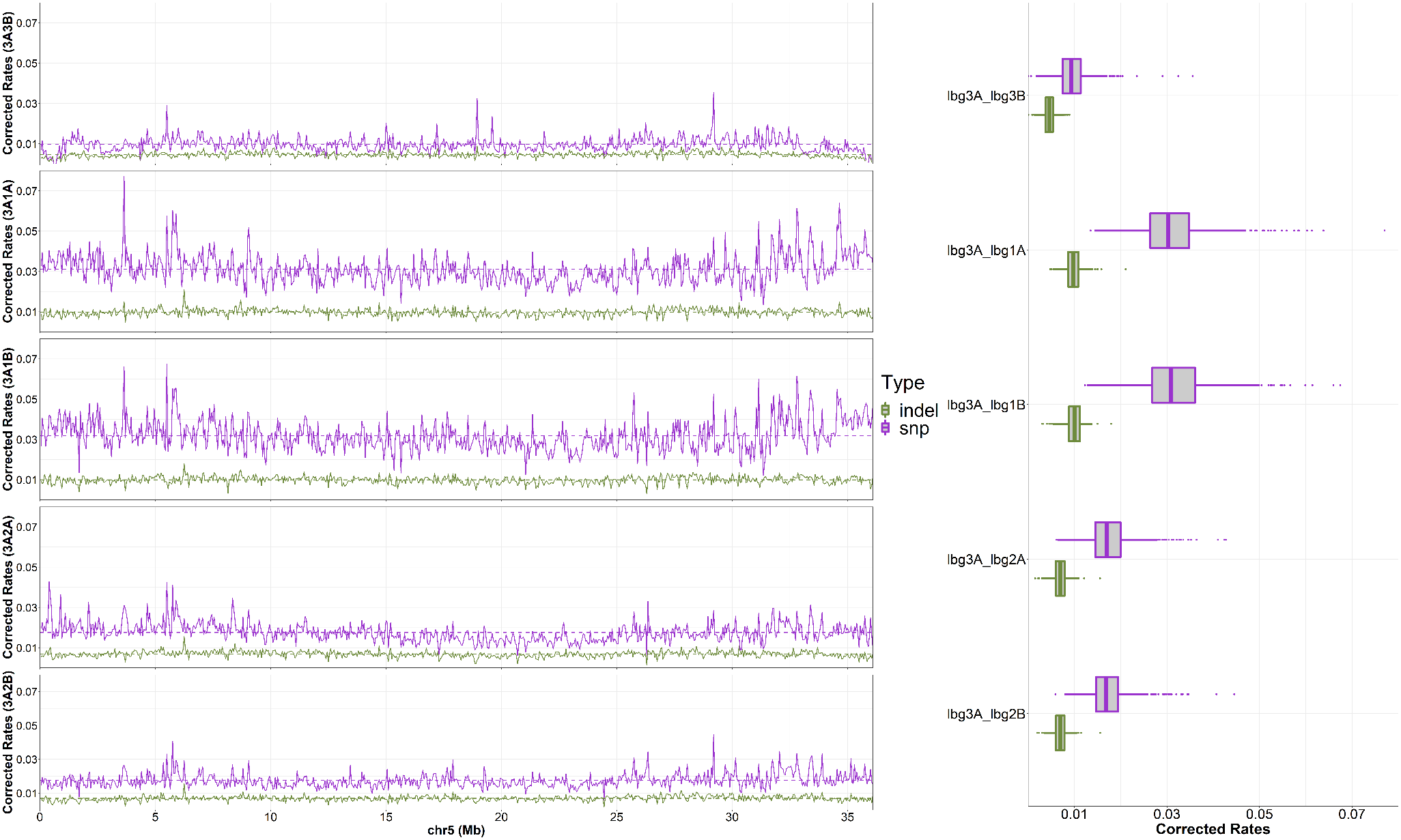

**Fig. S24 SNPs and INDELs based mapping to Lb_3A chromosome 5.**

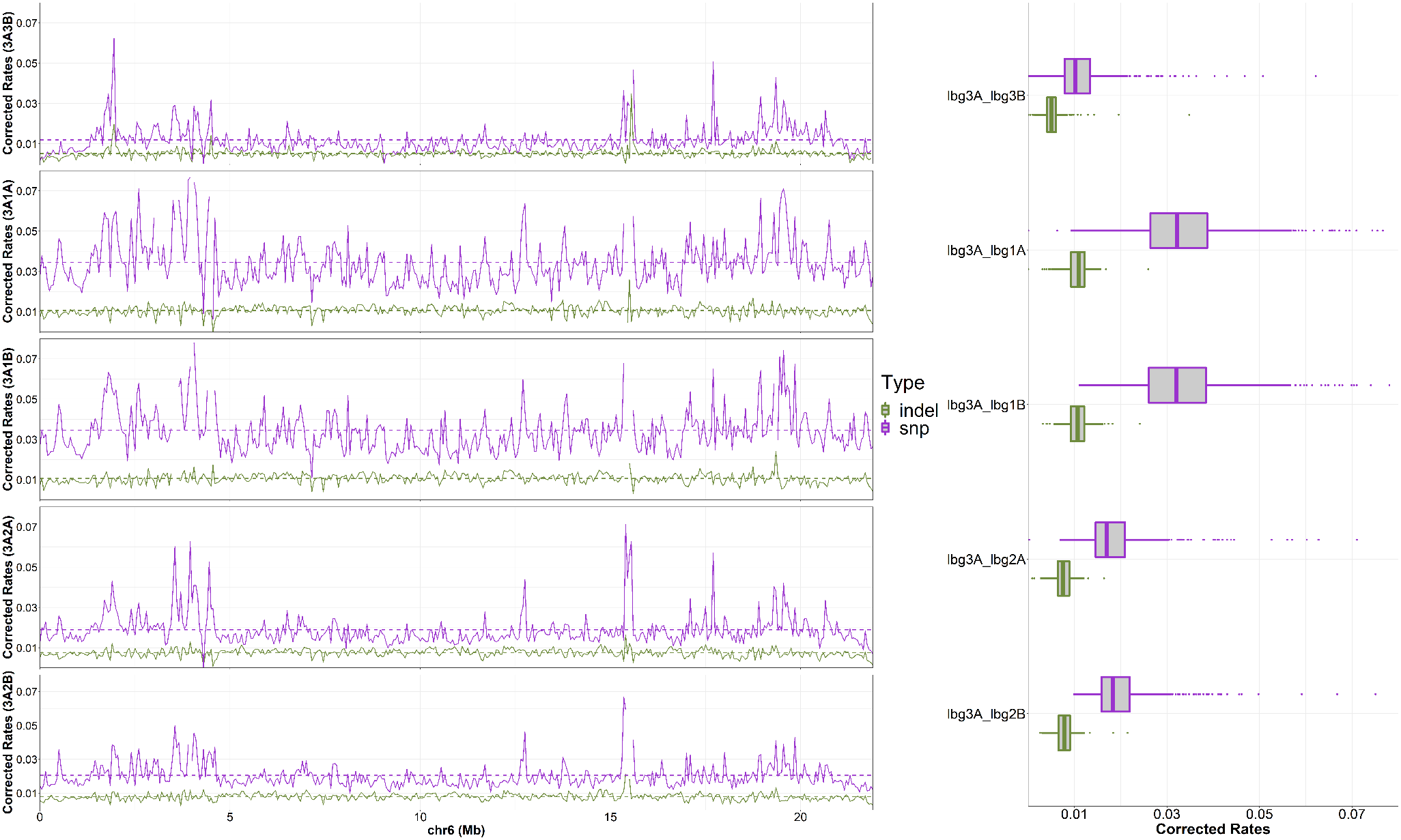

**Fig. S25 SNPs and INDELs based mapping to Lb_3A chromosome 6.**

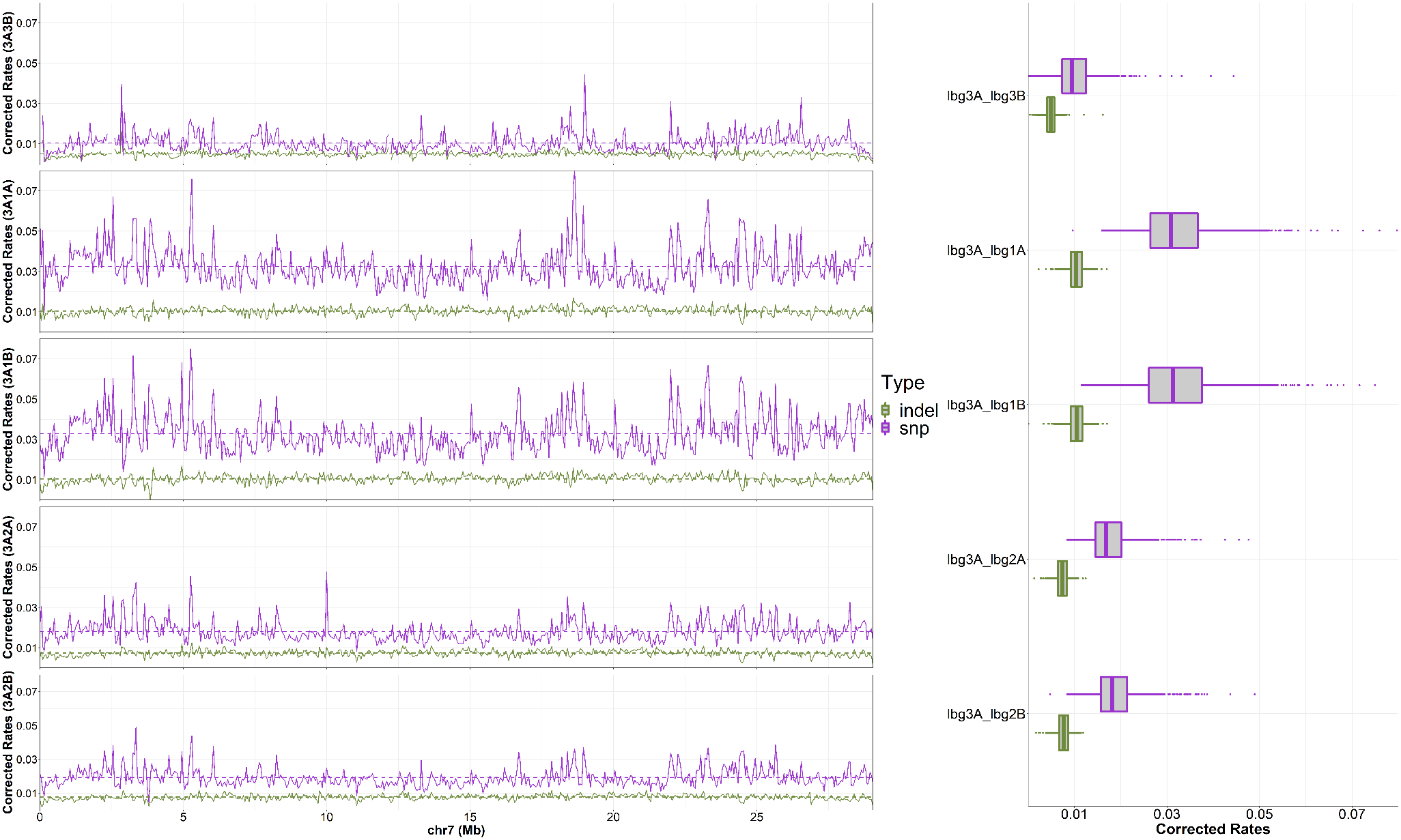

**Fig. S26 SNPs and INDELs based mapping to Lb_3A chromosome 7.**

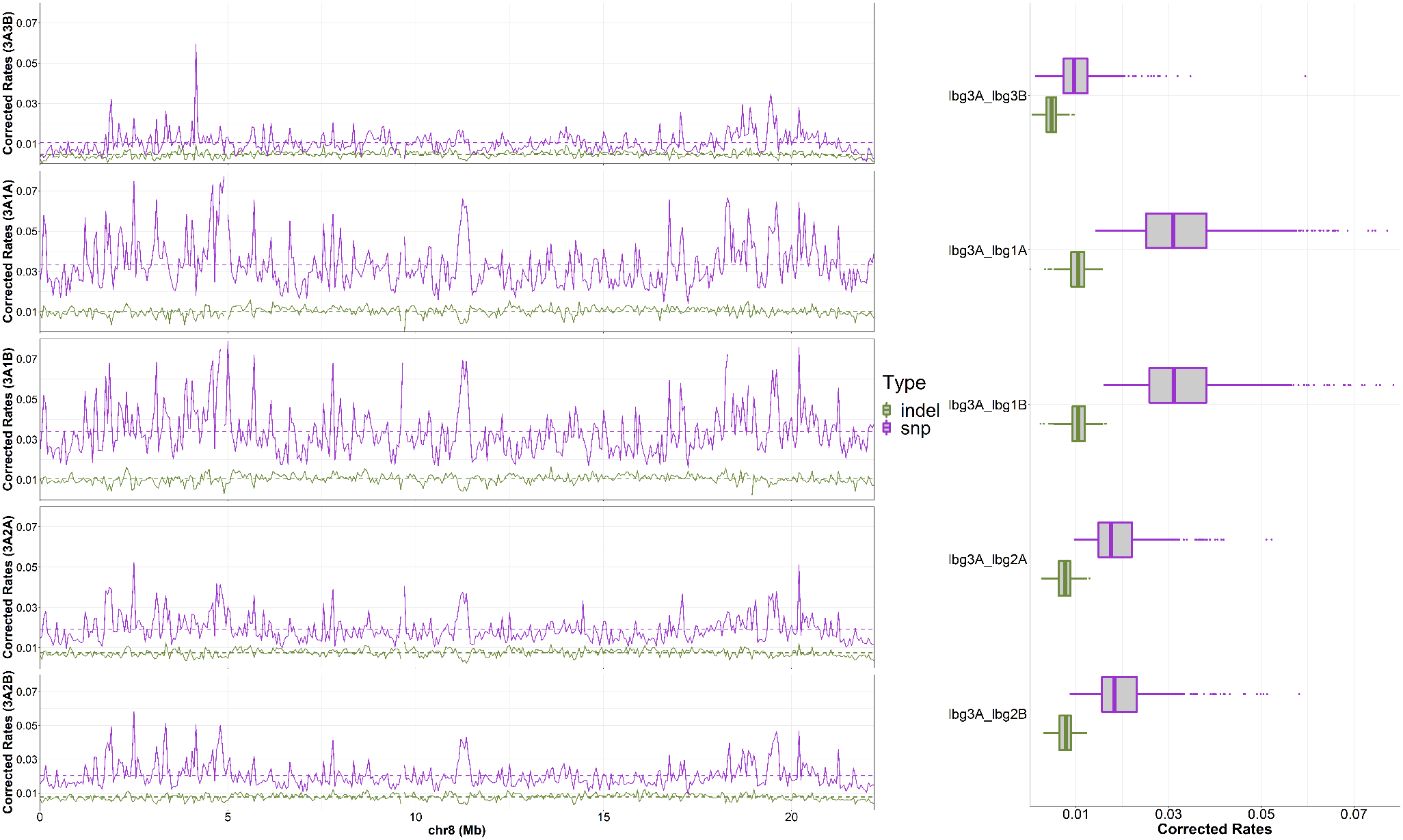

**Fig. S27 SNPs and INDELs based mapping to Lb_3A chromosome 8.**

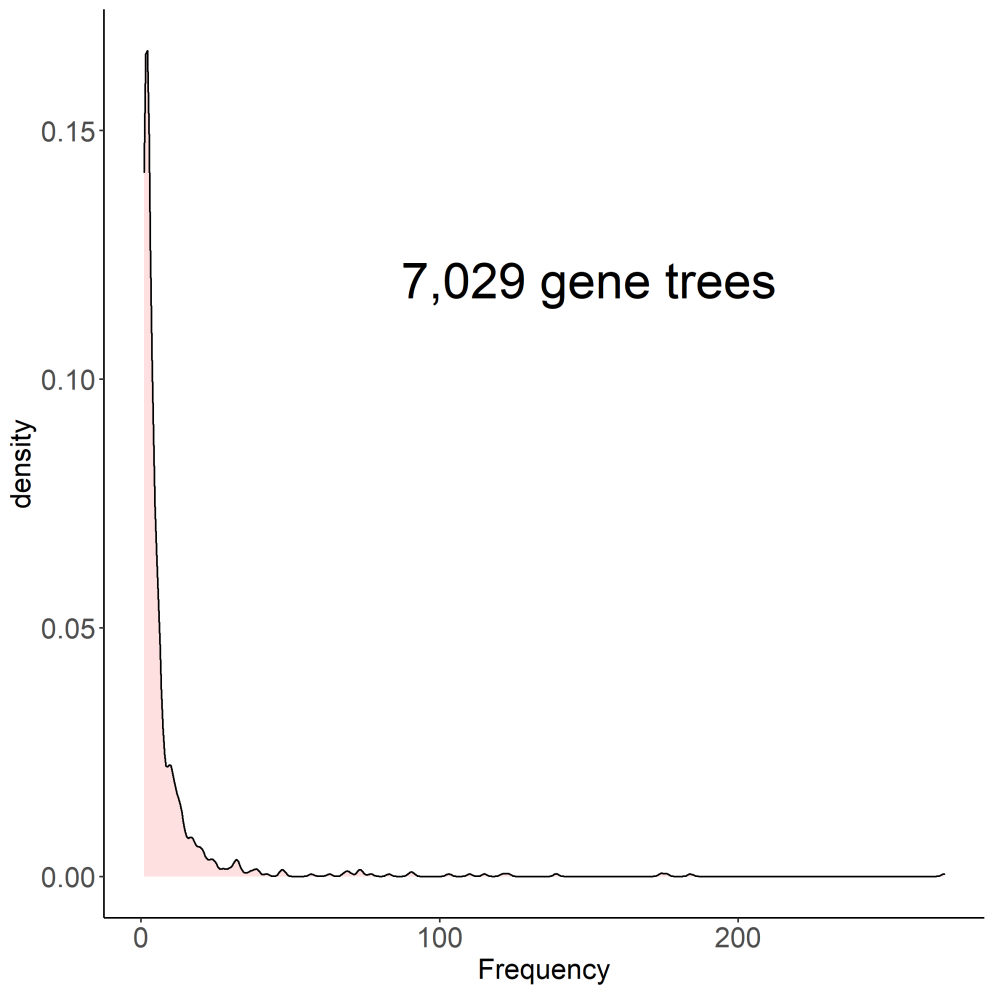

**Fig. S28 Density of phylogenetic trees constructed by each of 7,029 allelic genes.**

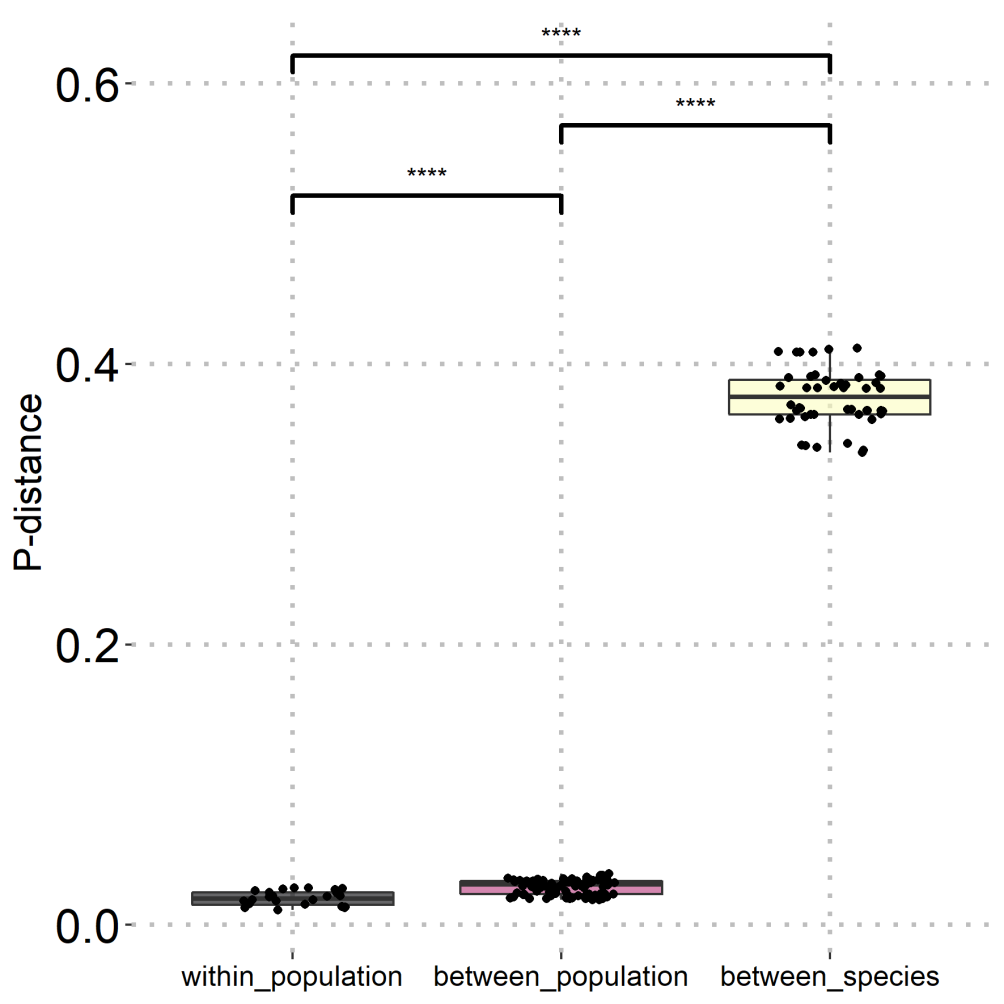

**Fig. S29 P-distances in inter-strain, intra-strain and inter-specific levels**

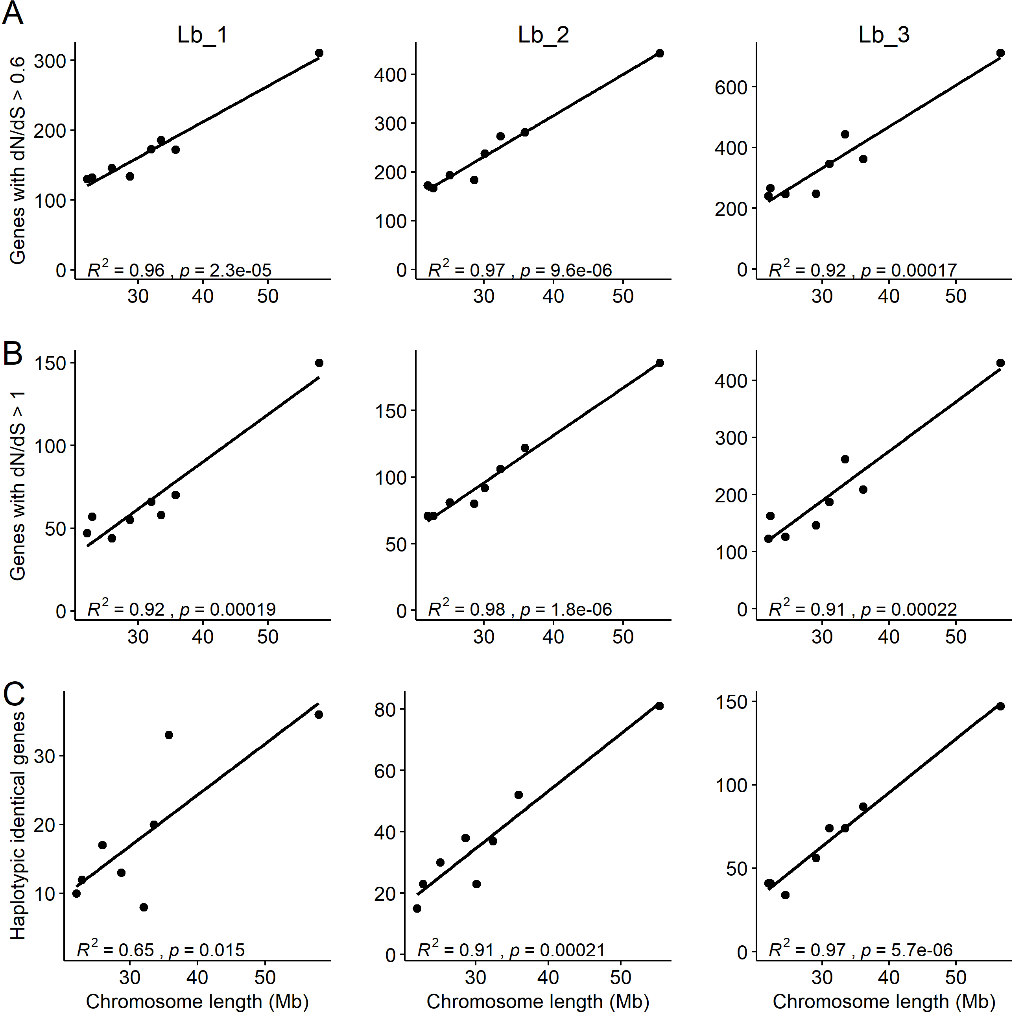

**Fig. S30 Correlation analyses between special genes (dN/dS>0.6, dN/dS>1 and identical allelic genes) and chromosome lengths.**

**Tables**

**Table S1 The sequencing output of three *Liposcelis bostrychophila* populations**

| **Population** | **Pacbio CLR data** | **DNA Illumina** | **HiC** | **RNA Illumina** |
| --- | --- | --- | --- | --- |
| ***Lb*_1** | 37,663,460,218 | 20,670,764,100 | 88,123,349,100 | 56,092,261,500 |
| ***Lb*_2** | 56,396,813,535 | 21,882,138,000 | 55,529,262,600 | 14,392,562,700 |
| ***Lb*_3** | 27,911,976,065 | 29,305,834,200 | 59,982,409,500 | 17,109,310,800 |

**Table S2 Structural and functional annotation of three strains of *Liposcelis bostrychophila* and *Liposcelis brunnea***

|  | **Lb_1** | **Lb_2** | **Lb_3** | ***Liposcelis brunnea*** |
| --- | --- | --- | --- | --- |
| **Gene numbers** | 34,469 | 34,989 | 34,630 | 15,543 |
| **BUSCO completeness** | 97.9% | 97.1% | 97.5% | 97.2% |
| **Interproscan hits** | 22,903 | 22,597 | 22,423 | 10,724 |
| **eggNOG-mapper hits** | 20,750 | 20,632 | 20,698 | 10,097 |
| **nr hits** | 26,004 | 25,881 | 26,034 | 12,175 |

**Table S3 Diploid chromosomes of three *Liposcelis bostrychophila* strains.**

|  | ***Lb*_1** | | ***Lb*_2** | | ***Lb*_3** | |
| --- | --- | --- | --- | --- | --- | --- |
| **Diploid size (Mb)** | 517.6 | | 502.0 | | 509.4 | |
| **Haplotype** | *Lb*_1A | *Lb*_1B | *Lb*_2A | *Lb*_2B | *Lb*_3A | *Lb*_3B |
| **Chromosome 1 (Mb)** | 32.0 | 30.8 | 30.1 | 31.1 | 31.1 | 30.8 |
| **Chromosome 2 (Mb)** | 33.5 | 34.4 | 32.4 | 32.6 | 33.4 | 33.9 |
| **Chromosome 3 (Mb)** | 25.9 | 27.0 | 25.1 | 25.1 | 24.5 | 24.5 |
| **Chromosome 4 (Mb)** | 57.9 | 56.4 | 55.3 | 55.3 | 56.6 | 56.3 |
| **Chromosome 5 (Mb)** | 35.7 | 36.6 | 35.9 | 35.2 | 36.1 | 36.5 |
| **Chromosome 6 (Mb)** | 22.9 | 21.4 | 22.7 | 21.3 | 21.9 | 21.6 |
| **Chromosome 7 (Mb)** | 28.7 | 29.8 | 28.6 | 27.7 | 29.1 | 28.6 |
| **Chromosome 8 (Mb)** | 22.1 | 22.4 | 21.9 | 21.5 | 22.2 | 22.4 |

**Table S4 Statistics of genes/gene trees**

|  | **Ortholog Gene numbers** |
| --- | --- |
| **chr1** | 792 |
| **chr2** | 923 |
| **chr3** | 611 |
| **chr4** | 1,736 |
| **chr5** | 1,117 |
| **chr6** | 575 |
| **chr7** | 741 |
| **chr8** | 534 |

**Table S5 824 tree topologies from 7,029 individual genes.**

| **Tree** | **Frequency** | **Topology** | **Counts** | **Frequency** |
| --- | --- | --- | --- | --- |
| TREE 1 | 0.038270024 | (Lbru,((LB1B,LB1A),((LB2A,LB2B),(LB3B,LB3A)))); | 269 | 3.83% |
| TREE 2 | 0.026177266 | (Lbru,(LB1A,(LB1B,((LB2A,LB2B),(LB3B,LB3A))))); | 184 | 2.62% |
| TREE 3 | 0.025039124 | (Lbru,((LB1B,LB1A),(LB2B,(LB2A,(LB3B,LB3A))))); | 176 | 2.50% |
| TREE 4 | 0.024754588 | (Lbru,((LB1B,LB1A),(LB2A,(LB2B,(LB3B,LB3A))))); | 174 | 2.48% |
| TREE 5 | 0.019775217 | (Lbru,(LB1B,(LB1A,((LB2A,LB2B),(LB3B,LB3A))))); | 139 | 1.98% |
| TREE 6 | 0.017498933 | (Lbru,(LB1A,(LB1B,(LB2A,(LB2B,(LB3B,LB3A)))))); | 123 | 1.75% |
| TREE 7 | 0.017214397 | (Lbru,(LB1B,(LB1A,(LB2A,(LB2B,(LB3B,LB3A)))))); | 121 | 1.72% |
| TREE 8 | 0.016360791 | (Lbru,(((LB2A,LB2B),(LB1B,LB1A)),(LB3B,LB3A))); | 115 | 1.64% |
| TREE 9 | 0.015649452 | (Lbru,(LB1A,(LB1B,(LB2B,(LB2A,(LB3B,LB3A)))))); | 110 | 1.56% |
| TREE 10 | 0.014653578 | (Lbru,(LB1B,(LB1A,(LB2B,(LB2A,(LB3B,LB3A)))))); | 103 | 1.47% |
| TREE 11 | 0.012946365 | (Lbru,(LB2A,((LB1B,LB1A),(LB2B,(LB3B,LB3A))))); | 91 | 1.29% |
| TREE 12 | 0.012804097 | (Lbru,(LB2B,((LB1B,LB1A),(LB2A,(LB3B,LB3A))))); | 90 | 1.28% |
| TREE 13 | 0.011808223 | (Lbru,((LB1B,LB1A),(LB2B,(LB3A,(LB3B,LB2A))))); | 83 | 1.18% |
| TREE 14 | 0.010954617 | (Lbru,((LB2A,LB2B),((LB1B,LB1A),(LB3B,LB3A)))); | 77 | 1.10% |
| TREE 15 | 0.010527813 | (Lbru,(LB2B,(LB2A,((LB1B,LB1A),(LB3B,LB3A))))); | 74 | 1.05% |
| TREE 16 | 0.010385546 | (Lbru,(LB2B,((LB2A,(LB1B,LB1A)),(LB3B,LB3A)))); | 73 | 1.04% |
| TREE 17 | 0.010385546 | (Lbru,(LB2A,((LB2B,(LB1B,LB1A)),(LB3B,LB3A)))); | 73 | 1.04% |
| TREE 18 | 0.009958742 | (Lbru,((LB1B,LB1A),(LB2B,(LB3B,(LB3A,LB2A))))); | 70 | 1.00% |
| TREE 19 | 0.009816475 | (Lbru,((LB2A,(LB2B,(LB1B,LB1A))),(LB3B,LB3A))); | 69 | 0.98% |
| TREE 20 | 0.009674207 | (Lbru,((LB2B,(LB2A,(LB1B,LB1A))),(LB3B,LB3A))); | 68 | 0.97% |
| TREE 21 | 0.008962868 | (Lbru,(LB2A,(LB2B,((LB1B,LB1A),(LB3B,LB3A))))); | 63 | 0.90% |
| TREE 22 | 0.008109262 | (Lbru,((LB1B,LB1A),(LB2A,(LB3A,(LB3B,LB2B))))); | 57 | 0.81% |
| TREE 23 | 0.006828852 | (Lbru,(LB2B,((LB1B,LB1A),(LB3A,(LB3B,LB2A))))); | 48 | 0.68% |
| TREE 24 | 0.006686584 | (Lbru,((LB2A,(LB1B,LB1A)),(LB2B,(LB3B,LB3A)))); | 47 | 0.67% |
| TREE 25 | 0.006686584 | (Lbru,((LB2B,(LB1B,LB1A)),(LB2A,(LB3B,LB3A)))); | 47 | 0.67% |
| TREE 26 | 0.005975245 | (Lbru,(LB3B,(LB3A,((LB2A,LB2B),(LB1B,LB1A))))); | 42 | 0.60% |
| TREE 27 | 0.005548442 | (Lbru,((LB1B,LB1A),(LB3A,(LB3B,(LB2A,LB2B))))); | 39 | 0.55% |
| TREE 28 | 0.005548442 | (Lbru,(LB1B,(LB1A,(LB2B,(LB3B,(LB3A,LB2A)))))); | 39 | 0.55% |
| TREE 29 | 0.005406174 | (Lbru,(LB1A,(LB1B,(LB2A,(LB3A,(LB3B,LB2B)))))); | 38 | 0.54% |
| TREE 30 | 0.005263907 | (Lbru,(LB1B,(LB1A,(LB2B,(LB3A,(LB3B,LB2A)))))); | 37 | 0.53% |
| TREE 31 | 0.005121639 | (Lbru,(LB1A,(LB1B,(LB2B,(LB3A,(LB3B,LB2A)))))); | 36 | 0.51% |
| TREE 32 | 0.004837103 | (Lbru,(LB3A,(LB3B,((LB2A,LB2B),(LB1B,LB1A))))); | 34 | 0.48% |
| TREE 33 | 0.004694836 | (Lbru,(LB3B,(LB3A,(LB2B,(LB2A,(LB1B,LB1A)))))); | 33 | 0.47% |
| TREE 34 | 0.004552568 | (Lbru,((LB1B,LB1A),(LB3B,(LB3A,(LB2A,LB2B))))); | 32 | 0.46% |
| TREE 35 | 0.004552568 | (Lbru,(LB1A,(LB1B,(LB3B,(LB3A,(LB2A,LB2B)))))); | 32 | 0.46% |
| TREE 36 | 0.004552568 | (Lbru,(LB2A,((LB1B,LB1A),(LB3A,(LB3B,LB2B))))); | 32 | 0.46% |
| TREE 37 | 0.004552568 | (Lbru,((LB1B,LB1A),(LB2A,(LB3B,(LB3A,LB2B))))); | 32 | 0.46% |
| TREE 38 | 0.004552568 | (Lbru,((LB1B,LB1A),(LB3A,(LB2B,(LB3B,LB2A))))); | 32 | 0.46% |
| TREE 39 | 0.0044103 | (Lbru,(LB1B,(LB1A,(LB3B,(LB3A,(LB2A,LB2B)))))); | 31 | 0.44% |
| TREE 40 | 0.004268032 | (Lbru,(LB3A,(LB3B,(LB2A,(LB2B,(LB1B,LB1A)))))); | 30 | 0.43% |
| TREE 41 | 0.004268032 | (Lbru,((LB1B,LB1A),(LB3B,(LB2B,(LB3A,LB2A))))); | 30 | 0.43% |
| TREE 42 | 0.004125765 | (Lbru,(LB1B,(LB1A,(LB2A,(LB3A,(LB3B,LB2B)))))); | 29 | 0.41% |
| TREE 43 | 0.003983497 | (Lbru,((LB1B,(LB2A,(LB1A,LB2B))),(LB3B,LB3A))); | 28 | 0.40% |
| TREE 44 | 0.003841229 | (Lbru,(LB1A,(LB1B,(LB2B,(LB3B,(LB3A,LB2A)))))); | 27 | 0.38% |
| TREE 45 | 0.003841229 | (Lbru,((LB1B,LB1A),((LB3B,LB2B),(LB3A,LB2A)))); | 27 | 0.38% |
| TREE 46 | 0.003556694 | (Lbru,(LB3B,((LB1B,LB1A),(LB3A,(LB2A,LB2B))))); | 25 | 0.36% |
| TREE 47 | 0.003556694 | (Lbru,((LB1B,LB1A),((LB3B,LB2A),(LB3A,LB2B)))); | 25 | 0.36% |
| TREE 48 | 0.003556694 | (Lbru,((LB1B,LB1A),(LB3A,(LB2A,(LB3B,LB2B))))); | 25 | 0.36% |
| TREE 49 | 0.003414426 | (Lbru,(LB1A,(LB1B,(LB2A,(LB3B,(LB3A,LB2B)))))); | 24 | 0.34% |
| TREE 50 | 0.003414426 | (Lbru,(LB1A,(LB2B,(LB1B,(LB2A,(LB3B,LB3A)))))); | 24 | 0.34% |
| TREE 51 | 0.003414426 | (Lbru,(LB2B,((LB1B,LB1A),(LB3B,(LB3A,LB2A))))); | 24 | 0.34% |
| TREE 52 | 0.003272158 | (Lbru,(LB1B,((LB1A,(LB2A,LB2B)),(LB3B,LB3A)))); | 23 | 0.33% |
| TREE 53 | 0.003272158 | (Lbru,(LB1A,(LB2A,(LB2B,(LB3A,(LB1B,LB3B)))))); | 23 | 0.33% |
| TREE 54 | 0.003272158 | (Lbru,(LB3B,(LB3A,(LB2A,(LB2B,(LB1B,LB1A)))))); | 23 | 0.33% |
| TREE 55 | 0.00312989 | (Lbru,(LB1A,((LB2A,LB2B),(LB1B,(LB3B,LB3A))))); | 22 | 0.31% |
| TREE 56 | 0.00312989 | (Lbru,(LB1A,(LB1B,(LB3A,(LB3B,(LB2A,LB2B)))))); | 22 | 0.31% |
| TREE 57 | 0.002987623 | (Lbru,(LB1B,(LB2A,(LB2B,(LB3A,(LB1A,LB3B)))))); | 21 | 0.30% |
| TREE 58 | 0.002987623 | (Lbru,(LB3B,((LB2B,(LB1B,LB1A)),(LB3A,LB2A)))); | 21 | 0.30% |
| TREE 59 | 0.002987623 | (Lbru,(LB3B,((LB1B,LB1A),(LB2B,(LB3A,LB2A))))); | 21 | 0.30% |
| TREE 60 | 0.002987623 | (Lbru,(LB1A,((LB1B,(LB2A,LB2B)),(LB3B,LB3A)))); | 21 | 0.30% |
| TREE 61 | 0.002845355 | (Lbru,(LB1B,((LB2A,LB2B),(LB1A,(LB3B,LB3A))))); | 20 | 0.28% |
| TREE 62 | 0.002845355 | (Lbru,((LB1A,(LB1B,(LB2A,LB2B))),(LB3B,LB3A))); | 20 | 0.28% |
| TREE 63 | 0.002845355 | (Lbru,(LB2B,(LB3A,((LB1B,LB1A),(LB3B,LB2A))))); | 20 | 0.28% |
| TREE 64 | 0.002845355 | (Lbru,(LB1B,(LB1A,(LB3B,(LB2A,(LB3A,LB2B)))))); | 20 | 0.28% |
| TREE 65 | 0.002845355 | (Lbru,(LB1B,(LB1A,(LB2A,(LB3B,(LB3A,LB2B)))))); | 20 | 0.28% |
| TREE 66 | 0.002845355 | (Lbru,(LB3A,((LB2B,(LB1B,LB1A)),(LB3B,LB2A)))); | 20 | 0.28% |
| TREE 67 | 0.002703087 | (Lbru,(LB1A,(LB1B,(LB3B,(LB2B,(LB3A,LB2A)))))); | 19 | 0.27% |
| TREE 68 | 0.002703087 | (Lbru,(LB3A,(LB3B,(LB2B,(LB2A,(LB1B,LB1A)))))); | 19 | 0.27% |
| TREE 69 | 0.002703087 | (Lbru,(LB1B,(LB2A,(LB1A,(LB2B,(LB3B,LB3A)))))); | 19 | 0.27% |
| TREE 70 | 0.002703087 | (Lbru,((LB1A,(LB2A,LB2B)),(LB1B,(LB3B,LB3A)))); | 19 | 0.27% |
| TREE 71 | 0.002560819 | (Lbru,((LB1B,LB1A),(LB3B,(LB2A,(LB3A,LB2B))))); | 18 | 0.26% |
| TREE 72 | 0.002560819 | (Lbru,(LB1A,(LB1B,(LB3B,(LB2A,(LB3A,LB2B)))))); | 18 | 0.26% |
| TREE 73 | 0.002560819 | (Lbru,(LB2A,((LB3B,LB2B),(LB3A,(LB1B,LB1A))))); | 18 | 0.26% |
| TREE 74 | 0.002560819 | (Lbru,(LB1A,(LB1B,((LB3B,LB2A),(LB3A,LB2B))))); | 18 | 0.26% |
| TREE 75 | 0.002560819 | (Lbru,(LB3A,((LB1B,LB1A),(LB2B,(LB3B,LB2A))))); | 18 | 0.26% |
| TREE 76 | 0.002418552 | (Lbru,(LB1B,(LB1A,((LB3B,LB2A),(LB3A,LB2B))))); | 17 | 0.24% |
| TREE 77 | 0.002418552 | (Lbru,(LB2A,(LB1B,(LB2B,(LB3A,(LB1A,LB3B)))))); | 17 | 0.24% |
| TREE 78 | 0.002418552 | (Lbru,(LB1A,((LB1B,LB2B),(LB2A,(LB3B,LB3A))))); | 17 | 0.24% |
| TREE 79 | 0.002418552 | (Lbru,(LB1B,((LB2A,(LB1A,LB2B)),(LB3B,LB3A)))); | 17 | 0.24% |
| TREE 80 | 0.002418552 | (Lbru,((LB2A,(LB1B,LB1A)),(LB3A,(LB3B,LB2B)))); | 17 | 0.24% |
| TREE 81 | 0.002418552 | (Lbru,((LB2A,LB2B),(LB1B,(LB1A,(LB3B,LB3A))))); | 17 | 0.24% |
| TREE 82 | 0.002418552 | (Lbru,((LB2B,(LB1B,LB1A)),(LB3A,(LB3B,LB2A)))); | 17 | 0.24% |
| TREE 83 | 0.002418552 | (Lbru,(LB1A,(LB1B,((LB3B,LB2B),(LB3A,LB2A))))); | 17 | 0.24% |
| TREE 84 | 0.002276284 | (Lbru,(LB2B,(LB3B,(LB3A,(LB2A,(LB1B,LB1A)))))); | 16 | 0.23% |
| TREE 85 | 0.002276284 | (Lbru,(LB2A,((LB1B,LB1A),(LB3B,(LB3A,LB2B))))); | 16 | 0.23% |
| TREE 86 | 0.002276284 | (Lbru,(LB3A,((LB1B,LB1A),(LB3B,(LB2A,LB2B))))); | 16 | 0.23% |
| TREE 87 | 0.002276284 | (Lbru,(LB3A,((LB2A,LB2B),(LB3B,(LB1B,LB1A))))); | 16 | 0.23% |
| TREE 88 | 0.002276284 | (Lbru,(LB1A,(LB1B,(LB3A,(LB2B,(LB3B,LB2A)))))); | 16 | 0.23% |
| TREE 89 | 0.002276284 | (Lbru,(LB1B,(LB1A,(LB3A,(LB2B,(LB3B,LB2A)))))); | 16 | 0.23% |
| TREE 90 | 0.002134016 | (Lbru,(LB2A,((LB1B,(LB1A,LB2B)),(LB3B,LB3A)))); | 15 | 0.21% |
| TREE 91 | 0.002134016 | (Lbru,((LB1B,(LB1A,LB2B)),(LB3A,(LB3B,LB2A)))); | 15 | 0.21% |
| TREE 92 | 0.002134016 | (Lbru,(LB1B,((LB1A,LB2A),(LB2B,(LB3B,LB3A))))); | 15 | 0.21% |
| TREE 93 | 0.002134016 | (Lbru,((LB2A,(LB1B,(LB1A,LB2B))),(LB3B,LB3A))); | 15 | 0.21% |
| TREE 94 | 0.002134016 | (Lbru,((LB1B,(LB2B,(LB1A,LB2A))),(LB3B,LB3A))); | 15 | 0.21% |
| TREE 95 | 0.001991748 | (Lbru,(LB1B,(LB1A,(LB3B,(LB2B,(LB3A,LB2A)))))); | 14 | 0.20% |
| TREE 96 | 0.001991748 | (Lbru,(LB1A,(LB2A,((LB1B,LB2B),(LB3B,LB3A))))); | 14 | 0.20% |
| TREE 97 | 0.001991748 | (Lbru,(LB1A,(LB2A,(LB1B,(LB2B,(LB3B,LB3A)))))); | 14 | 0.20% |
| TREE 98 | 0.001991748 | (Lbru,(LB1A,(LB1B,(LB3A,(LB2A,(LB3B,LB2B)))))); | 14 | 0.20% |
| TREE 99 | 0.001991748 | (Lbru,(LB2B,(LB2A,(LB1A,(LB1B,(LB3B,LB3A)))))); | 14 | 0.20% |
| TREE 100 | 0.001991748 | (Lbru,(LB1B,((LB1A,LB2B),(LB2A,(LB3B,LB3A))))); | 14 | 0.20% |
| TREE 101 | 0.001991748 | (Lbru,(LB1B,(LB2A,(LB2B,(LB3B,(LB1A,LB3A)))))); | 14 | 0.20% |
| TREE 102 | 0.001991748 | (Lbru,((LB2B,(LB1B,LB1A)),(LB3B,(LB3A,LB2A)))); | 14 | 0.20% |
| TREE 103 | 0.001991748 | (Lbru,(LB2A,((LB1A,LB2B),(LB1B,(LB3B,LB3A))))); | 14 | 0.20% |
| TREE 104 | 0.001991748 | (Lbru,(LB2A,(LB1B,(LB2B,(LB3B,(LB1A,LB3A)))))); | 14 | 0.20% |
| TREE 105 | 0.001849481 | (Lbru,(LB2A,((LB3B,(LB1B,LB1A)),(LB3A,LB2B)))); | 13 | 0.18% |
| TREE 106 | 0.001849481 | (Lbru,((LB1A,(LB1B,LB2A)),(LB2B,(LB3B,LB3A)))); | 13 | 0.18% |
| TREE 107 | 0.001849481 | (Lbru,((LB1A,(LB2A,(LB1B,LB2B))),(LB3B,LB3A))); | 13 | 0.18% |
| TREE 108 | 0.001849481 | (Lbru,((LB1B,(LB1A,LB2B)),(LB2A,(LB3B,LB3A)))); | 13 | 0.18% |
| TREE 109 | 0.001849481 | (Lbru,(LB2B,((LB3B,LB2A),(LB3A,(LB1B,LB1A))))); | 13 | 0.18% |
| TREE 110 | 0.001849481 | (Lbru,(LB1B,((LB2A,LB2B),(LB3A,(LB1A,LB3B))))); | 13 | 0.18% |
| TREE 111 | 0.001849481 | (Lbru,((LB2A,(LB1B,LB1A)),(LB3B,(LB3A,LB2B)))); | 13 | 0.18% |
| TREE 112 | 0.001849481 | (Lbru,((LB2B,(LB1B,(LB1A,LB2A))),(LB3B,LB3A))); | 13 | 0.18% |
| TREE 113 | 0.001849481 | (Lbru,(LB2B,(LB1A,(LB1B,(LB2A,(LB3B,LB3A)))))); | 13 | 0.18% |
| TREE 114 | 0.001849481 | (Lbru,(LB1A,((LB1B,LB2A),(LB2B,(LB3B,LB3A))))); | 13 | 0.18% |
| TREE 115 | 0.001849481 | (Lbru,(LB2B,(LB3A,(LB3B,(LB2A,(LB1B,LB1A)))))); | 13 | 0.18% |
| TREE 116 | 0.001849481 | (Lbru,(LB2B,(LB1B,(LB2A,(LB3A,(LB1A,LB3B)))))); | 13 | 0.18% |
| TREE 117 | 0.001849481 | (Lbru,(LB1A,(LB2A,(LB2B,(LB3B,(LB1B,LB3A)))))); | 13 | 0.18% |
| TREE 118 | 0.001707213 | (Lbru,(LB1A,(LB2B,((LB1B,LB2A),(LB3B,LB3A))))); | 12 | 0.17% |
| TREE 119 | 0.001707213 | (Lbru,(LB2B,(LB2A,(LB1B,(LB3A,(LB1A,LB3B)))))); | 12 | 0.17% |
| TREE 120 | 0.001707213 | (Lbru,(LB3B,(LB2A,(LB2B,(LB3A,(LB1B,LB1A)))))); | 12 | 0.17% |
| TREE 121 | 0.001707213 | (Lbru,((LB3B,(LB2A,(LB1B,LB1A))),(LB3A,LB2B))); | 12 | 0.17% |
| TREE 122 | 0.001707213 | (Lbru,((LB2A,LB2B),(LB1B,(LB3A,(LB1A,LB3B))))); | 12 | 0.17% |
| TREE 123 | 0.001707213 | (Lbru,((LB3B,(LB2A,LB2B)),(LB3A,(LB1B,LB1A)))); | 12 | 0.17% |
| TREE 124 | 0.001707213 | (Lbru,(LB3A,(LB2B,(LB2A,(LB3B,(LB1B,LB1A)))))); | 12 | 0.17% |
| TREE 125 | 0.001707213 | (Lbru,(LB2B,(LB1A,(LB1B,(LB3A,(LB3B,LB2A)))))); | 12 | 0.17% |
| TREE 126 | 0.001707213 | (Lbru,(LB1B,((LB1A,LB2A),(LB3A,(LB3B,LB2B))))); | 12 | 0.17% |
| TREE 127 | 0.001707213 | (Lbru,(LB3B,((LB2A,LB2B),(LB3A,(LB1B,LB1A))))); | 12 | 0.17% |
| TREE 128 | 0.001707213 | (Lbru,((LB2A,(LB1A,(LB1B,LB2B))),(LB3B,LB3A))); | 12 | 0.17% |
| TREE 129 | 0.001707213 | (Lbru,((LB2A,LB2B),(LB1B,(LB3B,(LB1A,LB3A))))); | 12 | 0.17% |
| TREE 130 | 0.001707213 | (Lbru,((LB1B,(LB2A,LB2B)),(LB1A,(LB3B,LB3A)))); | 12 | 0.17% |
| TREE 131 | 0.001564945 | (Lbru,(LB1A,((LB2A,(LB1B,LB2B)),(LB3B,LB3A)))); | 11 | 0.16% |
| TREE 132 | 0.001564945 | (Lbru,(((LB1B,LB1A),(LB3B,LB2B)),(LB3A,LB2A))); | 11 | 0.16% |
| TREE 133 | 0.001564945 | (Lbru,(LB2A,(LB2B,(LB3A,(LB3B,(LB1B,LB1A)))))); | 11 | 0.16% |
| TREE 134 | 0.001564945 | (Lbru,((LB1B,(LB1A,LB2A)),(LB3A,(LB3B,LB2B)))); | 11 | 0.16% |
| TREE 135 | 0.001564945 | (Lbru,((LB3B,(LB1B,LB1A)),(LB2B,(LB3A,LB2A)))); | 11 | 0.16% |
| TREE 136 | 0.001564945 | (Lbru,((LB1A,(LB1B,LB2B)),(LB2A,(LB3B,LB3A)))); | 11 | 0.16% |
| TREE 137 | 0.001564945 | (Lbru,((LB2A,LB2B),(LB3B,(LB3A,(LB1B,LB1A))))); | 11 | 0.16% |
| TREE 138 | 0.001564945 | (Lbru,(LB1B,((LB2B,(LB1A,LB2A)),(LB3B,LB3A)))); | 11 | 0.16% |
| TREE 139 | 0.001564945 | (Lbru,(LB3A,((LB2A,(LB1B,LB1A)),(LB3B,LB2B)))); | 11 | 0.16% |
| TREE 140 | 0.001564945 | (Lbru,(LB2B,(LB1B,(LB1A,(LB2A,(LB3B,LB3A)))))); | 11 | 0.16% |
| TREE 141 | 0.001564945 | (Lbru,(LB2A,(LB3A,(LB3B,(LB2B,(LB1B,LB1A)))))); | 11 | 0.16% |
| TREE 142 | 0.001564945 | (Lbru,(LB1A,(LB2B,(LB2A,(LB1B,(LB3B,LB3A)))))); | 11 | 0.16% |
| TREE 143 | 0.001564945 | (Lbru,(LB1B,(LB2A,(LB3B,(LB2B,(LB1A,LB3A)))))); | 11 | 0.16% |
| TREE 144 | 0.001564945 | (Lbru,(LB1B,(LB2A,((LB1A,LB2B),(LB3B,LB3A))))); | 11 | 0.16% |
| TREE 145 | 0.001564945 | (Lbru,(LB3B,((LB1B,LB1A),(LB2A,(LB3A,LB2B))))); | 11 | 0.16% |
| TREE 146 | 0.001422677 | (Lbru,(LB3B,(LB2B,((LB1B,LB1A),(LB3A,LB2A))))); | 10 | 0.14% |
| TREE 147 | 0.001422677 | (Lbru,(LB2B,((LB1B,(LB1A,LB2A)),(LB3B,LB3A)))); | 10 | 0.14% |
| TREE 148 | 0.001422677 | (Lbru,(LB1A,(LB2B,(LB2A,(LB3A,(LB1B,LB3B)))))); | 10 | 0.14% |
| TREE 149 | 0.001422677 | (Lbru,((LB3B,(LB1B,LB1A)),(LB3A,(LB2A,LB2B)))); | 10 | 0.14% |
| TREE 150 | 0.001422677 | (Lbru,(LB2A,(LB2B,(LB1A,(LB1B,(LB3B,LB3A)))))); | 10 | 0.14% |
| TREE 151 | 0.001422677 | (Lbru,((LB2A,LB2B),(LB3A,(LB3B,(LB1B,LB1A))))); | 10 | 0.14% |
| TREE 152 | 0.001422677 | (Lbru,(LB3A,((LB1B,LB1A),(LB2A,(LB3B,LB2B))))); | 10 | 0.14% |
| TREE 153 | 0.001422677 | (Lbru,(LB2A,(LB1B,(LB1A,(LB2B,(LB3B,LB3A)))))); | 10 | 0.14% |
| TREE 154 | 0.001422677 | (Lbru,(LB1A,((LB1B,LB3B),(LB3A,(LB2A,LB2B))))); | 10 | 0.14% |
| TREE 155 | 0.001422677 | (Lbru,(LB1B,(LB1A,(LB3A,(LB3B,(LB2A,LB2B)))))); | 10 | 0.14% |
| TREE 156 | 0.001422677 | (Lbru,(LB2A,(LB1A,(LB3A,(LB3B,(LB1B,LB2B)))))); | 10 | 0.14% |
| TREE 157 | 0.001422677 | (Lbru,((LB2A,LB2B),(LB1A,(LB1B,(LB3B,LB3A))))); | 10 | 0.14% |
| TREE 158 | 0.001422677 | (Lbru,(LB1B,(LB2B,((LB1A,LB2A),(LB3B,LB3A))))); | 10 | 0.14% |
| TREE 159 | 0.001422677 | (Lbru,(((LB1B,LB2B),(LB1A,LB2A)),(LB3B,LB3A))); | 10 | 0.14% |
| TREE 160 | 0.001422677 | (Lbru,(LB2A,(LB3B,(LB3A,(LB2B,(LB1B,LB1A)))))); | 10 | 0.14% |
| TREE 161 | 0.001422677 | (Lbru,(LB1B,((LB1A,LB2B),(LB3A,(LB3B,LB2A))))); | 10 | 0.14% |
| TREE 162 | 0.001422677 | (Lbru,(LB3B,(LB3A,(LB1B,(LB2A,(LB1A,LB2B)))))); | 10 | 0.14% |
| TREE 163 | 0.001422677 | (Lbru,(LB2B,(LB2A,(LB3B,(LB3A,(LB1B,LB1A)))))); | 10 | 0.14% |
| TREE 164 | 0.001422677 | (Lbru,(LB1B,(LB2B,(LB1A,(LB2A,(LB3B,LB3A)))))); | 10 | 0.14% |
| TREE 165 | 0.001422677 | (Lbru,(LB1B,(LB2A,(LB3B,(LB3A,(LB1A,LB2B)))))); | 10 | 0.14% |
| TREE 166 | 0.00128041 | (Lbru,(LB2B,(LB3B,((LB1B,LB1A),(LB3A,LB2A))))); | 9 | 0.13% |
| TREE 167 | 0.00128041 | (Lbru,(LB2A,(LB3A,((LB1B,LB1A),(LB3B,LB2B))))); | 9 | 0.13% |
| TREE 168 | 0.00128041 | (Lbru,(LB2B,(LB2A,(LB1B,(LB3B,(LB1A,LB3A)))))); | 9 | 0.13% |
| TREE 169 | 0.00128041 | (Lbru,(LB1A,((LB1B,LB2B),(LB3A,(LB3B,LB2A))))); | 9 | 0.13% |
| TREE 170 | 0.00128041 | (Lbru,(LB2A,(LB2B,(LB3B,(LB3A,(LB1B,LB1A)))))); | 9 | 0.13% |
| TREE 171 | 0.00128041 | (Lbru,(LB3A,(LB2A,(LB2B,(LB3B,(LB1B,LB1A)))))); | 9 | 0.13% |
| TREE 172 | 0.00128041 | (Lbru,((LB1B,(LB1A,LB2A)),(LB2B,(LB3B,LB3A)))); | 9 | 0.13% |
| TREE 173 | 0.00128041 | (Lbru,(LB2A,(LB1B,(LB3A,(LB3B,(LB1A,LB2B)))))); | 9 | 0.13% |
| TREE 174 | 0.00128041 | (Lbru,((LB1A,LB2A),(LB1B,(LB2B,(LB3B,LB3A))))); | 9 | 0.13% |
| TREE 175 | 0.00128041 | (Lbru,(LB1B,(LB2A,(LB3A,(LB2B,(LB1A,LB3B)))))); | 9 | 0.13% |
| TREE 176 | 0.00128041 | (Lbru,(LB1A,(LB2A,(LB2B,(LB1B,(LB3B,LB3A)))))); | 9 | 0.13% |
| TREE 177 | 0.00128041 | (Lbru,(LB1A,(LB3B,((LB2A,LB2B),(LB1B,LB3A))))); | 9 | 0.13% |
| TREE 178 | 0.00128041 | (Lbru,((LB2B,(LB1A,(LB1B,LB2A))),(LB3B,LB3A))); | 9 | 0.13% |
| TREE 179 | 0.00128041 | (Lbru,((LB1B,LB2A),(LB2B,(LB3A,(LB1A,LB3B))))); | 9 | 0.13% |
| TREE 180 | 0.00128041 | (Lbru,(LB3A,(LB2B,(LB3B,(LB2A,(LB1B,LB1A)))))); | 9 | 0.13% |
| TREE 181 | 0.00128041 | (Lbru,(LB1A,(LB2A,(LB3A,(LB3B,(LB1B,LB2B)))))); | 9 | 0.13% |
| TREE 182 | 0.00128041 | (Lbru,((LB1B,LB2B),((LB1A,LB2A),(LB3B,LB3A)))); | 9 | 0.13% |
| TREE 183 | 0.00128041 | (Lbru,(LB1A,(LB2A,(LB3A,(LB2B,(LB1B,LB3B)))))); | 9 | 0.13% |
| TREE 184 | 0.00128041 | (Lbru,(LB2A,(LB1B,((LB1A,LB2B),(LB3B,LB3A))))); | 9 | 0.13% |
| TREE 185 | 0.00128041 | (Lbru,(LB2B,(LB1A,((LB1B,LB2A),(LB3B,LB3A))))); | 9 | 0.13% |
| TREE 186 | 0.001138142 | (Lbru,(LB2B,(LB2A,(LB1B,(LB1A,(LB3B,LB3A)))))); | 8 | 0.11% |
| TREE 187 | 0.001138142 | (Lbru,(LB1B,(LB2B,(LB2A,(LB3A,(LB1A,LB3B)))))); | 8 | 0.11% |
| TREE 188 | 0.001138142 | (Lbru,((LB2A,(LB3B,(LB1A,LB2B))),(LB1B,LB3A))); | 8 | 0.11% |
| TREE 189 | 0.001138142 | (Lbru,(LB2B,(LB2A,(LB3A,(LB3B,(LB1B,LB1A)))))); | 8 | 0.11% |
| TREE 190 | 0.001138142 | (Lbru,(LB2A,(LB1B,(LB1A,(LB3A,(LB3B,LB2B)))))); | 8 | 0.11% |
| TREE 191 | 0.001138142 | (Lbru,(LB1A,(LB2B,(LB1B,(LB3B,(LB3A,LB2A)))))); | 8 | 0.11% |
| TREE 192 | 0.001138142 | (Lbru,(LB3A,(LB3B,(LB1B,(LB1A,(LB2A,LB2B)))))); | 8 | 0.11% |
| TREE 193 | 0.001138142 | (Lbru,(LB3B,(LB3A,(LB2B,(LB1B,(LB1A,LB2A)))))); | 8 | 0.11% |
| TREE 194 | 0.001138142 | (Lbru,((LB1B,LB2B),(LB1A,(LB2A,(LB3B,LB3A))))); | 8 | 0.11% |
| TREE 195 | 0.001138142 | (Lbru,(LB3A,(LB3B,(LB1B,(LB2A,(LB1A,LB2B)))))); | 8 | 0.11% |
| TREE 196 | 0.001138142 | (Lbru,((LB3B,LB2B),((LB1B,LB1A),(LB3A,LB2A)))); | 8 | 0.11% |
| TREE 197 | 0.001138142 | (Lbru,(LB3B,(LB2A,(LB3A,(LB2B,(LB1B,LB1A)))))); | 8 | 0.11% |
| TREE 198 | 0.001138142 | (Lbru,(LB1B,(LB2B,(LB1A,(LB3A,(LB3B,LB2A)))))); | 8 | 0.11% |
| TREE 199 | 0.000995874 | (Lbru,(LB2A,(LB1A,(LB2B,(LB3A,(LB1B,LB3B)))))); | 7 | 0.10% |
| TREE 200 | 0.000995874 | (Lbru,(LB1B,(LB2B,(LB2A,(LB1A,(LB3B,LB3A)))))); | 7 | 0.10% |
| TREE 201 | 0.000995874 | (Lbru,(LB2A,(LB1A,(LB1B,(LB2B,(LB3B,LB3A)))))); | 7 | 0.10% |
| TREE 202 | 0.000995874 | (Lbru,(LB1A,(LB2A,(LB3B,(LB2B,(LB1B,LB3A)))))); | 7 | 0.10% |
| TREE 203 | 0.000995874 | (Lbru,(LB1A,(LB2B,(LB2A,(LB3B,(LB1B,LB3A)))))); | 7 | 0.10% |
| TREE 204 | 0.000995874 | (Lbru,((LB1A,(LB1B,LB2A)),(LB3A,(LB3B,LB2B)))); | 7 | 0.10% |
| TREE 205 | 0.000995874 | (Lbru,((LB1B,LB2A),(LB3B,(LB2B,(LB1A,LB3A))))); | 7 | 0.10% |
| TREE 206 | 0.000995874 | (Lbru,(LB3B,(LB2B,(LB2A,(LB3A,(LB1B,LB1A)))))); | 7 | 0.10% |
| TREE 207 | 0.000995874 | (Lbru,((LB1B,LB2A),(LB3A,(LB2B,(LB1A,LB3B))))); | 7 | 0.10% |
| TREE 208 | 0.000995874 | (Lbru,(LB2B,((LB1B,LB2A),(LB3A,(LB1A,LB3B))))); | 7 | 0.10% |
| TREE 209 | 0.000995874 | (Lbru,((LB2B,(LB1A,LB2A)),(LB1B,(LB3B,LB3A)))); | 7 | 0.10% |
| TREE 210 | 0.000995874 | (Lbru,(LB1B,(LB3A,(LB3B,(LB1A,(LB2A,LB2B)))))); | 7 | 0.10% |
| TREE 211 | 0.000995874 | (Lbru,(LB2B,((LB1B,LB2A),(LB1A,(LB3B,LB3A))))); | 7 | 0.10% |
| TREE 212 | 0.000995874 | (Lbru,(LB2B,((LB1A,LB2A),(LB1B,(LB3B,LB3A))))); | 7 | 0.10% |
| TREE 213 | 0.000995874 | (Lbru,(((LB1B,LB2A),(LB1A,LB2B)),(LB3B,LB3A))); | 7 | 0.10% |
| TREE 214 | 0.000995874 | (Lbru,(LB1A,((LB1B,LB2B),(LB3B,(LB3A,LB2A))))); | 7 | 0.10% |
| TREE 215 | 0.000995874 | (Lbru,(LB1A,((LB2A,LB2B),(LB3A,(LB1B,LB3B))))); | 7 | 0.10% |
| TREE 216 | 0.000995874 | (Lbru,(LB1B,(LB1A,(LB3A,(LB2A,(LB3B,LB2B)))))); | 7 | 0.10% |
| TREE 217 | 0.000995874 | (Lbru,(LB2B,(LB1B,(LB3A,(LB2A,(LB1A,LB3B)))))); | 7 | 0.10% |
| TREE 218 | 0.000995874 | (Lbru,(LB1B,(LB2B,(LB2A,(LB3B,(LB1A,LB3A)))))); | 7 | 0.10% |
| TREE 219 | 0.000995874 | (Lbru,(LB3A,(LB3B,((LB1B,LB2A),(LB1A,LB2B))))); | 7 | 0.10% |
| TREE 220 | 0.000995874 | (Lbru,(LB1A,(LB3B,(LB3A,(LB1B,(LB2A,LB2B)))))); | 7 | 0.10% |
| TREE 221 | 0.000995874 | (Lbru,(LB2A,((LB1A,(LB1B,LB2B)),(LB3B,LB3A)))); | 7 | 0.10% |
| TREE 222 | 0.000995874 | (Lbru,(LB2B,((LB3B,(LB1B,LB1A)),(LB3A,LB2A)))); | 7 | 0.10% |
| TREE 223 | 0.000995874 | (Lbru,(LB1B,(LB1A,((LB3B,LB2B),(LB3A,LB2A))))); | 7 | 0.10% |
| TREE 224 | 0.000853606 | (Lbru,(LB3A,((LB1B,LB2A),(LB2B,(LB1A,LB3B))))); | 6 | 0.09% |
| TREE 225 | 0.000853606 | (Lbru,(LB1B,(LB3A,(LB1A,(LB3B,(LB2A,LB2B)))))); | 6 | 0.09% |
| TREE 226 | 0.000853606 | (Lbru,(LB3A,(LB2A,((LB1B,LB1A),(LB3B,LB2B))))); | 6 | 0.09% |
| TREE 227 | 0.000853606 | (Lbru,((LB1B,(LB1A,LB2B)),(LB3B,(LB3A,LB2A)))); | 6 | 0.09% |
| TREE 228 | 0.000853606 | (Lbru,(LB3A,(LB1A,(LB1B,(LB3B,(LB2A,LB2B)))))); | 6 | 0.09% |
| TREE 229 | 0.000853606 | (Lbru,((LB2B,(LB3B,LB2A)),(LB3A,(LB1B,LB1A)))); | 6 | 0.09% |
| TREE 230 | 0.000853606 | (Lbru,(LB1A,((LB2B,(LB1B,LB3B)),(LB3A,LB2A)))); | 6 | 0.09% |
| TREE 231 | 0.000853606 | (Lbru,(LB1A,((LB3B,LB2B),(LB1B,(LB3A,LB2A))))); | 6 | 0.09% |
| TREE 232 | 0.000853606 | (Lbru,(LB1A,((LB1B,LB2A),(LB3A,(LB3B,LB2B))))); | 6 | 0.09% |
| TREE 233 | 0.000853606 | (Lbru,(LB1A,((LB2B,(LB1B,LB2A)),(LB3B,LB3A)))); | 6 | 0.09% |
| TREE 234 | 0.000853606 | (Lbru,((LB1B,(LB1A,(LB2A,LB2B))),(LB3B,LB3A))); | 6 | 0.09% |
| TREE 235 | 0.000853606 | (Lbru,((LB1B,(LB1A,LB3B)),(LB2B,(LB3A,LB2A)))); | 6 | 0.09% |
| TREE 236 | 0.000853606 | (Lbru,(LB2A,(LB1A,(LB2B,(LB3B,(LB1B,LB3A)))))); | 6 | 0.09% |
| TREE 237 | 0.000853606 | (Lbru,(LB2B,(LB3B,(LB2A,(LB3A,(LB1B,LB1A)))))); | 6 | 0.09% |
| TREE 238 | 0.000853606 | (Lbru,(LB3B,(LB2B,(LB3A,(LB2A,(LB1B,LB1A)))))); | 6 | 0.09% |
| TREE 239 | 0.000853606 | (Lbru,((LB1B,(LB2A,(LB3B,LB3A))),(LB1A,LB2B))); | 6 | 0.09% |
| TREE 240 | 0.000853606 | (Lbru,(LB3B,(LB1A,(LB2A,(LB2B,(LB1B,LB3A)))))); | 6 | 0.09% |
| TREE 241 | 0.000853606 | (Lbru,(LB3B,((LB1B,LB2A),(LB2B,(LB1A,LB3A))))); | 6 | 0.09% |
| TREE 242 | 0.000853606 | (Lbru,(LB3A,((LB1B,(LB1A,LB2A)),(LB3B,LB2B)))); | 6 | 0.09% |
| TREE 243 | 0.000853606 | (Lbru,(LB3A,(LB3B,(LB2B,(LB1B,(LB1A,LB2A)))))); | 6 | 0.09% |
| TREE 244 | 0.000853606 | (Lbru,(LB2A,(LB2B,(LB3A,(LB1A,(LB1B,LB3B)))))); | 6 | 0.09% |
| TREE 245 | 0.000853606 | (Lbru,((LB2B,(LB1B,LB2A)),(LB1A,(LB3B,LB3A)))); | 6 | 0.09% |
| TREE 246 | 0.000853606 | (Lbru,((LB1B,LB2A),(LB1A,(LB2B,(LB3B,LB3A))))); | 6 | 0.09% |
| TREE 247 | 0.000853606 | (Lbru,(LB2B,((LB2A,(LB1A,LB3B)),(LB1B,LB3A)))); | 6 | 0.09% |
| TREE 248 | 0.000853606 | (Lbru,((LB1A,LB2A),(LB3A,(LB3B,(LB1B,LB2B))))); | 6 | 0.09% |
| TREE 249 | 0.000853606 | (Lbru,(LB2B,(LB3B,(LB1B,(LB2A,(LB1A,LB3A)))))); | 6 | 0.09% |
| TREE 250 | 0.000853606 | (Lbru,(LB2A,((LB1B,LB2B),(LB3A,(LB1A,LB3B))))); | 6 | 0.09% |
| TREE 251 | 0.000853606 | (Lbru,((LB1B,LB2B),(LB2A,(LB3A,(LB1A,LB3B))))); | 6 | 0.09% |
| TREE 252 | 0.000853606 | (Lbru,((LB1A,(LB2A,LB2B)),(LB3A,(LB1B,LB3B)))); | 6 | 0.09% |
| TREE 253 | 0.000853606 | (Lbru,(((LB1B,LB1A),(LB3B,LB2A)),(LB3A,LB2B))); | 6 | 0.09% |
| TREE 254 | 0.000853606 | (Lbru,((LB3B,LB2B),(LB2A,(LB3A,(LB1B,LB1A))))); | 6 | 0.09% |
| TREE 255 | 0.000853606 | (Lbru,((LB2A,LB2B),(LB1A,(LB3A,(LB1B,LB3B))))); | 6 | 0.09% |
| TREE 256 | 0.000853606 | (Lbru,(LB1B,(LB2A,(LB3A,(LB3B,(LB1A,LB2B)))))); | 6 | 0.09% |
| TREE 257 | 0.000853606 | (Lbru,((LB1B,(LB1A,LB2A)),(LB3B,(LB3A,LB2B)))); | 6 | 0.09% |
| TREE 258 | 0.000853606 | (Lbru,(LB3A,(LB1B,(LB2B,(LB2A,(LB1A,LB3B)))))); | 6 | 0.09% |
| TREE 259 | 0.000853606 | (Lbru,((LB1A,(LB2B,(LB1B,LB2A))),(LB3B,LB3A))); | 6 | 0.09% |
| TREE 260 | 0.000853606 | (Lbru,(LB2A,(LB3B,((LB1B,LB1A),(LB3A,LB2B))))); | 6 | 0.09% |
| TREE 261 | 0.000853606 | (Lbru,(LB1A,((LB3B,LB2A),(LB2B,(LB1B,LB3A))))); | 6 | 0.09% |
| TREE 262 | 0.000853606 | (Lbru,(LB1B,((LB3B,(LB2A,LB2B)),(LB1A,LB3A)))); | 6 | 0.09% |
| TREE 263 | 0.000853606 | (Lbru,((LB2A,(LB1A,LB2B)),(LB1B,(LB3B,LB3A)))); | 6 | 0.09% |
| TREE 264 | 0.000853606 | (Lbru,(LB1B,(LB2A,(LB2B,(LB1A,(LB3B,LB3A)))))); | 6 | 0.09% |
| TREE 265 | 0.000853606 | (Lbru,((LB3B,LB2B),(LB3A,(LB2A,(LB1B,LB1A))))); | 6 | 0.09% |
| TREE 266 | 0.000853606 | (Lbru,(LB2A,(LB2B,(LB1B,(LB3B,(LB1A,LB3A)))))); | 6 | 0.09% |
| TREE 267 | 0.000853606 | (Lbru,(LB1B,(LB3B,(LB3A,(LB2B,(LB1A,LB2A)))))); | 6 | 0.09% |
| TREE 268 | 0.000853606 | (Lbru,(LB2A,(LB1B,(LB2B,(LB1A,(LB3B,LB3A)))))); | 6 | 0.09% |
| TREE 269 | 0.000711339 | (Lbru,(LB1B,(LB3B,(LB2A,(LB2B,(LB1A,LB3A)))))); | 5 | 0.07% |
| TREE 270 | 0.000711339 | (Lbru,(LB1B,((LB2A,LB2B),(LB3B,(LB1A,LB3A))))); | 5 | 0.07% |
| TREE 271 | 0.000711339 | (Lbru,(LB1B,(LB3B,(LB3A,(LB2A,(LB1A,LB2B)))))); | 5 | 0.07% |
| TREE 272 | 0.000711339 | (Lbru,((LB1A,(LB1B,LB2B)),(LB3A,(LB3B,LB2A)))); | 5 | 0.07% |
| TREE 273 | 0.000711339 | (Lbru,(LB2B,(LB1A,(LB3A,(LB2A,(LB1B,LB3B)))))); | 5 | 0.07% |
| TREE 274 | 0.000711339 | (Lbru,(LB2B,(LB1B,(LB1A,(LB3A,(LB3B,LB2A)))))); | 5 | 0.07% |
| TREE 275 | 0.000711339 | (Lbru,(LB3B,(LB2B,(LB3A,(LB1B,(LB1A,LB2A)))))); | 5 | 0.07% |
| TREE 276 | 0.000711339 | (Lbru,(LB3A,(LB1B,(LB3B,(LB2B,(LB1A,LB2A)))))); | 5 | 0.07% |
| TREE 277 | 0.000711339 | (Lbru,(LB2A,(LB2B,(LB1B,(LB3A,(LB1A,LB3B)))))); | 5 | 0.07% |
| TREE 278 | 0.000711339 | (Lbru,(LB2A,((LB2B,(LB1A,LB3B)),(LB1B,LB3A)))); | 5 | 0.07% |
| TREE 279 | 0.000711339 | (Lbru,(LB2B,(LB2A,(LB1A,(LB3A,(LB1B,LB3B)))))); | 5 | 0.07% |
| TREE 280 | 0.000711339 | (Lbru,(LB1A,((LB3B,(LB2A,LB2B)),(LB1B,LB3A)))); | 5 | 0.07% |
| TREE 281 | 0.000711339 | (Lbru,(LB1B,((LB1A,LB3B),(LB3A,(LB2A,LB2B))))); | 5 | 0.07% |
| TREE 282 | 0.000711339 | (Lbru,(LB3B,((LB2A,(LB1B,LB1A)),(LB3A,LB2B)))); | 5 | 0.07% |
| TREE 283 | 0.000711339 | (Lbru,(LB3B,(LB3A,((LB1B,LB2A),(LB1A,LB2B))))); | 5 | 0.07% |
| TREE 284 | 0.000711339 | (Lbru,(LB1A,((LB2A,LB2B),(LB3B,(LB1B,LB3A))))); | 5 | 0.07% |
| TREE 285 | 0.000711339 | (Lbru,(LB3A,(LB3B,(LB2A,(LB1B,(LB1A,LB2B)))))); | 5 | 0.07% |
| TREE 286 | 0.000711339 | (Lbru,(LB3A,(LB2A,(LB3B,(LB2B,(LB1B,LB1A)))))); | 5 | 0.07% |
| TREE 287 | 0.000711339 | (Lbru,(LB2A,(LB2B,(LB1B,(LB1A,(LB3B,LB3A)))))); | 5 | 0.07% |
| TREE 288 | 0.000711339 | (Lbru,(LB2B,(LB1A,(LB2A,(LB1B,(LB3B,LB3A)))))); | 5 | 0.07% |
| TREE 289 | 0.000711339 | (Lbru,(LB2B,(LB1A,(LB2A,(LB3A,(LB1B,LB3B)))))); | 5 | 0.07% |
| TREE 290 | 0.000711339 | (Lbru,(LB3B,(LB3A,(LB1A,(LB1B,(LB2A,LB2B)))))); | 5 | 0.07% |
| TREE 291 | 0.000711339 | (Lbru,(LB3A,((LB1B,(LB1A,LB2B)),(LB3B,LB2A)))); | 5 | 0.07% |
| TREE 292 | 0.000711339 | (Lbru,(LB1B,(LB2B,((LB1A,LB3B),(LB3A,LB2A))))); | 5 | 0.07% |
| TREE 293 | 0.000711339 | (Lbru,(LB3A,(LB3B,(LB1A,(LB1B,(LB2A,LB2B)))))); | 5 | 0.07% |
| TREE 294 | 0.000711339 | (Lbru,(LB2B,(LB1B,(LB2A,(LB3B,(LB1A,LB3A)))))); | 5 | 0.07% |
| TREE 295 | 0.000711339 | (Lbru,(LB1A,((LB1B,LB2A),(LB3B,(LB3A,LB2B))))); | 5 | 0.07% |
| TREE 296 | 0.000711339 | (Lbru,(LB3B,(LB3A,(LB1B,(LB2B,(LB1A,LB2A)))))); | 5 | 0.07% |
| TREE 297 | 0.000711339 | (Lbru,(LB1B,(LB3B,(LB2A,(LB3A,(LB1A,LB2B)))))); | 5 | 0.07% |
| TREE 298 | 0.000711339 | (Lbru,(LB1A,(LB2B,((LB1B,LB3B),(LB3A,LB2A))))); | 5 | 0.07% |
| TREE 299 | 0.000711339 | (Lbru,(LB1A,((LB3B,LB2B),(LB2A,(LB1B,LB3A))))); | 5 | 0.07% |
| TREE 300 | 0.000711339 | (Lbru,(LB3A,(LB2B,(LB3B,(LB1B,(LB1A,LB2A)))))); | 5 | 0.07% |
| TREE 301 | 0.000711339 | (Lbru,(LB1A,(LB2B,(LB1B,(LB3A,(LB3B,LB2A)))))); | 5 | 0.07% |
| TREE 302 | 0.000711339 | (Lbru,((LB2A,(LB1B,LB2B)),(LB1A,(LB3B,LB3A)))); | 5 | 0.07% |
| TREE 303 | 0.000711339 | (Lbru,(LB3A,((LB1A,(LB1B,LB2B)),(LB3B,LB2A)))); | 5 | 0.07% |
| TREE 304 | 0.000711339 | (Lbru,((LB1A,LB2A),((LB1B,LB2B),(LB3B,LB3A)))); | 5 | 0.07% |
| TREE 305 | 0.000711339 | (Lbru,((LB1B,(LB1A,LB3B)),(LB3A,(LB2A,LB2B)))); | 5 | 0.07% |
| TREE 306 | 0.000711339 | (Lbru,(LB3A,(LB1B,(LB1A,(LB2B,(LB3B,LB2A)))))); | 5 | 0.07% |
| TREE 307 | 0.000711339 | (Lbru,((LB1B,(LB2A,LB2B)),(LB3B,(LB1A,LB3A)))); | 5 | 0.07% |
| TREE 308 | 0.000711339 | (Lbru,(LB2A,(LB1B,(LB3B,(LB3A,(LB1A,LB2B)))))); | 5 | 0.07% |
| TREE 309 | 0.000711339 | (Lbru,((LB3B,(LB2B,(LB1B,LB1A))),(LB3A,LB2A))); | 5 | 0.07% |
| TREE 310 | 0.000711339 | (Lbru,(LB1A,(LB3A,(LB3B,(LB2B,(LB1B,LB2A)))))); | 5 | 0.07% |
| TREE 311 | 0.000711339 | (Lbru,((LB3B,(LB1B,LB1A)),(LB2A,(LB3A,LB2B)))); | 5 | 0.07% |
| TREE 312 | 0.000711339 | (Lbru,(LB2B,((LB1A,(LB1B,LB2A)),(LB3B,LB3A)))); | 5 | 0.07% |
| TREE 313 | 0.000711339 | (Lbru,((LB3B,(LB2A,(LB1A,LB2B))),(LB1B,LB3A))); | 5 | 0.07% |
| TREE 314 | 0.000711339 | (Lbru,(LB1B,(LB3B,(LB3A,(LB1A,(LB2A,LB2B)))))); | 5 | 0.07% |
| TREE 315 | 0.000711339 | (Lbru,(LB2A,((LB1B,LB2B),(LB3B,(LB1A,LB3A))))); | 5 | 0.07% |
| TREE 316 | 0.000711339 | (Lbru,(LB3B,((LB2A,LB2B),(LB1B,(LB1A,LB3A))))); | 5 | 0.07% |
| TREE 317 | 0.000711339 | (Lbru,((LB2A,(LB3B,LB2B)),(LB3A,(LB1B,LB1A)))); | 5 | 0.07% |
| TREE 318 | 0.000711339 | (Lbru,(LB2A,(LB1A,(LB3A,(LB1B,(LB3B,LB2B)))))); | 5 | 0.07% |
| TREE 319 | 0.000711339 | (Lbru,((LB2A,(LB1B,LB2B)),(LB3B,(LB1A,LB3A)))); | 5 | 0.07% |
| TREE 320 | 0.000711339 | (Lbru,(LB2A,((LB1A,(LB1B,LB3B)),(LB3A,LB2B)))); | 5 | 0.07% |
| TREE 321 | 0.000711339 | (Lbru,(LB1B,((LB3B,LB2A),(LB2B,(LB1A,LB3A))))); | 5 | 0.07% |
| TREE 322 | 0.000711339 | (Lbru,(LB1B,(LB3A,(LB3B,(LB2B,(LB1A,LB2A)))))); | 5 | 0.07% |
| TREE 323 | 0.000569071 | (Lbru,(LB3B,(LB1A,(LB1B,(LB2B,(LB3A,LB2A)))))); | 4 | 0.06% |
| TREE 324 | 0.000569071 | (Lbru,(LB3B,(LB1A,(LB1B,(LB2A,(LB3A,LB2B)))))); | 4 | 0.06% |
| TREE 325 | 0.000569071 | (Lbru,(LB1B,(LB3B,(LB2B,(LB2A,(LB1A,LB3A)))))); | 4 | 0.06% |
| TREE 326 | 0.000569071 | (Lbru,(LB1A,(LB3B,(LB1B,(LB3A,(LB2A,LB2B)))))); | 4 | 0.06% |
| TREE 327 | 0.000569071 | (Lbru,(LB2A,(LB3B,(LB1B,(LB1A,(LB3A,LB2B)))))); | 4 | 0.06% |
| TREE 328 | 0.000569071 | (Lbru,(LB1A,(LB3A,(LB3B,(LB1B,(LB2A,LB2B)))))); | 4 | 0.06% |
| TREE 329 | 0.000569071 | (Lbru,(LB2B,((LB1A,LB2A),(LB3A,(LB1B,LB3B))))); | 4 | 0.06% |
| TREE 330 | 0.000569071 | (Lbru,(LB3B,((LB2A,LB2B),(LB1A,(LB1B,LB3A))))); | 4 | 0.06% |
| TREE 331 | 0.000569071 | (Lbru,(LB3B,(LB2A,((LB1B,LB1A),(LB3A,LB2B))))); | 4 | 0.06% |
| TREE 332 | 0.000569071 | (Lbru,(LB2A,(LB2B,(LB1A,(LB3A,(LB1B,LB3B)))))); | 4 | 0.06% |
| TREE 333 | 0.000569071 | (Lbru,(LB3B,(LB2A,(LB1B,(LB3A,(LB1A,LB2B)))))); | 4 | 0.06% |
| TREE 334 | 0.000569071 | (Lbru,(LB1A,(LB2B,(LB3A,(LB2A,(LB1B,LB3B)))))); | 4 | 0.06% |
| TREE 335 | 0.000569071 | (Lbru,(LB2A,(LB1B,(LB3B,(LB2B,(LB1A,LB3A)))))); | 4 | 0.06% |
| TREE 336 | 0.000569071 | (Lbru,(LB3A,(LB1B,(LB2A,(LB3B,(LB1A,LB2B)))))); | 4 | 0.06% |
| TREE 337 | 0.000569071 | (Lbru,((LB1A,LB2A),(LB3B,(LB3A,(LB1B,LB2B))))); | 4 | 0.06% |
| TREE 338 | 0.000569071 | (Lbru,(LB2A,(LB1B,(LB3A,(LB2B,(LB1A,LB3B)))))); | 4 | 0.06% |
| TREE 339 | 0.000569071 | (Lbru,(LB3B,((LB1A,LB2A),(LB3A,(LB1B,LB2B))))); | 4 | 0.06% |
| TREE 340 | 0.000569071 | (Lbru,(LB2B,(LB3A,(LB2A,(LB3B,(LB1B,LB1A)))))); | 4 | 0.06% |
| TREE 341 | 0.000569071 | (Lbru,(LB1A,((LB2B,(LB3B,LB2A)),(LB1B,LB3A)))); | 4 | 0.06% |
| TREE 342 | 0.000569071 | (Lbru,((LB2A,LB2B),(LB3A,(LB1B,(LB1A,LB3B))))); | 4 | 0.06% |
| TREE 343 | 0.000569071 | (Lbru,(LB2A,(LB1A,(LB1B,(LB3A,(LB3B,LB2B)))))); | 4 | 0.06% |
| TREE 344 | 0.000569071 | (Lbru,(LB1B,(LB3A,(LB3B,(LB2A,(LB1A,LB2B)))))); | 4 | 0.06% |
| TREE 345 | 0.000569071 | (Lbru,(LB2B,(LB1B,((LB1A,LB2A),(LB3B,LB3A))))); | 4 | 0.06% |
| TREE 346 | 0.000569071 | (Lbru,((LB1A,LB2A),(LB1B,(LB3A,(LB3B,LB2B))))); | 4 | 0.06% |
| TREE 347 | 0.000569071 | (Lbru,(LB2B,((LB2A,(LB1B,LB3B)),(LB1A,LB3A)))); | 4 | 0.06% |
| TREE 348 | 0.000569071 | (Lbru,((LB1A,LB2A),(LB3A,(LB2B,(LB1B,LB3B))))); | 4 | 0.06% |
| TREE 349 | 0.000569071 | (Lbru,(LB3B,(LB3A,(LB1B,(LB1A,(LB2A,LB2B)))))); | 4 | 0.06% |
| TREE 350 | 0.000569071 | (Lbru,((LB1B,LB2B),((LB1A,LB3B),(LB3A,LB2A)))); | 4 | 0.06% |
| TREE 351 | 0.000569071 | (Lbru,(LB2A,(LB1A,(LB3A,(LB2B,(LB1B,LB3B)))))); | 4 | 0.06% |
| TREE 352 | 0.000569071 | (Lbru,(LB2A,(LB1A,(LB3B,(LB2B,(LB1B,LB3A)))))); | 4 | 0.06% |
| TREE 353 | 0.000569071 | (Lbru,(LB3B,(LB1A,(LB2A,(LB3A,(LB1B,LB2B)))))); | 4 | 0.06% |
| TREE 354 | 0.000569071 | (Lbru,(LB3A,(LB2B,((LB1B,LB1A),(LB3B,LB2A))))); | 4 | 0.06% |
| TREE 355 | 0.000569071 | (Lbru,(LB1B,((LB2B,(LB3B,LB2A)),(LB1A,LB3A)))); | 4 | 0.06% |
| TREE 356 | 0.000569071 | (Lbru,(LB2A,(LB3A,(LB1B,(LB3B,(LB1A,LB2B)))))); | 4 | 0.06% |
| TREE 357 | 0.000569071 | (Lbru,((LB1B,(LB2A,(LB1A,LB3B))),(LB3A,LB2B))); | 4 | 0.06% |
| TREE 358 | 0.000569071 | (Lbru,(LB2B,((LB1A,LB2A),(LB3B,(LB1B,LB3A))))); | 4 | 0.06% |
| TREE 359 | 0.000569071 | (Lbru,((LB2B,(LB3B,(LB1B,LB1A))),(LB3A,LB2A))); | 4 | 0.06% |
| TREE 360 | 0.000569071 | (Lbru,(LB1A,(LB2A,((LB1B,LB3B),(LB3A,LB2B))))); | 4 | 0.06% |
| TREE 361 | 0.000569071 | (Lbru,(LB2B,(LB1A,(LB2A,(LB3B,(LB1B,LB3A)))))); | 4 | 0.06% |
| TREE 362 | 0.000569071 | (Lbru,(LB2A,((LB1B,LB2B),(LB1A,(LB3B,LB3A))))); | 4 | 0.06% |
| TREE 363 | 0.000569071 | (Lbru,(LB3B,(LB1A,((LB2A,LB2B),(LB1B,LB3A))))); | 4 | 0.06% |
| TREE 364 | 0.000569071 | (Lbru,(LB2A,(LB1A,(LB2B,(LB1B,(LB3B,LB3A)))))); | 4 | 0.06% |
| TREE 365 | 0.000569071 | (Lbru,(LB2B,((LB1B,LB2A),(LB3B,(LB1A,LB3A))))); | 4 | 0.06% |
| TREE 366 | 0.000569071 | (Lbru,((LB1B,LB2A),(LB3A,(LB3B,(LB1A,LB2B))))); | 4 | 0.06% |
| TREE 367 | 0.000569071 | (Lbru,(LB3A,(LB2B,(LB1B,(LB1A,(LB3B,LB2A)))))); | 4 | 0.06% |
| TREE 368 | 0.000569071 | (Lbru,(LB3B,(LB1A,(LB1B,(LB3A,(LB2A,LB2B)))))); | 4 | 0.06% |
| TREE 369 | 0.000569071 | (Lbru,(LB1B,(LB2B,(LB3A,(LB2A,(LB1A,LB3B)))))); | 4 | 0.06% |
| TREE 370 | 0.000569071 | (Lbru,((LB1B,LB2B),(LB2A,(LB3B,(LB1A,LB3A))))); | 4 | 0.06% |
| TREE 371 | 0.000569071 | (Lbru,(LB2A,((LB1A,LB2B),(LB3A,(LB1B,LB3B))))); | 4 | 0.06% |
| TREE 372 | 0.000569071 | (Lbru,(LB2B,(LB1A,((LB3B,LB2A),(LB1B,LB3A))))); | 4 | 0.06% |
| TREE 373 | 0.000569071 | (Lbru,((LB1B,LB2A),(LB2B,(LB1A,(LB3B,LB3A))))); | 4 | 0.06% |
| TREE 374 | 0.000569071 | (Lbru,((LB1B,LB2A),(LB2B,(LB3B,(LB1A,LB3A))))); | 4 | 0.06% |
| TREE 375 | 0.000569071 | (Lbru,((LB1A,LB2B),(LB1B,(LB3A,(LB3B,LB2A))))); | 4 | 0.06% |
| TREE 376 | 0.000569071 | (Lbru,(LB3B,(LB3A,(LB2A,(LB1B,(LB1A,LB2B)))))); | 4 | 0.06% |
| TREE 377 | 0.000569071 | (Lbru,(LB1B,(LB3A,(LB2A,(LB2B,(LB1A,LB3B)))))); | 4 | 0.06% |
| TREE 378 | 0.000569071 | (Lbru,((LB2A,(LB3A,(LB1B,LB3B))),(LB1A,LB2B))); | 4 | 0.06% |
| TREE 379 | 0.000569071 | (Lbru,(LB2A,(LB2B,(LB1A,(LB3B,(LB1B,LB3A)))))); | 4 | 0.06% |
| TREE 380 | 0.000569071 | (Lbru,(LB1B,(LB2B,(LB3A,(LB3B,(LB1A,LB2A)))))); | 4 | 0.06% |
| TREE 381 | 0.000569071 | (Lbru,(LB2B,(LB1B,(LB3B,(LB2A,(LB1A,LB3A)))))); | 4 | 0.06% |
| TREE 382 | 0.000569071 | (Lbru,(LB1A,(LB3A,(LB1B,(LB3B,(LB2A,LB2B)))))); | 4 | 0.06% |
| TREE 383 | 0.000569071 | (Lbru,(LB2A,(LB1B,(LB1A,(LB3B,(LB3A,LB2B)))))); | 4 | 0.06% |
| TREE 384 | 0.000569071 | (Lbru,((LB2A,(LB1B,(LB3B,LB3A))),(LB1A,LB2B))); | 4 | 0.06% |
| TREE 385 | 0.000569071 | (Lbru,(LB2B,(LB1B,(LB3A,(LB3B,(LB1A,LB2A)))))); | 4 | 0.06% |
| TREE 386 | 0.000569071 | (Lbru,((LB3B,LB2A),((LB1B,LB1A),(LB3A,LB2B)))); | 4 | 0.06% |
| TREE 387 | 0.000569071 | (Lbru,(LB3A,((LB2A,LB2B),(LB1A,(LB1B,LB3B))))); | 4 | 0.06% |
| TREE 388 | 0.000569071 | (Lbru,((LB3B,LB2A),(LB2B,(LB3A,(LB1B,LB1A))))); | 4 | 0.06% |
| TREE 389 | 0.000426803 | (Lbru,((LB2A,(LB1B,LB2B)),(LB3A,(LB1A,LB3B)))); | 3 | 0.04% |
| TREE 390 | 0.000426803 | (Lbru,((LB2B,(LB1B,LB2A)),(LB3A,(LB1A,LB3B)))); | 3 | 0.04% |
| TREE 391 | 0.000426803 | (Lbru,(LB2A,((LB1B,(LB3B,LB2B)),(LB1A,LB3A)))); | 3 | 0.04% |
| TREE 392 | 0.000426803 | (Lbru,(LB3B,((LB1A,(LB1B,LB2A)),(LB3A,LB2B)))); | 3 | 0.04% |
| TREE 393 | 0.000426803 | (Lbru,((LB1B,(LB2A,LB2B)),(LB3A,(LB1A,LB3B)))); | 3 | 0.04% |
| TREE 394 | 0.000426803 | (Lbru,(LB1A,(LB2A,(LB3B,(LB3A,(LB1B,LB2B)))))); | 3 | 0.04% |
| TREE 395 | 0.000426803 | (Lbru,((LB2A,(LB1A,LB2B)),(LB3A,(LB1B,LB3B)))); | 3 | 0.04% |
| TREE 396 | 0.000426803 | (Lbru,((LB2A,(LB3B,(LB1B,LB3A))),(LB1A,LB2B))); | 3 | 0.04% |
| TREE 397 | 0.000426803 | (Lbru,((LB1B,(LB1A,(LB3B,LB2B))),(LB3A,LB2A))); | 3 | 0.04% |
| TREE 398 | 0.000426803 | (Lbru,(LB1A,((LB1B,LB3B),(LB2B,(LB3A,LB2A))))); | 3 | 0.04% |
| TREE 399 | 0.000426803 | (Lbru,(LB1A,((LB1B,(LB3B,LB2B)),(LB3A,LB2A)))); | 3 | 0.04% |
| TREE 400 | 0.000426803 | (Lbru,(LB3A,(LB2A,(LB1A,(LB1B,(LB3B,LB2B)))))); | 3 | 0.04% |
| TREE 401 | 0.000426803 | (Lbru,(LB3B,((LB1A,(LB1B,LB2B)),(LB3A,LB2A)))); | 3 | 0.04% |
| TREE 402 | 0.000426803 | (Lbru,((LB1A,LB2B),((LB1B,LB2A),(LB3B,LB3A)))); | 3 | 0.04% |
| TREE 403 | 0.000426803 | (Lbru,((LB3B,(LB2B,(LB1B,LB2A))),(LB1A,LB3A))); | 3 | 0.04% |
| TREE 404 | 0.000426803 | (Lbru,((LB1A,LB2A),(LB2B,(LB1B,(LB3B,LB3A))))); | 3 | 0.04% |
| TREE 405 | 0.000426803 | (Lbru,(LB1B,((LB2A,(LB3B,LB2B)),(LB1A,LB3A)))); | 3 | 0.04% |
| TREE 406 | 0.000426803 | (Lbru,(LB2A,(LB3B,(LB1B,(LB3A,(LB1A,LB2B)))))); | 3 | 0.04% |
| TREE 407 | 0.000426803 | (Lbru,(LB2A,(LB3A,(LB1B,(LB2B,(LB1A,LB3B)))))); | 3 | 0.04% |
| TREE 408 | 0.000426803 | (Lbru,(LB3A,(LB3B,(LB2B,(LB1A,(LB1B,LB2A)))))); | 3 | 0.04% |
| TREE 409 | 0.000426803 | (Lbru,(LB2B,(LB3A,(LB3B,(LB1B,(LB1A,LB2A)))))); | 3 | 0.04% |
| TREE 410 | 0.000426803 | (Lbru,(LB3A,(LB1B,(LB2B,(LB3B,(LB1A,LB2A)))))); | 3 | 0.04% |
| TREE 411 | 0.000426803 | (Lbru,((LB2A,(LB3B,(LB1B,LB1A))),(LB3A,LB2B))); | 3 | 0.04% |
| TREE 412 | 0.000426803 | (Lbru,(LB1B,(LB2A,(LB1A,(LB3A,(LB3B,LB2B)))))); | 3 | 0.04% |
| TREE 413 | 0.000426803 | (Lbru,(LB3A,(LB3B,(LB1A,(LB2A,(LB1B,LB2B)))))); | 3 | 0.04% |
| TREE 414 | 0.000426803 | (Lbru,(LB2A,(LB1A,(LB3B,(LB3A,(LB1B,LB2B)))))); | 3 | 0.04% |
| TREE 415 | 0.000426803 | (Lbru,((LB2A,LB2B),(LB3B,(LB1B,(LB1A,LB3A))))); | 3 | 0.04% |
| TREE 416 | 0.000426803 | (Lbru,(LB3B,((LB1A,LB2A),(LB2B,(LB1B,LB3A))))); | 3 | 0.04% |
| TREE 417 | 0.000426803 | (Lbru,(LB1B,(LB3B,(LB1A,(LB3A,(LB2A,LB2B)))))); | 3 | 0.04% |
| TREE 418 | 0.000426803 | (Lbru,((LB1A,(LB1B,(LB3B,LB2A))),(LB3A,LB2B))); | 3 | 0.04% |
| TREE 419 | 0.000426803 | (Lbru,(LB3A,(LB2B,((LB1A,LB2A),(LB1B,LB3B))))); | 3 | 0.04% |
| TREE 420 | 0.000426803 | (Lbru,(LB1A,(LB2A,(LB1B,(LB3B,(LB3A,LB2B)))))); | 3 | 0.04% |
| TREE 421 | 0.000426803 | (Lbru,((LB3B,(LB2A,LB2B)),(LB1B,(LB1A,LB3A)))); | 3 | 0.04% |
| TREE 422 | 0.000426803 | (Lbru,(LB3B,((LB1B,(LB1A,LB2B)),(LB3A,LB2A)))); | 3 | 0.04% |
| TREE 423 | 0.000426803 | (Lbru,(LB2A,(LB1A,((LB1B,LB3B),(LB3A,LB2B))))); | 3 | 0.04% |
| TREE 424 | 0.000426803 | (Lbru,(LB1B,((LB1A,LB3B),(LB2B,(LB3A,LB2A))))); | 3 | 0.04% |
| TREE 425 | 0.000426803 | (Lbru,((LB1A,(LB1B,LB2A)),(LB3B,(LB3A,LB2B)))); | 3 | 0.04% |
| TREE 426 | 0.000426803 | (Lbru,((LB1B,(LB1A,LB3B)),(LB2A,(LB3A,LB2B)))); | 3 | 0.04% |
| TREE 427 | 0.000426803 | (Lbru,((LB3B,(LB1A,LB2B)),(LB2A,(LB1B,LB3A)))); | 3 | 0.04% |
| TREE 428 | 0.000426803 | (Lbru,(LB2B,(LB1B,(LB2A,(LB1A,(LB3B,LB3A)))))); | 3 | 0.04% |
| TREE 429 | 0.000426803 | (Lbru,(LB1B,((LB1A,LB2B),(LB3B,(LB3A,LB2A))))); | 3 | 0.04% |
| TREE 430 | 0.000426803 | (Lbru,((LB2B,(LB3B,(LB1A,LB2A))),(LB1B,LB3A))); | 3 | 0.04% |
| TREE 431 | 0.000426803 | (Lbru,(LB2B,(LB2A,((LB1A,LB3B),(LB1B,LB3A))))); | 3 | 0.04% |
| TREE 432 | 0.000426803 | (Lbru,((LB2A,LB2B),((LB1B,LB3B),(LB1A,LB3A)))); | 3 | 0.04% |
| TREE 433 | 0.000426803 | (Lbru,((LB2A,LB2B),(LB1A,(LB3B,(LB1B,LB3A))))); | 3 | 0.04% |
| TREE 434 | 0.000426803 | (Lbru,(LB2A,(LB3B,(LB2B,(LB1B,(LB1A,LB3A)))))); | 3 | 0.04% |
| TREE 435 | 0.000426803 | (Lbru,((LB3B,LB2B),(LB3A,(LB1B,(LB1A,LB2A))))); | 3 | 0.04% |
| TREE 436 | 0.000426803 | (Lbru,(LB1B,((LB3B,(LB1A,LB2A)),(LB3A,LB2B)))); | 3 | 0.04% |
| TREE 437 | 0.000426803 | (Lbru,(LB3B,(LB3A,(LB2A,(LB1A,(LB1B,LB2B)))))); | 3 | 0.04% |
| TREE 438 | 0.000426803 | (Lbru,(LB3A,(LB2A,(LB2B,(LB1B,(LB1A,LB3B)))))); | 3 | 0.04% |
| TREE 439 | 0.000426803 | (Lbru,((LB3B,(LB1B,(LB2A,LB2B))),(LB1A,LB3A))); | 3 | 0.04% |
| TREE 440 | 0.000426803 | (Lbru,(LB3A,(LB2A,(LB1B,(LB1A,(LB3B,LB2B)))))); | 3 | 0.04% |
| TREE 441 | 0.000426803 | (Lbru,(LB3B,(LB1B,(LB2A,(LB3A,(LB1A,LB2B)))))); | 3 | 0.04% |
| TREE 442 | 0.000426803 | (Lbru,(LB1B,((LB1A,LB2A),(LB3B,(LB3A,LB2B))))); | 3 | 0.04% |
| TREE 443 | 0.000426803 | (Lbru,(LB2B,(LB2A,(LB3A,(LB1B,(LB1A,LB3B)))))); | 3 | 0.04% |
| TREE 444 | 0.000426803 | (Lbru,((LB2B,(LB3B,LB2A)),(LB1A,(LB1B,LB3A)))); | 3 | 0.04% |
| TREE 445 | 0.000426803 | (Lbru,(LB3B,(LB1B,(LB3A,(LB2A,(LB1A,LB2B)))))); | 3 | 0.04% |
| TREE 446 | 0.000426803 | (Lbru,((LB1B,LB3B),((LB1A,LB2B),(LB3A,LB2A)))); | 3 | 0.04% |
| TREE 447 | 0.000426803 | (Lbru,((LB3B,LB2A),(LB3A,(LB1A,(LB1B,LB2B))))); | 3 | 0.04% |
| TREE 448 | 0.000426803 | (Lbru,(LB1A,(LB2B,(LB3B,(LB3A,(LB1B,LB2A)))))); | 3 | 0.04% |
| TREE 449 | 0.000426803 | (Lbru,(LB3B,(LB1B,((LB1A,LB2B),(LB3A,LB2A))))); | 3 | 0.04% |
| TREE 450 | 0.000426803 | (Lbru,(LB3B,(LB2B,(LB1A,(LB3A,(LB1B,LB2A)))))); | 3 | 0.04% |
| TREE 451 | 0.000426803 | (Lbru,(LB3A,((LB1B,LB2A),(LB1A,(LB3B,LB2B))))); | 3 | 0.04% |
| TREE 452 | 0.000426803 | (Lbru,(LB3B,((LB2A,(LB1B,LB2B)),(LB1A,LB3A)))); | 3 | 0.04% |
| TREE 453 | 0.000426803 | (Lbru,(LB2A,(LB3B,(LB3A,(LB1B,(LB1A,LB2B)))))); | 3 | 0.04% |
| TREE 454 | 0.000426803 | (Lbru,(LB1A,((LB3B,LB2A),(LB1B,(LB3A,LB2B))))); | 3 | 0.04% |
| TREE 455 | 0.000426803 | (Lbru,(LB3A,(LB1A,(LB1B,(LB2B,(LB3B,LB2A)))))); | 3 | 0.04% |
| TREE 456 | 0.000426803 | (Lbru,((LB1B,LB2A),(LB1A,(LB3A,(LB3B,LB2B))))); | 3 | 0.04% |
| TREE 457 | 0.000426803 | (Lbru,((LB1B,LB2B),(LB1A,(LB3A,(LB3B,LB2A))))); | 3 | 0.04% |
| TREE 458 | 0.000426803 | (Lbru,((LB1B,(LB2B,(LB1A,LB3B))),(LB3A,LB2A))); | 3 | 0.04% |
| TREE 459 | 0.000426803 | (Lbru,((LB1A,LB2A),(LB1B,(LB3B,(LB3A,LB2B))))); | 3 | 0.04% |
| TREE 460 | 0.000426803 | (Lbru,(LB1B,((LB3B,(LB1A,LB2B)),(LB3A,LB2A)))); | 3 | 0.04% |
| TREE 461 | 0.000426803 | (Lbru,((LB1B,LB3B),(LB2A,(LB3A,(LB1A,LB2B))))); | 3 | 0.04% |
| TREE 462 | 0.000426803 | (Lbru,(LB1B,(LB3B,(LB1A,(LB2B,(LB3A,LB2A)))))); | 3 | 0.04% |
| TREE 463 | 0.000426803 | (Lbru,((LB2B,(LB3B,(LB1B,LB3A))),(LB1A,LB2A))); | 3 | 0.04% |
| TREE 464 | 0.000426803 | (Lbru,((LB2B,(LB2A,(LB1A,LB3B))),(LB1B,LB3A))); | 3 | 0.04% |
| TREE 465 | 0.000426803 | (Lbru,(LB1A,(LB3B,(LB3A,(LB2B,(LB1B,LB2A)))))); | 3 | 0.04% |
| TREE 466 | 0.000426803 | (Lbru,(LB2B,(LB1A,(LB3B,(LB3A,(LB1B,LB2A)))))); | 3 | 0.04% |
| TREE 467 | 0.000426803 | (Lbru,(LB1B,(LB2B,(LB1A,(LB3B,(LB3A,LB2A)))))); | 3 | 0.04% |
| TREE 468 | 0.000426803 | (Lbru,((LB2B,(LB1A,LB2A)),(LB3B,(LB1B,LB3A)))); | 3 | 0.04% |
| TREE 469 | 0.000426803 | (Lbru,(LB2A,((LB1A,LB2B),(LB3B,(LB1B,LB3A))))); | 3 | 0.04% |
| TREE 470 | 0.000426803 | (Lbru,(LB1A,(LB3B,(LB2A,(LB3A,(LB1B,LB2B)))))); | 3 | 0.04% |
| TREE 471 | 0.000426803 | (Lbru,(LB2B,(LB2A,(LB1A,(LB3B,(LB1B,LB3A)))))); | 3 | 0.04% |
| TREE 472 | 0.000426803 | (Lbru,(LB3A,(LB2A,(LB1A,(LB2B,(LB1B,LB3B)))))); | 3 | 0.04% |
| TREE 473 | 0.000426803 | (Lbru,(LB1B,(LB2B,((LB3B,LB2A),(LB1A,LB3A))))); | 3 | 0.04% |
| TREE 474 | 0.000426803 | (Lbru,(LB3B,(LB1B,(LB1A,(LB3A,(LB2A,LB2B)))))); | 3 | 0.04% |
| TREE 475 | 0.000426803 | (Lbru,(LB1A,(LB3B,(LB3A,(LB2A,(LB1B,LB2B)))))); | 3 | 0.04% |
| TREE 476 | 0.000426803 | (Lbru,((LB3B,LB2B),(LB1A,(LB1B,(LB3A,LB2A))))); | 3 | 0.04% |
| TREE 477 | 0.000426803 | (Lbru,(LB1B,(LB3B,(LB1A,(LB2A,(LB3A,LB2B)))))); | 3 | 0.04% |
| TREE 478 | 0.000426803 | (Lbru,(LB2B,((LB3B,LB2A),(LB1B,(LB1A,LB3A))))); | 3 | 0.04% |
| TREE 479 | 0.000426803 | (Lbru,(LB3A,((LB1A,(LB2A,LB2B)),(LB1B,LB3B)))); | 3 | 0.04% |
| TREE 480 | 0.000426803 | (Lbru,((LB3B,LB2A),(LB3A,(LB2B,(LB1B,LB1A))))); | 3 | 0.04% |
| TREE 481 | 0.000426803 | (Lbru,(LB1A,(LB3A,(LB2A,(LB2B,(LB1B,LB3B)))))); | 3 | 0.04% |
| TREE 482 | 0.000426803 | (Lbru,(LB2B,(LB1A,(LB1B,(LB3B,(LB3A,LB2A)))))); | 3 | 0.04% |
| TREE 483 | 0.000426803 | (Lbru,(LB2A,(LB3A,(LB3B,(LB1B,(LB1A,LB2B)))))); | 3 | 0.04% |
| TREE 484 | 0.000426803 | (Lbru,(LB1A,(LB3A,(LB1B,(LB2B,(LB3B,LB2A)))))); | 3 | 0.04% |
| TREE 485 | 0.000426803 | (Lbru,((LB1A,(LB1B,LB3B)),(LB3A,(LB2A,LB2B)))); | 3 | 0.04% |
| TREE 486 | 0.000284535 | (Lbru,(LB2A,(LB1B,((LB1A,LB3B),(LB3A,LB2B))))); | 2 | 0.03% |
| TREE 487 | 0.000284535 | (Lbru,((LB2A,(LB1A,LB3B)),(LB1B,(LB3A,LB2B)))); | 2 | 0.03% |
| TREE 488 | 0.000284535 | (Lbru,(LB2A,(LB1A,(LB1B,(LB3B,(LB3A,LB2B)))))); | 2 | 0.03% |
| TREE 489 | 0.000284535 | (Lbru,((LB3B,LB2A),(LB1B,(LB1A,(LB3A,LB2B))))); | 2 | 0.03% |
| TREE 490 | 0.000284535 | (Lbru,(LB3B,((LB2B,(LB1A,LB2A)),(LB1B,LB3A)))); | 2 | 0.03% |
| TREE 491 | 0.000284535 | (Lbru,(LB1B,((LB3B,LB2A),(LB1A,(LB3A,LB2B))))); | 2 | 0.03% |
| TREE 492 | 0.000284535 | (Lbru,((LB1B,(LB2A,(LB3B,LB2B))),(LB1A,LB3A))); | 2 | 0.03% |
| TREE 493 | 0.000284535 | (Lbru,(LB3B,(LB3A,(LB1A,(LB2B,(LB1B,LB2A)))))); | 2 | 0.03% |
| TREE 494 | 0.000284535 | (Lbru,((LB1B,LB2B),(LB3A,(LB2A,(LB1A,LB3B))))); | 2 | 0.03% |
| TREE 495 | 0.000284535 | (Lbru,(LB1B,(LB3A,(LB2B,(LB3B,(LB1A,LB2A)))))); | 2 | 0.03% |
| TREE 496 | 0.000284535 | (Lbru,((LB3B,(LB1A,LB2A)),(LB1B,(LB3A,LB2B)))); | 2 | 0.03% |
| TREE 497 | 0.000284535 | (Lbru,(LB3A,(LB3B,(LB1B,(LB2B,(LB1A,LB2A)))))); | 2 | 0.03% |
| TREE 498 | 0.000284535 | (Lbru,(LB3B,((LB1A,LB2B),(LB3A,(LB1B,LB2A))))); | 2 | 0.03% |
| TREE 499 | 0.000284535 | (Lbru,((LB3B,LB2A),(LB2B,(LB1B,(LB1A,LB3A))))); | 2 | 0.03% |
| TREE 500 | 0.000284535 | (Lbru,(LB2A,((LB3B,(LB1A,LB2B)),(LB1B,LB3A)))); | 2 | 0.03% |
| TREE 501 | 0.000284535 | (Lbru,(LB3B,(LB2B,(LB3A,(LB1A,(LB1B,LB2A)))))); | 2 | 0.03% |
| TREE 502 | 0.000284535 | (Lbru,((LB1A,(LB3B,LB2A)),(LB2B,(LB1B,LB3A)))); | 2 | 0.03% |
| TREE 503 | 0.000284535 | (Lbru,(LB1B,(LB3B,(LB2A,(LB1A,(LB3A,LB2B)))))); | 2 | 0.03% |
| TREE 504 | 0.000284535 | (Lbru,(LB3B,(LB1B,(LB2A,(LB1A,(LB3A,LB2B)))))); | 2 | 0.03% |
| TREE 505 | 0.000284535 | (Lbru,(LB2B,((LB1B,(LB1A,LB3B)),(LB3A,LB2A)))); | 2 | 0.03% |
| TREE 506 | 0.000284535 | (Lbru,((LB2A,(LB1A,LB3B)),(LB3A,(LB1B,LB2B)))); | 2 | 0.03% |
| TREE 507 | 0.000284535 | (Lbru,(LB3A,(LB2B,((LB1B,LB2A),(LB1A,LB3B))))); | 2 | 0.03% |
| TREE 508 | 0.000284535 | (Lbru,(LB1A,(LB3B,(LB1B,(LB2A,(LB3A,LB2B)))))); | 2 | 0.03% |
| TREE 509 | 0.000284535 | (Lbru,(LB1A,(LB3A,((LB1B,LB2A),(LB3B,LB2B))))); | 2 | 0.03% |
| TREE 510 | 0.000284535 | (Lbru,((LB1A,LB2B),(LB3B,(LB3A,(LB1B,LB2A))))); | 2 | 0.03% |
| TREE 511 | 0.000284535 | (Lbru,(LB1A,(LB3A,(LB3B,(LB2A,(LB1B,LB2B)))))); | 2 | 0.03% |
| TREE 512 | 0.000284535 | (Lbru,(((LB2A,LB2B),(LB1B,LB3B)),(LB1A,LB3A))); | 2 | 0.03% |
| TREE 513 | 0.000284535 | (Lbru,(LB3A,(LB3B,(LB2A,(LB1A,(LB1B,LB2B)))))); | 2 | 0.03% |
| TREE 514 | 0.000284535 | (Lbru,(LB2A,((LB1B,LB3B),(LB2B,(LB1A,LB3A))))); | 2 | 0.03% |
| TREE 515 | 0.000284535 | (Lbru,(LB1B,((LB1A,(LB3B,LB2A)),(LB3A,LB2B)))); | 2 | 0.03% |
| TREE 516 | 0.000284535 | (Lbru,(LB2B,(LB3A,(LB2A,(LB1A,(LB1B,LB3B)))))); | 2 | 0.03% |
| TREE 517 | 0.000284535 | (Lbru,(((LB2A,LB2B),(LB1A,LB3B)),(LB1B,LB3A))); | 2 | 0.03% |
| TREE 518 | 0.000284535 | (Lbru,(LB1B,(LB3A,(LB2A,(LB3B,(LB1A,LB2B)))))); | 2 | 0.03% |
| TREE 519 | 0.000284535 | (Lbru,(LB2B,((LB1A,LB3B),(LB1B,(LB3A,LB2A))))); | 2 | 0.03% |
| TREE 520 | 0.000284535 | (Lbru,(LB3B,(LB1A,(LB2B,(LB3A,(LB1B,LB2A)))))); | 2 | 0.03% |
| TREE 521 | 0.000284535 | (Lbru,(LB3A,(LB2A,(LB3B,(LB1B,(LB1A,LB2B)))))); | 2 | 0.03% |
| TREE 522 | 0.000284535 | (Lbru,(LB3A,(LB1B,(LB1A,(LB3B,(LB2A,LB2B)))))); | 2 | 0.03% |
| TREE 523 | 0.000284535 | (Lbru,(LB2B,((LB3B,LB2A),(LB1A,(LB1B,LB3A))))); | 2 | 0.03% |
| TREE 524 | 0.000284535 | (Lbru,(LB1B,((LB1A,LB3B),(LB2A,(LB3A,LB2B))))); | 2 | 0.03% |
| TREE 525 | 0.000284535 | (Lbru,(LB1B,(LB2B,(LB3A,(LB1A,(LB3B,LB2A)))))); | 2 | 0.03% |
| TREE 526 | 0.000284535 | (Lbru,(LB2A,(LB3B,(LB2B,(LB3A,(LB1B,LB1A)))))); | 2 | 0.03% |
| TREE 527 | 0.000284535 | (Lbru,(LB1B,(LB3A,((LB1A,LB2A),(LB3B,LB2B))))); | 2 | 0.03% |
| TREE 528 | 0.000284535 | (Lbru,((LB3B,(LB1A,(LB2A,LB2B))),(LB1B,LB3A))); | 2 | 0.03% |
| TREE 529 | 0.000284535 | (Lbru,(LB2A,((LB1A,LB3B),(LB2B,(LB1B,LB3A))))); | 2 | 0.03% |
| TREE 530 | 0.000284535 | (Lbru,(LB3A,(LB1B,(LB1A,(LB2A,(LB3B,LB2B)))))); | 2 | 0.03% |
| TREE 531 | 0.000284535 | (Lbru,(LB3A,((LB1B,(LB2A,LB2B)),(LB1A,LB3B)))); | 2 | 0.03% |
| TREE 532 | 0.000284535 | (Lbru,(LB3B,(LB3A,(LB1A,(LB2A,(LB1B,LB2B)))))); | 2 | 0.03% |
| TREE 533 | 0.000284535 | (Lbru,(LB2A,((LB1A,LB3B),(LB1B,(LB3A,LB2B))))); | 2 | 0.03% |
| TREE 534 | 0.000284535 | (Lbru,(LB3B,(LB2A,(LB1B,(LB1A,(LB3A,LB2B)))))); | 2 | 0.03% |
| TREE 535 | 0.000284535 | (Lbru,(LB2B,(LB2A,(LB3B,(LB1A,(LB1B,LB3A)))))); | 2 | 0.03% |
| TREE 536 | 0.000284535 | (Lbru,(LB2A,(LB1A,((LB1B,LB2B),(LB3B,LB3A))))); | 2 | 0.03% |
| TREE 537 | 0.000284535 | (Lbru,(LB3B,((LB1B,LB2B),(LB2A,(LB1A,LB3A))))); | 2 | 0.03% |
| TREE 538 | 0.000284535 | (Lbru,(LB2B,(LB3A,(LB1B,(LB3B,(LB1A,LB2A)))))); | 2 | 0.03% |
| TREE 539 | 0.000284535 | (Lbru,(LB2A,(LB3B,(LB3A,(LB1A,(LB1B,LB2B)))))); | 2 | 0.03% |
| TREE 540 | 0.000284535 | (Lbru,((LB2B,(LB3B,LB2A)),(LB1B,(LB1A,LB3A)))); | 2 | 0.03% |
| TREE 541 | 0.000284535 | (Lbru,((LB1B,(LB3B,LB2B)),(LB1A,(LB3A,LB2A)))); | 2 | 0.03% |
| TREE 542 | 0.000284535 | (Lbru,((LB2A,(LB3B,LB2B)),(LB1B,(LB1A,LB3A)))); | 2 | 0.03% |
| TREE 543 | 0.000284535 | (Lbru,(LB1A,(LB2A,(LB1B,(LB3A,(LB3B,LB2B)))))); | 2 | 0.03% |
| TREE 544 | 0.000284535 | (Lbru,(LB3B,(LB1A,((LB1B,LB2A),(LB3A,LB2B))))); | 2 | 0.03% |
| TREE 545 | 0.000284535 | (Lbru,((LB1A,LB2A),((LB1B,LB3B),(LB3A,LB2B)))); | 2 | 0.03% |
| TREE 546 | 0.000284535 | (Lbru,(LB2B,(LB3A,(LB1A,(LB1B,(LB3B,LB2A)))))); | 2 | 0.03% |
| TREE 547 | 0.000284535 | (Lbru,(LB1B,((LB3B,LB2B),(LB2A,(LB1A,LB3A))))); | 2 | 0.03% |
| TREE 548 | 0.000284535 | (Lbru,((LB3B,LB2B),(LB2A,(LB1B,(LB1A,LB3A))))); | 2 | 0.03% |
| TREE 549 | 0.000284535 | (Lbru,(LB3B,(LB2A,(LB3A,(LB1B,(LB1A,LB2B)))))); | 2 | 0.03% |
| TREE 550 | 0.000284535 | (Lbru,(LB2B,(LB1B,(LB3B,(LB3A,(LB1A,LB2A)))))); | 2 | 0.03% |
| TREE 551 | 0.000284535 | (Lbru,(LB3B,(LB2B,((LB1A,LB2A),(LB1B,LB3A))))); | 2 | 0.03% |
| TREE 552 | 0.000284535 | (Lbru,((LB1B,LB2B),(LB1A,(LB3B,(LB3A,LB2A))))); | 2 | 0.03% |
| TREE 553 | 0.000284535 | (Lbru,(LB2B,((LB1B,(LB3B,LB2A)),(LB1A,LB3A)))); | 2 | 0.03% |
| TREE 554 | 0.000284535 | (Lbru,(LB2B,((LB1B,LB3B),(LB1A,(LB3A,LB2A))))); | 2 | 0.03% |
| TREE 555 | 0.000284535 | (Lbru,((LB1B,LB2A),(LB3B,(LB1A,(LB3A,LB2B))))); | 2 | 0.03% |
| TREE 556 | 0.000284535 | (Lbru,(LB3A,(LB3B,((LB1B,LB2B),(LB1A,LB2A))))); | 2 | 0.03% |
| TREE 557 | 0.000284535 | (Lbru,(LB3B,(LB2B,(LB2A,(LB1B,(LB1A,LB3A)))))); | 2 | 0.03% |
| TREE 558 | 0.000284535 | (Lbru,(LB1A,(LB2B,(LB3B,(LB2A,(LB1B,LB3A)))))); | 2 | 0.03% |
| TREE 559 | 0.000284535 | (Lbru,(LB2B,(LB3A,(LB1B,(LB2A,(LB1A,LB3B)))))); | 2 | 0.03% |
| TREE 560 | 0.000284535 | (Lbru,(LB3A,(LB2B,(LB1B,(LB2A,(LB1A,LB3B)))))); | 2 | 0.03% |
| TREE 561 | 0.000284535 | (Lbru,((LB1A,LB2A),(LB3B,(LB2B,(LB1B,LB3A))))); | 2 | 0.03% |
| TREE 562 | 0.000284535 | (Lbru,(LB2B,(LB1B,(LB1A,(LB3B,(LB3A,LB2A)))))); | 2 | 0.03% |
| TREE 563 | 0.000284535 | (Lbru,(LB2B,(LB3A,(LB1A,(LB3B,(LB1B,LB2A)))))); | 2 | 0.03% |
| TREE 564 | 0.000284535 | (Lbru,(LB2B,(LB3A,(LB2A,(LB1B,(LB1A,LB3B)))))); | 2 | 0.03% |
| TREE 565 | 0.000284535 | (Lbru,(LB3A,(LB2B,(LB1B,(LB3B,(LB1A,LB2A)))))); | 2 | 0.03% |
| TREE 566 | 0.000284535 | (Lbru,((LB3B,(LB1A,(LB1B,LB2B))),(LB3A,LB2A))); | 2 | 0.03% |
| TREE 567 | 0.000284535 | (Lbru,((LB3B,LB2A),(LB3A,(LB1B,(LB1A,LB2B))))); | 2 | 0.03% |
| TREE 568 | 0.000284535 | (Lbru,((LB3B,(LB2A,(LB1B,LB2B))),(LB1A,LB3A))); | 2 | 0.03% |
| TREE 569 | 0.000284535 | (Lbru,(LB2A,((LB3B,(LB1B,LB2B)),(LB1A,LB3A)))); | 2 | 0.03% |
| TREE 570 | 0.000284535 | (Lbru,(LB3A,(LB2A,((LB1A,LB2B),(LB1B,LB3B))))); | 2 | 0.03% |
| TREE 571 | 0.000284535 | (Lbru,(LB3B,(LB1B,(LB3A,(LB2B,(LB1A,LB2A)))))); | 2 | 0.03% |
| TREE 572 | 0.000284535 | (Lbru,(LB3A,(LB1B,(LB2A,(LB2B,(LB1A,LB3B)))))); | 2 | 0.03% |
| TREE 573 | 0.000284535 | (Lbru,(LB1B,(LB2B,(LB3B,(LB1A,(LB3A,LB2A)))))); | 2 | 0.03% |
| TREE 574 | 0.000284535 | (Lbru,(LB3B,(LB1A,((LB1B,LB2B),(LB3A,LB2A))))); | 2 | 0.03% |
| TREE 575 | 0.000284535 | (Lbru,((LB3B,(LB1B,(LB1A,LB2B))),(LB3A,LB2A))); | 2 | 0.03% |
| TREE 576 | 0.000284535 | (Lbru,(LB2A,(LB3A,((LB1B,LB2B),(LB1A,LB3B))))); | 2 | 0.03% |
| TREE 577 | 0.000284535 | (Lbru,(LB3A,(LB1A,(LB2A,(LB2B,(LB1B,LB3B)))))); | 2 | 0.03% |
| TREE 578 | 0.000284535 | (Lbru,((LB2B,(LB1A,LB3B)),(LB2A,(LB1B,LB3A)))); | 2 | 0.03% |
| TREE 579 | 0.000284535 | (Lbru,((LB1B,LB2A),(LB3B,(LB3A,(LB1A,LB2B))))); | 2 | 0.03% |
| TREE 580 | 0.000284535 | (Lbru,(LB2B,((LB1A,(LB3B,LB2A)),(LB1B,LB3A)))); | 2 | 0.03% |
| TREE 581 | 0.000284535 | (Lbru,(LB1B,(LB3B,((LB1A,LB2A),(LB3A,LB2B))))); | 2 | 0.03% |
| TREE 582 | 0.000284535 | (Lbru,(LB2B,(LB1A,(LB3B,(LB1B,(LB3A,LB2A)))))); | 2 | 0.03% |
| TREE 583 | 0.000284535 | (Lbru,(LB1B,((LB2B,(LB1A,LB3B)),(LB3A,LB2A)))); | 2 | 0.03% |
| TREE 584 | 0.000284535 | (Lbru,(LB3B,(LB2A,((LB1A,LB2B),(LB1B,LB3A))))); | 2 | 0.03% |
| TREE 585 | 0.000284535 | (Lbru,(LB3B,((LB1B,LB2B),(LB1A,(LB3A,LB2A))))); | 2 | 0.03% |
| TREE 586 | 0.000284535 | (Lbru,((LB2B,(LB3B,(LB1B,LB2A))),(LB1A,LB3A))); | 2 | 0.03% |
| TREE 587 | 0.000284535 | (Lbru,(LB1B,((LB3B,LB2B),(LB1A,(LB3A,LB2A))))); | 2 | 0.03% |
| TREE 588 | 0.000284535 | (Lbru,((LB2A,LB2B),(LB3A,(LB1A,(LB1B,LB3B))))); | 2 | 0.03% |
| TREE 589 | 0.000284535 | (Lbru,(LB1A,(LB2B,(LB3A,(LB3B,(LB1B,LB2A)))))); | 2 | 0.03% |
| TREE 590 | 0.000284535 | (Lbru,((LB2B,(LB1B,LB3B)),(LB2A,(LB1A,LB3A)))); | 2 | 0.03% |
| TREE 591 | 0.000284535 | (Lbru,((LB1A,LB2B),(LB3B,(LB1B,(LB3A,LB2A))))); | 2 | 0.03% |
| TREE 592 | 0.000284535 | (Lbru,((LB3B,LB2A),(LB1A,(LB3A,(LB1B,LB2B))))); | 2 | 0.03% |
| TREE 593 | 0.000284535 | (Lbru,((LB3B,LB2A),(LB1A,(LB1B,(LB3A,LB2B))))); | 2 | 0.03% |
| TREE 594 | 0.000284535 | (Lbru,(LB3A,((LB1A,LB2A),(LB1B,(LB3B,LB2B))))); | 2 | 0.03% |
| TREE 595 | 0.000284535 | (Lbru,((LB1A,(LB2B,(LB3B,LB2A))),(LB1B,LB3A))); | 2 | 0.03% |
| TREE 596 | 0.000284535 | (Lbru,(LB3A,((LB1A,(LB1B,LB2A)),(LB3B,LB2B)))); | 2 | 0.03% |
| TREE 597 | 0.000284535 | (Lbru,(LB3A,(LB2A,((LB1B,LB2B),(LB1A,LB3B))))); | 2 | 0.03% |
| TREE 598 | 0.000284535 | (Lbru,(LB3A,(LB1B,((LB2A,LB2B),(LB1A,LB3B))))); | 2 | 0.03% |
| TREE 599 | 0.000284535 | (Lbru,(LB3B,(LB1A,(LB3A,(LB1B,(LB2A,LB2B)))))); | 2 | 0.03% |
| TREE 600 | 0.000284535 | (Lbru,(LB3A,((LB2A,(LB1B,LB2B)),(LB1A,LB3B)))); | 2 | 0.03% |
| TREE 601 | 0.000284535 | (Lbru,((LB1B,(LB3B,(LB2A,LB2B))),(LB1A,LB3A))); | 2 | 0.03% |
| TREE 602 | 0.000284535 | (Lbru,(LB3A,((LB1B,LB2A),(LB3B,(LB1A,LB2B))))); | 2 | 0.03% |
| TREE 603 | 0.000284535 | (Lbru,((LB1B,(LB3B,LB2B)),(LB2A,(LB1A,LB3A)))); | 2 | 0.03% |
| TREE 604 | 0.000284535 | (Lbru,(LB1B,(LB3A,((LB1A,LB2B),(LB3B,LB2A))))); | 2 | 0.03% |
| TREE 605 | 0.000284535 | (Lbru,(LB3A,(LB1A,(LB3B,(LB1B,(LB2A,LB2B)))))); | 2 | 0.03% |
| TREE 606 | 0.000284535 | (Lbru,(LB3B,(LB1B,(LB3A,(LB1A,(LB2A,LB2B)))))); | 2 | 0.03% |
| TREE 607 | 0.000284535 | (Lbru,(LB3B,(LB2A,(LB2B,(LB1B,(LB1A,LB3A)))))); | 2 | 0.03% |
| TREE 608 | 0.000284535 | (Lbru,((LB1B,LB3B),(LB1A,(LB2A,(LB3A,LB2B))))); | 2 | 0.03% |
| TREE 609 | 0.000284535 | (Lbru,(LB1A,(LB3A,((LB1B,LB2B),(LB3B,LB2A))))); | 2 | 0.03% |
| TREE 610 | 0.000284535 | (Lbru,(LB1A,(LB2A,((LB3B,LB2B),(LB1B,LB3A))))); | 2 | 0.03% |
| TREE 611 | 0.000284535 | (Lbru,(LB3B,(LB2A,((LB1B,LB2B),(LB1A,LB3A))))); | 2 | 0.03% |
| TREE 612 | 0.000284535 | (Lbru,(LB3A,(LB1A,((LB2A,LB2B),(LB1B,LB3B))))); | 2 | 0.03% |
| TREE 613 | 0.000284535 | (Lbru,((LB1A,LB2A),(LB2B,(LB3A,(LB1B,LB3B))))); | 2 | 0.03% |
| TREE 614 | 0.000284535 | (Lbru,(LB3B,(LB1B,(LB1A,(LB2A,(LB3A,LB2B)))))); | 2 | 0.03% |
| TREE 615 | 0.000284535 | (Lbru,(LB1A,((LB3B,LB2A),(LB3A,(LB1B,LB2B))))); | 2 | 0.03% |
| TREE 616 | 0.000284535 | (Lbru,(LB2B,(LB1A,(LB3A,(LB1B,(LB3B,LB2A)))))); | 2 | 0.03% |
| TREE 617 | 0.000284535 | (Lbru,((LB1A,LB3B),(LB1B,(LB2A,(LB3A,LB2B))))); | 2 | 0.03% |
| TREE 618 | 0.000284535 | (Lbru,(LB1B,(LB3A,((LB2A,LB2B),(LB1A,LB3B))))); | 2 | 0.03% |
| TREE 619 | 0.000284535 | (Lbru,(LB2B,((LB3B,(LB1A,LB2A)),(LB1B,LB3A)))); | 2 | 0.03% |
| TREE 620 | 0.000284535 | (Lbru,((LB2A,(LB3B,LB2B)),(LB1A,(LB1B,LB3A)))); | 2 | 0.03% |
| TREE 621 | 0.000284535 | (Lbru,(LB2A,((LB1B,(LB1A,LB3B)),(LB3A,LB2B)))); | 2 | 0.03% |
| TREE 622 | 0.000284535 | (Lbru,(LB1A,((LB2A,(LB3B,LB2B)),(LB1B,LB3A)))); | 2 | 0.03% |
| TREE 623 | 0.000284535 | (Lbru,(LB2A,(LB2B,(LB3B,(LB1B,(LB1A,LB3A)))))); | 2 | 0.03% |
| TREE 624 | 0.000284535 | (Lbru,(LB3A,(LB1A,(LB2A,(LB3B,(LB1B,LB2B)))))); | 2 | 0.03% |
| TREE 625 | 0.000284535 | (Lbru,(LB2B,(LB3B,(LB2A,(LB1B,(LB1A,LB3A)))))); | 2 | 0.03% |
| TREE 626 | 0.000284535 | (Lbru,(LB2A,((LB3B,LB2B),(LB1B,(LB1A,LB3A))))); | 2 | 0.03% |
| TREE 627 | 0.000284535 | (Lbru,((LB1B,LB3B),(LB2B,(LB3A,(LB1A,LB2A))))); | 2 | 0.03% |
| TREE 628 | 0.000284535 | (Lbru,((LB2A,LB2B),(LB3B,(LB1A,(LB1B,LB3A))))); | 2 | 0.03% |
| TREE 629 | 0.000284535 | (Lbru,((LB1A,LB3B),((LB1B,LB2A),(LB3A,LB2B)))); | 2 | 0.03% |
| TREE 630 | 0.000284535 | (Lbru,((LB1A,LB2B),((LB3B,LB2A),(LB1B,LB3A)))); | 2 | 0.03% |
| TREE 631 | 0.000284535 | (Lbru,(LB1A,(LB3B,(LB1B,(LB2B,(LB3A,LB2A)))))); | 2 | 0.03% |
| TREE 632 | 0.000284535 | (Lbru,(LB3B,((LB1B,(LB1A,LB2A)),(LB3A,LB2B)))); | 2 | 0.03% |
| TREE 633 | 0.000284535 | (Lbru,(LB1A,(LB3A,(LB2B,(LB2A,(LB1B,LB3B)))))); | 2 | 0.03% |
| TREE 634 | 0.000284535 | (Lbru,((LB3B,(LB2A,LB2B)),(LB1A,(LB1B,LB3A)))); | 2 | 0.03% |
| TREE 635 | 0.000284535 | (Lbru,(LB2B,(LB3B,(LB1A,(LB1B,(LB3A,LB2A)))))); | 2 | 0.03% |
| TREE 636 | 0.000284535 | (Lbru,((LB1A,LB2B),((LB1B,LB3B),(LB3A,LB2A)))); | 2 | 0.03% |
| TREE 637 | 0.000284535 | (Lbru,((LB1B,LB2B),(LB3B,(LB2A,(LB1A,LB3A))))); | 2 | 0.03% |
| TREE 638 | 0.000284535 | (Lbru,(LB3A,((LB1A,LB2B),(LB2A,(LB1B,LB3B))))); | 2 | 0.03% |
| TREE 639 | 0.000284535 | (Lbru,(LB3A,(LB1A,((LB1B,LB2A),(LB3B,LB2B))))); | 2 | 0.03% |
| TREE 640 | 0.000284535 | (Lbru,((LB2A,(LB1B,LB3B)),(LB2B,(LB1A,LB3A)))); | 2 | 0.03% |
| TREE 641 | 0.000284535 | (Lbru,(LB3A,(LB1A,((LB1B,LB2B),(LB3B,LB2A))))); | 2 | 0.03% |
| TREE 642 | 0.000284535 | (Lbru,(LB3B,(LB2A,(LB1B,(LB2B,(LB1A,LB3A)))))); | 2 | 0.03% |
| TREE 643 | 0.000284535 | (Lbru,((LB1B,LB2A),((LB1A,LB2B),(LB3B,LB3A)))); | 2 | 0.03% |
| TREE 644 | 0.000284535 | (Lbru,(LB2B,((LB3B,(LB1B,LB2A)),(LB1A,LB3A)))); | 2 | 0.03% |
| TREE 645 | 0.000284535 | (Lbru,(LB3B,((LB1B,(LB2A,LB2B)),(LB1A,LB3A)))); | 2 | 0.03% |
| TREE 646 | 0.000284535 | (Lbru,((LB1B,(LB2B,(LB3B,LB2A))),(LB1A,LB3A))); | 2 | 0.03% |
| TREE 647 | 0.000284535 | (Lbru,((LB1A,LB2B),(LB1B,(LB3B,(LB3A,LB2A))))); | 2 | 0.03% |
| TREE 648 | 0.000284535 | (Lbru,(LB1A,(LB2B,((LB3B,LB2A),(LB1B,LB3A))))); | 2 | 0.03% |
| TREE 649 | 0.000284535 | (Lbru,((LB1A,(LB3B,LB2A)),(LB1B,(LB3A,LB2B)))); | 2 | 0.03% |
| TREE 650 | 0.000284535 | (Lbru,((LB1A,(LB3B,(LB2A,LB2B))),(LB1B,LB3A))); | 2 | 0.03% |
| TREE 651 | 0.000142268 | (Lbru,(LB3A,((LB1A,LB2A),(LB3B,(LB1B,LB2B))))); | 1 | 0.01% |
| TREE 652 | 0.000142268 | (Lbru,((LB3B,LB2B),(LB3A,(LB1A,(LB1B,LB2A))))); | 1 | 0.01% |
| TREE 653 | 0.000142268 | (Lbru,(LB3A,(LB2A,(LB1B,(LB3B,(LB1A,LB2B)))))); | 1 | 0.01% |
| TREE 654 | 0.000142268 | (Lbru,(LB2B,(LB3B,(LB1B,(LB3A,(LB1A,LB2A)))))); | 1 | 0.01% |
| TREE 655 | 0.000142268 | (Lbru,((LB1A,(LB2B,(LB1B,LB3B))),(LB3A,LB2A))); | 1 | 0.01% |
| TREE 656 | 0.000142268 | (Lbru,(LB3B,((LB1B,LB2B),(LB3A,(LB1A,LB2A))))); | 1 | 0.01% |
| TREE 657 | 0.000142268 | (Lbru,(LB3A,((LB2A,LB2B),(LB1B,(LB1A,LB3B))))); | 1 | 0.01% |
| TREE 658 | 0.000142268 | (Lbru,((LB2B,(LB1B,LB2A)),(LB3B,(LB1A,LB3A)))); | 1 | 0.01% |
| TREE 659 | 0.000142268 | (Lbru,(LB3A,(LB2B,(LB2A,(LB1A,(LB1B,LB3B)))))); | 1 | 0.01% |
| TREE 660 | 0.000142268 | (Lbru,((LB1B,(LB1A,(LB3B,LB2A))),(LB3A,LB2B))); | 1 | 0.01% |
| TREE 661 | 0.000142268 | (Lbru,(LB2B,(LB1B,(LB3B,(LB1A,(LB3A,LB2A)))))); | 1 | 0.01% |
| TREE 662 | 0.000142268 | (Lbru,(LB2B,((LB1A,LB3B),(LB3A,(LB1B,LB2A))))); | 1 | 0.01% |
| TREE 663 | 0.000142268 | (Lbru,(LB2A,(LB3B,(LB1A,(LB2B,(LB1B,LB3A)))))); | 1 | 0.01% |
| TREE 664 | 0.000142268 | (Lbru,((LB3B,LB2A),(LB2B,(LB1A,(LB1B,LB3A))))); | 1 | 0.01% |
| TREE 665 | 0.000142268 | (Lbru,((LB3B,LB2A),((LB1A,LB2B),(LB1B,LB3A)))); | 1 | 0.01% |
| TREE 666 | 0.000142268 | (Lbru,(LB2A,((LB1A,LB3B),(LB3A,(LB1B,LB2B))))); | 1 | 0.01% |
| TREE 667 | 0.000142268 | (Lbru,(LB3B,(LB3A,(LB2B,(LB1A,(LB1B,LB2A)))))); | 1 | 0.01% |
| TREE 668 | 0.000142268 | (Lbru,(LB3B,(LB1A,(LB2B,(LB2A,(LB1B,LB3A)))))); | 1 | 0.01% |
| TREE 669 | 0.000142268 | (Lbru,(LB1A,(LB3A,(LB2B,(LB1B,(LB3B,LB2A)))))); | 1 | 0.01% |
| TREE 670 | 0.000142268 | (Lbru,(LB1A,(LB3A,(LB1B,(LB2A,(LB3B,LB2B)))))); | 1 | 0.01% |
| TREE 671 | 0.000142268 | (Lbru,(LB2A,(LB1B,(LB3B,(LB1A,(LB3A,LB2B)))))); | 1 | 0.01% |
| TREE 672 | 0.000142268 | (Lbru,((LB1B,LB3B),(LB2B,(LB2A,(LB1A,LB3A))))); | 1 | 0.01% |
| TREE 673 | 0.000142268 | (Lbru,(LB1A,((LB3B,(LB1B,LB2A)),(LB3A,LB2B)))); | 1 | 0.01% |
| TREE 674 | 0.000142268 | (Lbru,(LB3A,(LB2A,(LB1A,(LB3B,(LB1B,LB2B)))))); | 1 | 0.01% |
| TREE 675 | 0.000142268 | (Lbru,((LB3B,(LB1B,LB2B)),(LB1A,(LB3A,LB2A)))); | 1 | 0.01% |
| TREE 676 | 0.000142268 | (Lbru,(LB2A,(LB1A,(LB3B,(LB1B,(LB3A,LB2B)))))); | 1 | 0.01% |
| TREE 677 | 0.000142268 | (Lbru,((LB1B,LB2A),(LB3A,(LB1A,(LB3B,LB2B))))); | 1 | 0.01% |
| TREE 678 | 0.000142268 | (Lbru,(LB2A,(LB3A,(LB2B,(LB3B,(LB1B,LB1A)))))); | 1 | 0.01% |
| TREE 679 | 0.000142268 | (Lbru,(LB3A,(LB1A,(LB1B,(LB2A,(LB3B,LB2B)))))); | 1 | 0.01% |
| TREE 680 | 0.000142268 | (Lbru,(LB3A,((LB1A,LB2B),(LB1B,(LB3B,LB2A))))); | 1 | 0.01% |
| TREE 681 | 0.000142268 | (Lbru,(LB3A,((LB1B,LB2B),(LB3B,(LB1A,LB2A))))); | 1 | 0.01% |
| TREE 682 | 0.000142268 | (Lbru,((LB1A,LB3B),((LB1B,LB2B),(LB3A,LB2A)))); | 1 | 0.01% |
| TREE 683 | 0.000142268 | (Lbru,((LB1B,LB3B),(LB1A,(LB3A,(LB2A,LB2B))))); | 1 | 0.01% |
| TREE 684 | 0.000142268 | (Lbru,((LB1B,LB3B),(LB2A,(LB2B,(LB1A,LB3A))))); | 1 | 0.01% |
| TREE 685 | 0.000142268 | (Lbru,((LB1A,(LB1B,LB3B)),(LB2A,(LB3A,LB2B)))); | 1 | 0.01% |
| TREE 686 | 0.000142268 | (Lbru,(LB2A,(LB3B,(LB1B,(LB2B,(LB1A,LB3A)))))); | 1 | 0.01% |
| TREE 687 | 0.000142268 | (Lbru,((LB1A,(LB1B,LB2B)),(LB3B,(LB3A,LB2A)))); | 1 | 0.01% |
| TREE 688 | 0.000142268 | (Lbru,(LB1B,((LB1A,(LB3B,LB2B)),(LB3A,LB2A)))); | 1 | 0.01% |
| TREE 689 | 0.000142268 | (Lbru,(LB3A,((LB1B,LB2B),(LB2A,(LB1A,LB3B))))); | 1 | 0.01% |
| TREE 690 | 0.000142268 | (Lbru,((LB2A,(LB1B,LB3B)),(LB3A,(LB1A,LB2B)))); | 1 | 0.01% |
| TREE 691 | 0.000142268 | (Lbru,(LB2B,(LB3A,(LB1A,(LB2A,(LB1B,LB3B)))))); | 1 | 0.01% |
| TREE 692 | 0.000142268 | (Lbru,(LB3A,((LB2B,(LB1A,LB2A)),(LB1B,LB3B)))); | 1 | 0.01% |
| TREE 693 | 0.000142268 | (Lbru,(LB2B,((LB1A,LB3B),(LB2A,(LB1B,LB3A))))); | 1 | 0.01% |
| TREE 694 | 0.000142268 | (Lbru,(LB1B,(LB3B,((LB2A,LB2B),(LB1A,LB3A))))); | 1 | 0.01% |
| TREE 695 | 0.000142268 | (Lbru,(((LB1A,LB2B),(LB3B,LB2A)),(LB1B,LB3A))); | 1 | 0.01% |
| TREE 696 | 0.000142268 | (Lbru,(LB2B,(LB3B,(LB1A,(LB2A,(LB1B,LB3A)))))); | 1 | 0.01% |
| TREE 697 | 0.000142268 | (Lbru,(LB2B,((LB1A,(LB1B,LB3B)),(LB3A,LB2A)))); | 1 | 0.01% |
| TREE 698 | 0.000142268 | (Lbru,((LB1A,LB2A),(LB3A,(LB1B,(LB3B,LB2B))))); | 1 | 0.01% |
| TREE 699 | 0.000142268 | (Lbru,((LB1B,LB3B),(LB3A,(LB1A,(LB2A,LB2B))))); | 1 | 0.01% |
| TREE 700 | 0.000142268 | (Lbru,(LB1A,(LB3B,((LB1B,LB2B),(LB3A,LB2A))))); | 1 | 0.01% |
| TREE 701 | 0.000142268 | (Lbru,((LB3B,LB2A),((LB1B,LB2B),(LB1A,LB3A)))); | 1 | 0.01% |
| TREE 702 | 0.000142268 | (Lbru,(LB2A,(LB3B,(LB1A,(LB1B,(LB3A,LB2B)))))); | 1 | 0.01% |
| TREE 703 | 0.000142268 | (Lbru,(LB2B,(LB1B,(LB3A,(LB1A,(LB3B,LB2A)))))); | 1 | 0.01% |
| TREE 704 | 0.000142268 | (Lbru,(LB3A,(LB2A,(LB1B,(LB2B,(LB1A,LB3B)))))); | 1 | 0.01% |
| TREE 705 | 0.000142268 | (Lbru,(LB2A,(LB3A,(LB1B,(LB1A,(LB3B,LB2B)))))); | 1 | 0.01% |
| TREE 706 | 0.000142268 | (Lbru,(LB3B,(LB2B,((LB1B,LB2A),(LB1A,LB3A))))); | 1 | 0.01% |
| TREE 707 | 0.000142268 | (Lbru,((LB2A,(LB2B,(LB1A,LB3B))),(LB1B,LB3A))); | 1 | 0.01% |
| TREE 708 | 0.000142268 | (Lbru,((LB1A,LB3B),(LB2B,(LB2A,(LB1B,LB3A))))); | 1 | 0.01% |
| TREE 709 | 0.000142268 | (Lbru,((LB1A,LB2B),(LB3A,(LB3B,(LB1B,LB2A))))); | 1 | 0.01% |
| TREE 710 | 0.000142268 | (Lbru,(LB3A,(LB1A,(LB3B,(LB2A,(LB1B,LB2B)))))); | 1 | 0.01% |
| TREE 711 | 0.000142268 | (Lbru,(LB2A,(LB2B,(LB3B,(LB1A,(LB1B,LB3A)))))); | 1 | 0.01% |
| TREE 712 | 0.000142268 | (Lbru,(LB3B,(LB2A,(LB1A,(LB3A,(LB1B,LB2B)))))); | 1 | 0.01% |
| TREE 713 | 0.000142268 | (Lbru,((LB1B,(LB3B,(LB1A,LB2A))),(LB3A,LB2B))); | 1 | 0.01% |
| TREE 714 | 0.000142268 | (Lbru,((LB1A,(LB2A,(LB1B,LB3B))),(LB3A,LB2B))); | 1 | 0.01% |
| TREE 715 | 0.000142268 | (Lbru,(LB2B,(LB3B,((LB1B,LB2A),(LB1A,LB3A))))); | 1 | 0.01% |
| TREE 716 | 0.000142268 | (Lbru,(LB2A,(LB3A,(LB1A,(LB2B,(LB1B,LB3B)))))); | 1 | 0.01% |
| TREE 717 | 0.000142268 | (Lbru,(LB3B,((LB1A,(LB2A,LB2B)),(LB1B,LB3A)))); | 1 | 0.01% |
| TREE 718 | 0.000142268 | (Lbru,(LB1B,(LB3B,(LB2B,(LB3A,(LB1A,LB2A)))))); | 1 | 0.01% |
| TREE 719 | 0.000142268 | (Lbru,(LB1A,((LB1B,LB3B),(LB2A,(LB3A,LB2B))))); | 1 | 0.01% |
| TREE 720 | 0.000142268 | (Lbru,(LB3A,(LB1A,(LB2B,(LB2A,(LB1B,LB3B)))))); | 1 | 0.01% |
| TREE 721 | 0.000142268 | (Lbru,((LB1A,LB3B),(LB2B,(LB3A,(LB1B,LB2A))))); | 1 | 0.01% |
| TREE 722 | 0.000142268 | (Lbru,(LB1B,(LB3A,(LB2A,(LB1A,(LB3B,LB2B)))))); | 1 | 0.01% |
| TREE 723 | 0.000142268 | (Lbru,(LB3B,(LB2A,(LB1A,(LB2B,(LB1B,LB3A)))))); | 1 | 0.01% |
| TREE 724 | 0.000142268 | (Lbru,(LB3A,((LB1A,LB2A),(LB2B,(LB1B,LB3B))))); | 1 | 0.01% |
| TREE 725 | 0.000142268 | (Lbru,(LB2A,(LB3A,(LB2B,(LB1A,(LB1B,LB3B)))))); | 1 | 0.01% |
| TREE 726 | 0.000142268 | (Lbru,(LB1A,(LB3B,((LB1B,LB2A),(LB3A,LB2B))))); | 1 | 0.01% |
| TREE 727 | 0.000142268 | (Lbru,(LB3A,(LB3B,(LB1A,(LB2B,(LB1B,LB2A)))))); | 1 | 0.01% |
| TREE 728 | 0.000142268 | (Lbru,(LB3A,(LB1A,(LB3B,(LB2B,(LB1B,LB2A)))))); | 1 | 0.01% |
| TREE 729 | 0.000142268 | (Lbru,(LB1A,(LB3A,(LB2B,(LB3B,(LB1B,LB2A)))))); | 1 | 0.01% |
| TREE 730 | 0.000142268 | (Lbru,(LB3B,(LB1A,(LB3A,(LB2B,(LB1B,LB2A)))))); | 1 | 0.01% |
| TREE 731 | 0.000142268 | (Lbru,(((LB1B,LB2B),(LB1A,LB3B)),(LB3A,LB2A))); | 1 | 0.01% |
| TREE 732 | 0.000142268 | (Lbru,(LB2B,(LB1B,((LB1A,LB3B),(LB3A,LB2A))))); | 1 | 0.01% |
| TREE 733 | 0.000142268 | (Lbru,((LB3B,LB2A),(LB1B,(LB3A,(LB1A,LB2B))))); | 1 | 0.01% |
| TREE 734 | 0.000142268 | (Lbru,(LB3B,(LB2B,(LB1A,(LB1B,(LB3A,LB2A)))))); | 1 | 0.01% |
| TREE 735 | 0.000142268 | (Lbru,((LB3B,LB2B),((LB1A,LB2A),(LB1B,LB3A)))); | 1 | 0.01% |
| TREE 736 | 0.000142268 | (Lbru,(LB1B,(LB2A,(LB3A,(LB1A,(LB3B,LB2B)))))); | 1 | 0.01% |
| TREE 737 | 0.000142268 | (Lbru,((LB1B,LB2B),(LB3A,(LB3B,(LB1A,LB2A))))); | 1 | 0.01% |
| TREE 738 | 0.000142268 | (Lbru,(LB2B,((LB1B,LB3B),(LB3A,(LB1A,LB2A))))); | 1 | 0.01% |
| TREE 739 | 0.000142268 | (Lbru,(LB3A,(LB1B,((LB1A,LB2B),(LB3B,LB2A))))); | 1 | 0.01% |
| TREE 740 | 0.000142268 | (Lbru,(LB1B,(LB3B,((LB1A,LB2B),(LB3A,LB2A))))); | 1 | 0.01% |
| TREE 741 | 0.000142268 | (Lbru,((LB1B,LB2A),((LB1A,LB3B),(LB3A,LB2B)))); | 1 | 0.01% |
| TREE 742 | 0.000142268 | (Lbru,((LB1B,(LB3B,(LB1A,LB2B))),(LB3A,LB2A))); | 1 | 0.01% |
| TREE 743 | 0.000142268 | (Lbru,(LB1A,((LB2A,(LB1B,LB3B)),(LB3A,LB2B)))); | 1 | 0.01% |
| TREE 744 | 0.000142268 | (Lbru,(LB3B,(LB1B,((LB2A,LB2B),(LB1A,LB3A))))); | 1 | 0.01% |
| TREE 745 | 0.000142268 | (Lbru,((LB2B,(LB1A,LB2A)),(LB3A,(LB1B,LB3B)))); | 1 | 0.01% |
| TREE 746 | 0.000142268 | (Lbru,(LB3A,(LB2B,(LB1A,(LB2A,(LB1B,LB3B)))))); | 1 | 0.01% |
| TREE 747 | 0.000142268 | (Lbru,((LB3B,(LB1B,(LB1A,LB2A))),(LB3A,LB2B))); | 1 | 0.01% |
| TREE 748 | 0.000142268 | (Lbru,((LB2A,(LB1A,(LB1B,LB3B))),(LB3A,LB2B))); | 1 | 0.01% |
| TREE 749 | 0.000142268 | (Lbru,((LB2B,(LB1B,LB3B)),(LB3A,(LB1A,LB2A)))); | 1 | 0.01% |
| TREE 750 | 0.000142268 | (Lbru,(LB1A,(LB3B,(LB2B,(LB3A,(LB1B,LB2A)))))); | 1 | 0.01% |
| TREE 751 | 0.000142268 | (Lbru,(LB2B,(LB1A,(LB3A,(LB3B,(LB1B,LB2A)))))); | 1 | 0.01% |
| TREE 752 | 0.000142268 | (Lbru,(LB1A,(LB3A,((LB2A,LB2B),(LB1B,LB3B))))); | 1 | 0.01% |
| TREE 753 | 0.000142268 | (Lbru,((LB2A,(LB1B,LB3B)),(LB1A,(LB3A,LB2B)))); | 1 | 0.01% |
| TREE 754 | 0.000142268 | (Lbru,(LB3B,(LB3A,((LB1B,LB2B),(LB1A,LB2A))))); | 1 | 0.01% |
| TREE 755 | 0.000142268 | (Lbru,((LB1A,(LB2A,(LB3B,LB2B))),(LB1B,LB3A))); | 1 | 0.01% |
| TREE 756 | 0.000142268 | (Lbru,(LB1B,((LB3B,LB2A),(LB3A,(LB1A,LB2B))))); | 1 | 0.01% |
| TREE 757 | 0.000142268 | (Lbru,(LB1A,((LB3B,LB2B),(LB3A,(LB1B,LB2A))))); | 1 | 0.01% |
| TREE 758 | 0.000142268 | (Lbru,((LB1B,LB3B),(LB2B,(LB1A,(LB3A,LB2A))))); | 1 | 0.01% |
| TREE 759 | 0.000142268 | (Lbru,((LB1B,LB2B),(LB3B,(LB3A,(LB1A,LB2A))))); | 1 | 0.01% |
| TREE 760 | 0.000142268 | (Lbru,(LB3A,(LB1A,(LB2B,(LB1B,(LB3B,LB2A)))))); | 1 | 0.01% |
| TREE 761 | 0.000142268 | (Lbru,(LB1A,(LB3A,(LB2A,(LB3B,(LB1B,LB2B)))))); | 1 | 0.01% |
| TREE 762 | 0.000142268 | (Lbru,((LB2A,(LB1A,(LB3B,LB2B))),(LB1B,LB3A))); | 1 | 0.01% |
| TREE 763 | 0.000142268 | (Lbru,(LB1B,((LB2A,(LB1A,LB3B)),(LB3A,LB2B)))); | 1 | 0.01% |
| TREE 764 | 0.000142268 | (Lbru,(LB3A,((LB2A,(LB1A,LB2B)),(LB1B,LB3B)))); | 1 | 0.01% |
| TREE 765 | 0.000142268 | (Lbru,(LB2B,(LB1B,((LB3B,LB2A),(LB1A,LB3A))))); | 1 | 0.01% |
| TREE 766 | 0.000142268 | (Lbru,(LB3B,((LB1A,LB2B),(LB2A,(LB1B,LB3A))))); | 1 | 0.01% |
| TREE 767 | 0.000142268 | (Lbru,(LB2B,((LB1B,LB3B),(LB2A,(LB1A,LB3A))))); | 1 | 0.01% |
| TREE 768 | 0.000142268 | (Lbru,(LB3B,((LB1A,LB2B),(LB1B,(LB3A,LB2A))))); | 1 | 0.01% |
| TREE 769 | 0.000142268 | (Lbru,(LB1B,(LB2B,(LB3B,(LB2A,(LB1A,LB3A)))))); | 1 | 0.01% |
| TREE 770 | 0.000142268 | (Lbru,(LB2B,(LB3B,(LB1B,(LB1A,(LB3A,LB2A)))))); | 1 | 0.01% |
| TREE 771 | 0.000142268 | (Lbru,(LB3A,(LB2B,(LB2A,(LB1B,(LB1A,LB3B)))))); | 1 | 0.01% |
| TREE 772 | 0.000142268 | (Lbru,(LB3B,(LB2B,(LB1B,(LB3A,(LB1A,LB2A)))))); | 1 | 0.01% |
| TREE 773 | 0.000142268 | (Lbru,(LB2A,(LB3B,((LB1A,LB2B),(LB1B,LB3A))))); | 1 | 0.01% |
| TREE 774 | 0.000142268 | (Lbru,((LB1A,LB2B),(LB3A,(LB1B,(LB3B,LB2A))))); | 1 | 0.01% |
| TREE 775 | 0.000142268 | (Lbru,((LB1B,LB3B),(LB2A,(LB1A,(LB3A,LB2B))))); | 1 | 0.01% |
| TREE 776 | 0.000142268 | (Lbru,((LB3B,(LB1B,LB2B)),(LB2A,(LB1A,LB3A)))); | 1 | 0.01% |
| TREE 777 | 0.000142268 | (Lbru,(LB1B,(LB3A,(LB1A,(LB2A,(LB3B,LB2B)))))); | 1 | 0.01% |
| TREE 778 | 0.000142268 | (Lbru,(LB1A,((LB3B,(LB1B,LB2B)),(LB3A,LB2A)))); | 1 | 0.01% |
| TREE 779 | 0.000142268 | (Lbru,((LB3B,LB2B),(LB1B,(LB2A,(LB1A,LB3A))))); | 1 | 0.01% |
| TREE 780 | 0.000142268 | (Lbru,((LB1B,(LB3B,LB2A)),(LB3A,(LB1A,LB2B)))); | 1 | 0.01% |
| TREE 781 | 0.000142268 | (Lbru,(LB3B,(LB2B,(LB1B,(LB2A,(LB1A,LB3A)))))); | 1 | 0.01% |
| TREE 782 | 0.000142268 | (Lbru,((LB1A,LB3B),(LB3A,(LB2B,(LB1B,LB2A))))); | 1 | 0.01% |
| TREE 783 | 0.000142268 | (Lbru,(LB1A,((LB1B,(LB3B,LB2A)),(LB3A,LB2B)))); | 1 | 0.01% |
| TREE 784 | 0.000142268 | (Lbru,(LB3B,(LB2A,(LB3A,(LB1A,(LB1B,LB2B)))))); | 1 | 0.01% |
| TREE 785 | 0.000142268 | (Lbru,(LB3A,(LB1B,((LB1A,LB2A),(LB3B,LB2B))))); | 1 | 0.01% |
| TREE 786 | 0.000142268 | (Lbru,(((LB1B,LB2A),(LB3B,LB2B)),(LB1A,LB3A))); | 1 | 0.01% |
| TREE 787 | 0.000142268 | (Lbru,(LB2A,(LB3A,(LB1A,(LB3B,(LB1B,LB2B)))))); | 1 | 0.01% |
| TREE 788 | 0.000142268 | (Lbru,(LB2B,(LB3A,(LB3B,(LB1A,(LB1B,LB2A)))))); | 1 | 0.01% |
| TREE 789 | 0.000142268 | (Lbru,((LB3B,(LB2B,(LB1A,LB2A))),(LB1B,LB3A))); | 1 | 0.01% |
| TREE 790 | 0.000142268 | (Lbru,(LB1B,(LB3A,(LB1A,(LB2B,(LB3B,LB2A)))))); | 1 | 0.01% |
| TREE 791 | 0.000142268 | (Lbru,((LB1A,LB3B),(LB3A,(LB2A,(LB1B,LB2B))))); | 1 | 0.01% |
| TREE 792 | 0.000142268 | (Lbru,(LB1A,(LB3B,(LB2B,(LB1B,(LB3A,LB2A)))))); | 1 | 0.01% |
| TREE 793 | 0.000142268 | (Lbru,((LB2A,(LB1B,(LB1A,LB3B))),(LB3A,LB2B))); | 1 | 0.01% |
| TREE 794 | 0.000142268 | (Lbru,(LB2A,(LB3B,(LB2B,(LB1A,(LB1B,LB3A)))))); | 1 | 0.01% |
| TREE 795 | 0.000142268 | (Lbru,((LB3B,LB2B),(LB1B,(LB3A,(LB1A,LB2A))))); | 1 | 0.01% |
| TREE 796 | 0.000142268 | (Lbru,((LB1B,LB3B),(LB3A,(LB2A,(LB1A,LB2B))))); | 1 | 0.01% |
| TREE 797 | 0.000142268 | (Lbru,((LB1A,LB3B),(LB3A,(LB1B,(LB2A,LB2B))))); | 1 | 0.01% |
| TREE 798 | 0.000142268 | (Lbru,(LB2A,((LB1B,LB3B),(LB3A,(LB1A,LB2B))))); | 1 | 0.01% |
| TREE 799 | 0.000142268 | (Lbru,((LB1A,LB2A),((LB3B,LB2B),(LB1B,LB3A)))); | 1 | 0.01% |
| TREE 800 | 0.000142268 | (Lbru,(LB1B,(LB2A,((LB3B,LB2B),(LB1A,LB3A))))); | 1 | 0.01% |
| TREE 801 | 0.000142268 | (Lbru,(LB2B,(LB3A,((LB1A,LB2A),(LB1B,LB3B))))); | 1 | 0.01% |
| TREE 802 | 0.000142268 | (Lbru,(LB3A,(LB2B,(LB3B,(LB1A,(LB1B,LB2A)))))); | 1 | 0.01% |
| TREE 803 | 0.000142268 | (Lbru,(LB2A,(LB3B,(LB1A,(LB3A,(LB1B,LB2B)))))); | 1 | 0.01% |
| TREE 804 | 0.000142268 | (Lbru,(LB3B,(LB1B,(LB2A,(LB2B,(LB1A,LB3A)))))); | 1 | 0.01% |
| TREE 805 | 0.000142268 | (Lbru,(LB2B,(LB2A,(LB3A,(LB1A,(LB1B,LB3B)))))); | 1 | 0.01% |
| TREE 806 | 0.000142268 | (Lbru,(LB1B,(LB2A,((LB1A,LB3B),(LB3A,LB2B))))); | 1 | 0.01% |
| TREE 807 | 0.000142268 | (Lbru,(LB3B,(LB1B,(LB2B,(LB2A,(LB1A,LB3A)))))); | 1 | 0.01% |
| TREE 808 | 0.000142268 | (Lbru,(LB1B,(LB3A,(LB2B,(LB2A,(LB1A,LB3B)))))); | 1 | 0.01% |
| TREE 809 | 0.000142268 | (Lbru,(LB2B,(LB1A,((LB1B,LB3B),(LB3A,LB2A))))); | 1 | 0.01% |
| TREE 810 | 0.000142268 | (Lbru,((LB1A,LB3B),(LB2A,(LB3A,(LB1B,LB2B))))); | 1 | 0.01% |
| TREE 811 | 0.000142268 | (Lbru,(LB3B,(LB2B,(LB1B,(LB1A,(LB3A,LB2A)))))); | 1 | 0.01% |
| TREE 812 | 0.000142268 | (Lbru,(LB3A,(LB1A,(LB2B,(LB3B,(LB1B,LB2A)))))); | 1 | 0.01% |
| TREE 813 | 0.000142268 | (Lbru,((LB1A,(LB3B,LB2A)),(LB3A,(LB1B,LB2B)))); | 1 | 0.01% |
| TREE 814 | 0.000142268 | (Lbru,((LB3B,(LB1B,LB2A)),(LB3A,(LB1A,LB2B)))); | 1 | 0.01% |
| TREE 815 | 0.000142268 | (Lbru,(LB2A,(LB3A,(LB3B,(LB1A,(LB1B,LB2B)))))); | 1 | 0.01% |
| TREE 816 | 0.000142268 | (Lbru,((LB1B,(LB3B,LB2A)),(LB1A,(LB3A,LB2B)))); | 1 | 0.01% |
| TREE 817 | 0.000142268 | (Lbru,(LB2A,(LB2B,((LB1B,LB3B),(LB1A,LB3A))))); | 1 | 0.01% |
| TREE 818 | 0.000142268 | (Lbru,(LB2A,(LB3A,(LB1A,(LB1B,(LB3B,LB2B)))))); | 1 | 0.01% |
| TREE 819 | 0.000142268 | (Lbru,(LB3B,(LB2B,(LB2A,(LB1A,(LB1B,LB3A)))))); | 1 | 0.01% |
| TREE 820 | 0.000142268 | (Lbru,((LB1B,LB3B),(LB1A,(LB2B,(LB3A,LB2A))))); | 1 | 0.01% |
| TREE 821 | 0.000142268 | (Lbru,(LB2A,(LB3B,((LB1B,LB2B),(LB1A,LB3A))))); | 1 | 0.01% |
| TREE 822 | 0.000142268 | (Lbru,(LB1A,(LB2B,(LB3A,(LB1B,(LB3B,LB2A)))))); | 1 | 0.01% |
| TREE 823 | 0.000142268 | (Lbru,((LB1A,(LB1B,(LB3B,LB2B))),(LB3A,LB2A))); | 1 | 0.01% |
| TREE 824 | 0.000142268 | (Lbru,(LB1A,(LB3B,(LB2A,(LB2B,(LB1B,LB3A)))))); | 1 | 0.01% |

**Table S6 Repeat region composition of three strains of *Liposcelis bostrychophila* and *Lipocselis brunnea***

|  | Lb_1 | Lb_2 | Lb_3 | Lbru |
| --- | --- | --- | --- | --- |
| Non RR | 0.635141915 | 0.650008542 | 0.633548051 | 0.841696457 |
| SINEs | 0.002469071 | 0.002438722 | 0.00274263 | 0.000174713 |
| LINEs | 0.021027192 | 0.01785206 | 0.019935219 | 0.025532506 |
| LTR | 0.025411032 | 0.018261848 | 0.021309196 | 0.011929577 |
| DNA transposons | 0.071046359 | 0.065389458 | 0.063178083 | 0.021594295 |
| Tandem repeats | 0.033960367 | 0.032297632 | 0.034076836 | 0.013382767 |
| Rolling-circles | 0.001513689 | 0.001689241 | 0.001948439 | 0.007198221 |
| Small_RNA | 0.002733627 | 0.002172077 | 0.00263314 | 7.16515E-05 |
| Unclassified RR | 0.206696748 | 0.209890419 | 0.220628405 | 0.078419813 |

**Table S7 Genes in each strain with dN/dS > 0.6.**

| **Chromosome** | **dN/dS > 0.6** | | |
| --- | --- | --- | --- |
|  | **Lb_1** | **Lb_2** | **Lb_3** |
| **chr1** | 173 | 238 | 346 |
| **chr2** | 186 | 273 | 443 |
| **chr3** | 146 | 193 | 247 |
| **chr4** | 311 | 443 | 712 |
| **chr5** | 172 | 281 | 362 |
| **chr6** | 132 | 167 | 241 |
| **chr7** | 134 | 184 | 248 |
| **chr8** | 130 | 172 | 266 |

**Table S8 Genes in each strain with dN/dS > 0.6 in *L. bostrychophila* strain 1.**

| Description | pvalue | p.adjust |
| --- | --- | --- |
| transcription cis-regulatory region binding | 2.50E-08 | 7.76E-05 |
| transcription regulatory region nucleic acid binding | 3.16E-08 | 7.76E-05 |
| DNA-binding transcription factor activity, RNA polymerase II-specific | 7.99E-08 | 0.000130813 |
| DNA-binding transcription factor activity, RNA polymerase II-specific | 1.06E-06 | 0.001261988 |
| transcription cis-regulatory region binding | 1.28E-06 | 0.001261988 |
| sequence-specific double-stranded DNA binding | 2.29E-06 | 0.001877955 |
| cis-regulatory region sequence-specific DNA binding | 2.76E-06 | 0.001933928 |
| RNA polymerase II transcription regulatory region sequence-specific DNA binding | 4.84E-06 | 0.002676184 |
| cis-regulatory region sequence-specific DNA binding | 4.95E-06 | 0.002676184 |
| DNA-binding transcription activator activity, RNA polymerase II-specific | 5.91E-06 | 0.002676184 |
| RNA polymerase II transcription regulatory region sequence-specific DNA binding | 5.99E-06 | 0.002676184 |
| regulation of reproductive process | 1.05E-05 | 0.004313989 |
| RNA polymerase II cis-regulatory region sequence-specific DNA binding | 1.59E-05 | 0.006027755 |
| DNA-binding transcription activator activity, RNA polymerase II-specific | 1.77E-05 | 0.006204172 |
| ventral spinal cord development | 2.09E-05 | 0.006748654 |
| cis-regulatory region sequence-specific DNA binding | 2.45E-05 | 0.006748654 |
| double-stranded DNA binding | 2.46E-05 | 0.006748654 |
| central nervous system neuron differentiation | 2.47E-05 | 0.006748654 |
| cell fate commitment | 3.39E-05 | 0.008011317 |
| DNA-binding transcription factor activity, RNA polymerase II-specific | 3.42E-05 | 0.008011317 |
| spinal cord motor neuron differentiation | 3.42E-05 | 0.008011317 |
| spinal cord development | 4.11E-05 | 0.009188481 |
| cell differentiation in spinal cord | 4.52E-05 | 0.009635105 |
| cell fate specification | 4.81E-05 | 0.009635105 |
| dorsal spinal cord development | 4.90E-05 | 0.009635105 |
| positive regulation of cell population proliferation | 5.65E-05 | 0.010672736 |
| neuron fate commitment | 6.01E-05 | 0.010928651 |
| limbic system development | 6.91E-05 | 0.012126886 |
| behavioral response to ethanol | 9.45E-05 | 0.015564717 |
| appendage morphogenesis | 9.50E-05 | 0.015564717 |
| regulation of cell fate commitment | 0.000104692 | 0.016592052 |
| learning or memory | 0.000114028 | 0.017291894 |
| forebrain development | 0.000116147 | 0.017291894 |
| regulation of animal organ morphogenesis | 0.000120577 | 0.017423401 |
| regulation of cell fate specification | 0.000125006 | 0.017547227 |
| modulation of chemical synaptic transmission | 0.000153898 | 0.021002817 |
| canonical Wnt signaling pathway | 0.000159224 | 0.021142346 |
| regulation of trans-synaptic signaling | 0.000177378 | 0.022933151 |
| appendage development | 0.000220148 | 0.027262761 |
| regulation of synaptic plasticity | 0.000228592 | 0.027262761 |
| sterol binding | 0.000238573 | 0.027262761 |
| cardiac ventricle development | 0.000239387 | 0.027262761 |
| cognition | 0.000241274 | 0.027262761 |
| regulation of system process | 0.000244377 | 0.027262761 |
| adult behavior | 0.00024971 | 0.027262761 |
| telencephalon development | 0.000274433 | 0.029310613 |
| cell fate determination | 0.000310112 | 0.032416619 |
| transcription coactivator binding | 0.000336971 | 0.032905587 |
| potassium channel regulator activity | 0.000336971 | 0.032905587 |
| germline ring canal | 0.000336971 | 0.032905587 |
| muscle organ development | 0.00034158 | 0.032905587 |
| axis specification | 0.000355672 | 0.033180866 |
| single-stranded RNA binding | 0.000360032 | 0.033180866 |
| eye morphogenesis | 0.000364699 | 0.033180866 |
| metamorphosis | 0.000375451 | 0.033538042 |
| photoreceptor cell differentiation | 0.000397182 | 0.034562145 |
| eye-antennal disc development | 0.000400986 | 0.034562145 |
| regulation of neuronal synaptic plasticity | 0.000445702 | 0.037754052 |
| post-embryonic animal organ development | 0.000490733 | 0.040863899 |
| cardiac ventricle morphogenesis | 0.000506717 | 0.041491651 |
| cell surface receptor signaling pathway involved in cell-cell signaling | 0.00052782 | 0.042511138 |
| nonassociative learning | 0.00054312 | 0.042701394 |
| molting cycle, chitin-based cuticle | 0.000547565 | 0.042701394 |
| molting cycle | 0.000560194 | 0.043003621 |
| instar larval or pupal morphogenesis | 0.000646442 | 0.048767386 |
| imaginal disc morphogenesis | 0.000656547 | 0.048767386 |
| ventricular septum development | 0.000665055 | 0.048767386 |
| regulation of pole plasm oskar mRNA localization | 0.000676051 | 0.048844699 |
| positive regulation of reproductive process | 0.000694639 | 0.049460293 |

**Table S9 Genes in each strain with dN/dS > 0.6 in *L. bostrychophila* strain 2.**

| ID | Description | pvalue | p.adjust |
| --- | --- | --- | --- |
| GO:0099537 | trans-synaptic signaling | 2.82E-07 | 0.00071526 |
| GO:0099536 | synaptic signaling | 3.31E-07 | 0.00071526 |
| GO:0007268 | chemical synaptic transmission | 5.39E-07 | 0.00071526 |
| GO:0098916 | anterograde trans-synaptic signaling | 5.39E-07 | 0.00071526 |
| GO:0003705 | DNA-binding transcription factor activity, RNA polymerase II-specific | 1.32E-06 | 0.001406117 |
| GO:0072178 | nephric duct morphogenesis | 9.70E-06 | 0.007098383 |
| GO:0061213 | positive regulation of mesonephros development | 1.07E-05 | 0.007098383 |
| GO:0061217 | regulation of mesonephros development | 1.07E-05 | 0.007098383 |
| GO:0048568 | embryonic organ development | 1.26E-05 | 0.007449756 |
| GO:0090184 | positive regulation of kidney development | 3.68E-05 | 0.019528532 |
| GO:0048562 | embryonic organ morphogenesis | 4.08E-05 | 0.019683093 |
| GO:0090189 | regulation of branching involved in ureteric bud morphogenesis | 5.47E-05 | 0.022360864 |
| GO:0090190 | positive regulation of branching involved in ureteric bud morphogenesis | 5.47E-05 | 0.022360864 |
| GO:0072176 | nephric duct development | 8.70E-05 | 0.032331418 |
| GO:0090183 | regulation of kidney development | 9.13E-05 | 0.032331418 |
| GO:0042659 | regulation of cell fate specification | 0.00012575 | 0.041741162 |
| GO:0048705 | skeletal system morphogenesis | 0.000134671 | 0.042072837 |

**Table S10 Genes in each strain with dN/dS > 0.6 in *L. bostrychophila* strain 3.**

| ID | Description | pvalue | p.adjust |
| --- | --- | --- | --- |
| GO:1901654 | response to ketone | 2.18E-07 | 0.001437797 |
| GO:0044391 | ribosomal subunit | 1.86E-05 | 0.042930131 |
| GO:0001077 | DNA-binding transcription activator activity, RNA polymerase II-specific | 1.95E-05 | 0.042930131 |

**Table S11 23 genes in GO:2000241 (regulation of reproductive process) with dN/dS > 0.6.**

| **Gene ID** | **dN/dS** |  | **Gene Name** | **Function** |
| --- | --- | --- | --- | --- |
| Lbbj_000160 | 0.76695 |  | EAF1 | RNA polymerase II transcription elongation factor |
| Lbbj_000571 | 1.22108 |  | MMP12 | metalloendopeptidase activity |
| Lbbj_000976 | 0.65547 |  | unc-63 | channel activity. It is involved in the biological process described with ion transport |
| Lbbj_000993 | 0.71021 |  | nAChRa3 | Acetylcholine-activated cation-selective channel activity. It is involved in the biological process described with ion transport |
| Lbbj_001330 | 1.52124 |  | KCNN2 | Calmodulin binding domain |
| Lbbj_003012 | 1.29762 |  | UNC119 | GMP-PDE, delta subunit |
| Lbbj_003101 | 0.61083 |  | ATXN2 | LsmAD |
| Lbbj_003575 | 0.6304 |  | TFDP1 | Transcription factor DP |
| Lbbj_003661 | 1.3 |  | SYT2 | transporter activity. It is involved in the biological process described with transport |
| Lbbj_010482 | 1.46032 |  | TCEB1 | Found in Skp1 protein family |
| Lbbj_011011 | 1.30012 |  | BTF3L4 | NAC domain |
| Lbbj_015604 | 1.31156 |  | UBE2B | Ubiquitin-conjugating enzyme |
| Lbbj_017032 | 0.61392 |  | ACVR1C | Protein tyrosine kinase |
| Lbbj_018107 | 1.29998 |  | RBM8A | Core component of the splicing-dependent multiprotein exon junction complex (EJC) deposited at splice junctions on mRNAs |
| Lbbj_018684 | 0.60469 |  | WNT5B | Ligand for members of the frizzled family of seven transmembrane receptors |
| Lbbj_018720 | 1.3 |  | ENSA | Protein phosphatase inhibitor that specifically inhibits protein phosphatase 2A (PP2A) during mitosis |
| Lbbj_026795 | 0.85279 |  | LASP1 | Nebulin repeat |
| Lbbj_026913 | 1.29984 |  | NFIB | Recognizes and binds the palindromic sequence 5'- TTGGCNNNNNGCCAA-3' present in viral and cellular promoters and in the origin of replication of adenovirus type 2. These proteins are individually capable of activating transcription and replication |
| Lbbj_027470 | 0.63693 |  | RAB6B | Rab subfamily of small GTPases |
| Lbbj_028148 | 0.83054 |  | STAU1 | Staufen C-terminal domain |
| Lbbj_028891 | 0.7416 |  | OXTR | 7 transmembrane receptor (rhodopsin family) |
| Lbbj_029589 | 0.66155 |  | AGO2 | Required for RNA-mediated gene silencing (RNAi) by the RNA-induced silencing complex (RISC) |
| Lbbj_032086 | 0.7073 |  | ADCY3 | Belongs to the adenylyl cyclase class-4 guanylyl cyclase family |
